# Topical Activatable Fluorescence Probes for Rapid Intraoperative Detection of Peritoneal Dissemination in High-Grade Serous Ovarian Carcinoma

**DOI:** 10.64898/2026.07.21.739701

**Authors:** Hanae Sekine, Kyohhei Fujita, Emiko Yoshida, Tasuku Ueno, Toru Komatsu, Takuo Hayashi, Daiki Ogishima, Yusen Sugimura, Yasuteru Urano, Yasuhisa Terao

## Abstract

**Background:** Epithelial ovarian cancer (EOC) is one of the most lethal gynecologic malignancies, largely because most patients are diagnosed at an advanced stage with peritoneal dissemination. High-grade serous carcinoma (HGSC), the most common and aggressive EOC subtype, requires complete cytoreductive surgery to improve the prognosis; however, minute disseminated lesions are often missed by conventional intraoperative inspection. Here, we aimed to develop fluorescence probes for rapid and sensitive intraoperative detection of HGSC peritoneal dissemination.

**Methods:** We screened a hydroxymethyl rhodamine green (HMRG)-based fluorescence probe library consisting of 381 protease- and aminopeptidase-reactive fluorescence probes using tumor and non-tumor specimens from patients with HGSC. The target enzyme of the hit probes was identified by means of enzyme assays, immunohistochemistry, and LC/MS analysis. Diagnostic utility was evaluated *ex vivo* using clinical specimens and *in vivo* using a peritoneal dissemination mouse model.

**Results:** Three probes—EK-HMRG, NA-HMRG and DA-HMRG—were selected as promising candidates for the detection of peritoneal dissemination in HGSC. Puromycin-sensitive aminopeptidase (PSA) was identified as a novel target enzyme of these probes. EK-HMRG, NA-HMRG and DA-HMRG rapidly detected peritoneal dissemination just a few millimeters in size with high sensitivity and specificity in clinical HGSC specimens and in a peritoneal dissemination mouse model after topical application.

**Conclusions:** The PSA-targeting topical fluorescence probes EK-HMRG, NA-HMRG and DA-HMRG are promising tools for real-time, highly sensitive intraoperative visualization of peritoneal dissemination in HGSC, and are promising candidates to improve complete resection rates.

## Introduction

Ovarian cancer remains one of the most lethal gynecologic malignancies, with approximately 70% of patients diagnosed with advanced disease involving peritoneal dissemination at the time of initial diagnosis.^(1–3)^ High-grade serous ovarian carcinoma (HGSOC), the most common subtype of epithelial ovarian cancer (EOC) accounting for approximately 75% of cases, is characterized by aggressive clinical behavior and a high recurrence rate, with more than 80% of patients diagnosed at an advanced stage.^(4)^ Standard treatment consists of cytoreductive surgery and platinum-based chemotherapy. Importantly, complete resection of macroscopic disease is strongly associated with improved long-term survival.^(5–8)^ Since residual disease may persist within macroscopically normal-appearing peritoneum despite apparently complete gross resection,^(9)^ there is an urgent need for intraoperative technologies that can reliably detect and help remove minute peritoneal lesions

Conventional preoperative imaging modalities, such as computed tomography (CT), magnetic resonance imaging (MRI), and positron emission tomography/computed tomography (PET/CT), are frequently used to evaluate tumor extent. However, these techniques have limited spatial resolution and insufficient sensitivity for detecting subcentimeter-size peritoneal lesions. CT and PET/CT generally fail to reliably detect lesions smaller than approximately 10 mm, and even MRI with diffusion-weighted sequences shows markedly reduced sensitivity for subcentimeter nodules, particularly in the peritoneal surface and mesentery where complex anatomical structures further obscure tumor visualization.^(10–15)^ Moreover, it is difficult to perform real-time intraoperative assessment of peritoneal dissemination with these modalities.

To address these challenges, intraoperative fluorescence imaging has emerged as a promising technique for real-time and sensitive tumor detection during surgery.^(16–19)^ Fluorescent agents used for this purpose include contrast fluorophores, such as indocyanine green (ICG), and molecularly targeted probes, which are broadly classified as “always-on” or “activatable.” ICG and the always-on FRα-targeted probe Cytalux (pafolacianine, OTL38) have been investigated for ovarian cancer detection.^(20)^ However, because they emit fluorescence continuously, ICG often produces background fluorescence, while Cytalux exhibits false-positive signals due to nonspecific accumulation in normal tissues, thereby limiting the reliable detection of minute cancer lesions.^(21–26)^ In contrast, activatable fluorescence probes targeting enzymatic activities generate fluorescence only after reacting with cancer-specific biomarker enzymes, enabling low-background imaging with a high tumor/normal fluorescence ratio (T/N). ^(27–31)^ Such probes are attractive for intraoperative cancer imaging due to the highly amplified fluorescence generated by enzyme-catalyzed turnover at lesion sites. We previously developed hydroxymethyl rhodamine green (HMRG) as a green scaffold fluorophore for enzyme-activatable probes.^(32,33)^ By conjugating HMRG to various substrates, we constructed a fluorescence probe library targeting aminopeptidases and proteases, consisting of 381 probes.^(34)^ These P2P1-HMRG probes, in which various amino acids are introduced at the P1 and P2 positions, exist in a non-fluorescent spirocyclized form but become highly fluorescent upon enzymatic cleavage of the substrate moiety.^(32,33)^ Screening of this library using clinical cancer specimens has identified effective cancer-imaging probe/biomarker enzyme pairs in various cancer types.^(34–37)^ The identified enzymes exhibited high activities, and the probes can sensitively detect cancer tissues in an intraoperative time scale. Thus, efficient probe/biomarker enzyme pairs also likely exist for HGSC.

In this study, we screened an HMRG-based fluorescence probe library to identify probe/biomarker enzyme pairs for detecting ovarian cancer peritoneal dissemination. We identified puromycin-sensitive aminopeptidase (PSA) as a novel activity-elevated biomarker enzyme and demonstrated that three hit PSA-targeting probes—EK-HMRG, NA-HMRG, and DA-HMRG—enable rapid and highly sensitive detection of peritoneal implants just a few millimeters in size in HGSC specimens upon topical application.

## Materials and Methods

### Clinical samples and study approval

A total of 22 specimens were collected from 20 patients with HGSC who underwent surgical resection between 2010 and 2025 at Juntendo University Hospital and Juntendo Nerima Hospital. All specimens were obtained immediately after surgery and included peritoneal dissemination lesions or omental metastases. Of the 20 patients, nine had received neoadjuvant chemotherapy (NAC) prior to surgery. Four samples were subjected to fluorescence imaging in a fresh condition (peritoneum 2, omentum 2) within a few hours after resection. Other specimens were temporarily preserved at −80 °C in a freezer and thawed on ice at room temperature for imaging (Supplementary Table S1). Written informed consent for the use of tissue samples and clinical data was obtained from all patients prior to inclusion in the study. The study protocol was reviewed and approved by the Institutional Review Board of Juntendo University (approval number: E22-0474-H01) and was conducted in accordance with the Declaration of Helsinki.

### Enzyme-activatable fluorescence probes

The fluorescence probe library was prepared according to the literature.^(34)^ EK-HMRG, NA-HMRG and DA-HMRG were obtained from Goryokayaku Co., Tokyo, Japan.

### Preparation of homogenized lysates

Tissue specimens were placed into sample tubes containing zirconia beads (5-4060-13; AS ONE Corp.), and 500 µL of tissue protein extraction reagent (T-PER) (Thermo Fisher Scientific) was added. Homogenization was performed using a bead-type cell disruptor (Micro Smash MS-100; TOMY Digital Biology), followed by centrifugation at 10,000 × g for 5 min at 4°C. The supernatant (500 µL) was collected into a 2.0 mL plastic tube. This procedure was repeated once more with an additional 500 µL of T-PER, and the second supernatant was combined with the first. Protein concentrations in the lysates were measured using a bicinchoninic acid (BCA) protein assay kit (Thermo Fisher Scientific), and lysates were stored at –80°C until use.

### Primary screening

As a primary screening, we evaluated the reactivity of fluorescence probes using tissue lysates prepared from peritoneal dissemination lesions and adjacent non-tumorous peritoneum from five patients with HGSC. For each case, paired tumor and normal lysates were applied to 384-well black plates (6008269; PerkinElmer) at a final protein concentration of 0.05 mg/mL (5 µL per well). An equal volume (5 µL) of probe solution (final concentration: 0.9 µM, containing DMSO as a cosolvent) was added to each well. Fluorescence intensity was measured at 0, 1, 2, and 3 h after probe application using a plate reader (EnVision, PerkinElmer) equipped with a 485/14 nm excitation filter and a 535/25 nm emission filter. The probes were converted to the hydrolysis product HMRG upon reaction with their target enzymes in lysates, and the conversion rates (CVR) were calculated by dividing the fluorescence intensity of each probe by that of HMRG at the same measurement time.^(34)^ To identify tumor-selective probes, evaluation of the probes was performed based on both CVR [3 h–0 h] (CVR_3 h – 0 h_) and T/N.

### Secondary screening

*Ex vivo* imaging-based secondary screening was performed using small tissue fragments (approximately 2–3 mm in diameter) obtained from peritoneal dissemination lesions and adjacent normal peritoneum of HGSC patients. A 50 μM probe solution (200 μL) in PBS containing 0.5% v/v DMSO as a cosolvent was added to each well of an 8-well chamber (80826, μ-slide 8 well ibiTreat) containing a surgical specimen (tumor or normal tissue). Fluorescence images were acquired with the Maestro *in vivo* imaging system (PerkinElmer) before and at 0, 1, 3, 5, 10, 20, and 30 min after the probe application. The blue-filter setting (excitation, 445 to 490 nm; emission, 515 nm long-pass) was used. The tunable filter was automatically stepped in 10-nm increments from 500 to 800 nm while the camera sequentially captured images at each wavelength. Fluorescence images at 540 nm were extracted, and fluorescence intensities were quantified by drawing regions of interest (ROIs) with the Maestro software.^(38)^

### Diced Gel Electrophoresis and LC-MS/MS analysis

Protein concentrations of lysates prepared from peritoneal dissemination lesions of HGSC patients were adjusted to 3.0 mg/mL based on BCA assay results. Two-dimensional electrophoresis under non-denaturing conditions was performed by isoelectric focusing followed by native PAGE.^(39)^ The resulting gel was diced and transferred to a 384-well plate (SAINOME), centrifuged at 3,000 rpm for 5 min, and incubated with 70 µL of EK-HMRG solution (final concentration: 10 µM) at 37°C for 1 h. Fluorescence intensity was measured using an EnVision, and the data were visualized as a heat map. The gel slice corresponding to the well with the highest fluorescence was collected, washed three times with distilled water, and stored at −80°C. This procedure was repeated independently four times, and the collected gel fragments were analyzed by LC-MS/MS. Protein digestion and peptide identification were outsourced to a commercial service provider (APRO Life Science Institute, Inc.). LC-MS/MS data were analyzed using MASCOT Server 2.8.1 (Matrix Science Ltd.) with a significance threshold of *P* < 0.05. The search was performed against a database containing all known protein species. Since the list of identified proteins may include non-specific binders or irrelevant proteins, each hit was manually reviewed using literature sources and enzymatic function databases, including BRENDA (https://www.brenda-enzymes.org/), to determine whether it was a candidate for enzymatic cleavage of the probe. Proteins selected through this process were subjected to further biochemical validation.

### Fluorescence assay using purified enzymes

To evaluate the enzymatic activity of puromycin-sensitive aminopeptidase (PSA) toward each probe, 1 µM probe solutions (EK-HMRG, NA-HMRG, or DA-HMRG) in PBS containing 0.01% v/v DMSO as a cosolvent was incubated with purified PSA (2.5 ng per well; 6410-ZN-010; R&D Systems) in the presence or absence of puromycin (30 µM/L) at 37°C (n = 4). The assay was conducted in a total reaction volume of 20 µL. Fluorescence intensity was measured using an EnVision equipped with a 485/14 nm excitation filter and a 535/25 nm emission filter.

### Enzyme inhibition assay using tissue lysates

To confirm the enzymatic activity responsible for probe activation, tissue lysates prepared from ovarian cancer peritoneal dissemination lesions (final protein concentration: 0.1 mg/mL) were incubated with 1 µM probe solutions (EK-HMRG, NA-HMRG, or DA-HMRG) in PBS containing 0.01% v/v DMSO as a cosolvent in the presence or absence of inhibitor, puromycin (30 µM), at 37°C for 120 min (n = 4). The assay was conducted in 384-well plates (4511; Corning) with a total reaction volume of 20 µL (10 µL of lysate was added to 10 µL of probe solution per well). Fluorescence intensity was measured using an EnVision equipped with a 485/14 nm excitation filter and a 535/25 nm emission filter.

### Fluorescence imaging of tumor specimens with and without enzyme inhibitor

Fluorescence imaging of the tumor specimens was performed using EK-HMRG, NA-HMRG and DA-HMRG (50 µM) in the presence or absence of puromycin (final concentration 500 μM). Sample specimens were thawed at room temperature, and fluorescence imaging was performed under the same conditions as described in the screening methods.

### Immunohistochemical (IHC) analysis

Immunohistochemical staining was performed using the Ventana BenchMark GX automated staining system (Ventana Medical Systems). After deparaffinization, antigen retrieval was carried out by heat treatment at 95°C for 30 min in citrate buffer at pH 6.0 (CC2 Buffer, Ventana). The sections were then incubated with the primary antibody against PSA (sc-390184; mouse monoclonal antibody) at 37°C for 32 min. Detection was performed using the ultraVIEW DAB Universal Detection Kit (Ventana Medical Systems), and nuclear counterstaining was carried out with hematoxylin (Ventana Medical Systems).

### *Ex vivo* fluorescence imaging of surgical specimens

*Ex vivo* fluorescence imaging was performed using freshly resected surgical specimens (approximately 2 cm in diameter) obtained from peritoneal dissemination of patients with HGSC, including both tumor and adjacent normal tissues. A 50 μM probe solution (3 mL) in PBS containing 0.5% v/v DMSO as a cosolvent was added to a 3.5 cm dish containing a specimen so that the tissue was completely soaked with probe solution. The fluorescence images were acquired before and at 1, 3, 5, 10, 20, and 30 min after the probe application using the Maestro *in vivo* imaging system (PerkinElmer) with an excitation filter of 445–490 nm and an emission filter of 515 nm long-pass. Fluorescence intensity was quantified by drawing ROIs using the Maestro software.^(38)^

### *In vivo* fluorescence imaging in mouse models

To evaluate the ability of candidate probes to detect peritoneal dissemination of HGSC, *in vivo* fluorescence imaging was performed using a mouse xenograft model. SKOV3-Luc human ovarian cancer cells (1.5 × 10⁶ cells in 200 µL PBS) were intraperitoneally inoculated into 7-week-old female BALB/c-nu/nu mice to establish peritoneal dissemination. Tumor progression was monitored on days 14 and 21 using the IVIS imaging system (PerkinElmer) after intraperitoneal administration of 3 mg of D-luciferin potassium salt in 200 µL of PBS. On day 28, mice were injected intraperitoneally with a 100 μM solution of EK-HMRG, NA-HMRG or DA-HMRG probe in PBS containing 0.5% v/v DMSO as a cosolvent (300 µL). After 15 min, mice were sacrificed by exposure to CO_2_ and the abdominal cavities were exposed. Fluorescence images were obtained with the Maestro *in vivo* imaging system (PerkinElmer) using the blue-filter setting (excitation, 445 to 490 nm; emission, 515 nm long-pass). The tunable filter was automatically stepped in 10 nm increments, from 500 to 800 nm, while the camera sequentially captured images at each wavelength.

### *Ex vivo* fluorescence imaging in an operating room

*Ex vivo* fluorescence imaging was performed immediately after surgical resection of the HGSC specimens in an operating room. Fresh surgical specimens obtained from peritoneal dissemination of patients with HGSC, including those who had received neoadjuvant chemotherapy with carboplatin, paclitaxel, and bevacizumab, were used. A 50 μM probe solution (3 mL) in PBS containing 0.5% v/v DMSO as a cosolvent was applied to specimens. The fluorescence images were recorded using an in-house-built portable fluorescence imager. Gain value was set at 700. Exposure time was set at 0.1 sec.

### Statistical analysis

Data processing and ROC curve analyses were performed using Python (Python Software Foundation). Threshold values were determined using Youden’s index. The smoothed bootstrap method was applied to compute confidence intervals for the ROC curve using 20,000 resamples.

## Results

### Screening of fluorescence probes to identify efficient probe/biomarker enzyme pairs for the detection of peritoneal dissemination of HGSC

To discover fluorescence probe/biomarker enzyme pairs suitable for the selective and efficient detection of peritoneal dissemination of HGSC, we performed a two-step screening using surgically resected clinical specimens (**Fig. 1C**).

**Figure 1.**
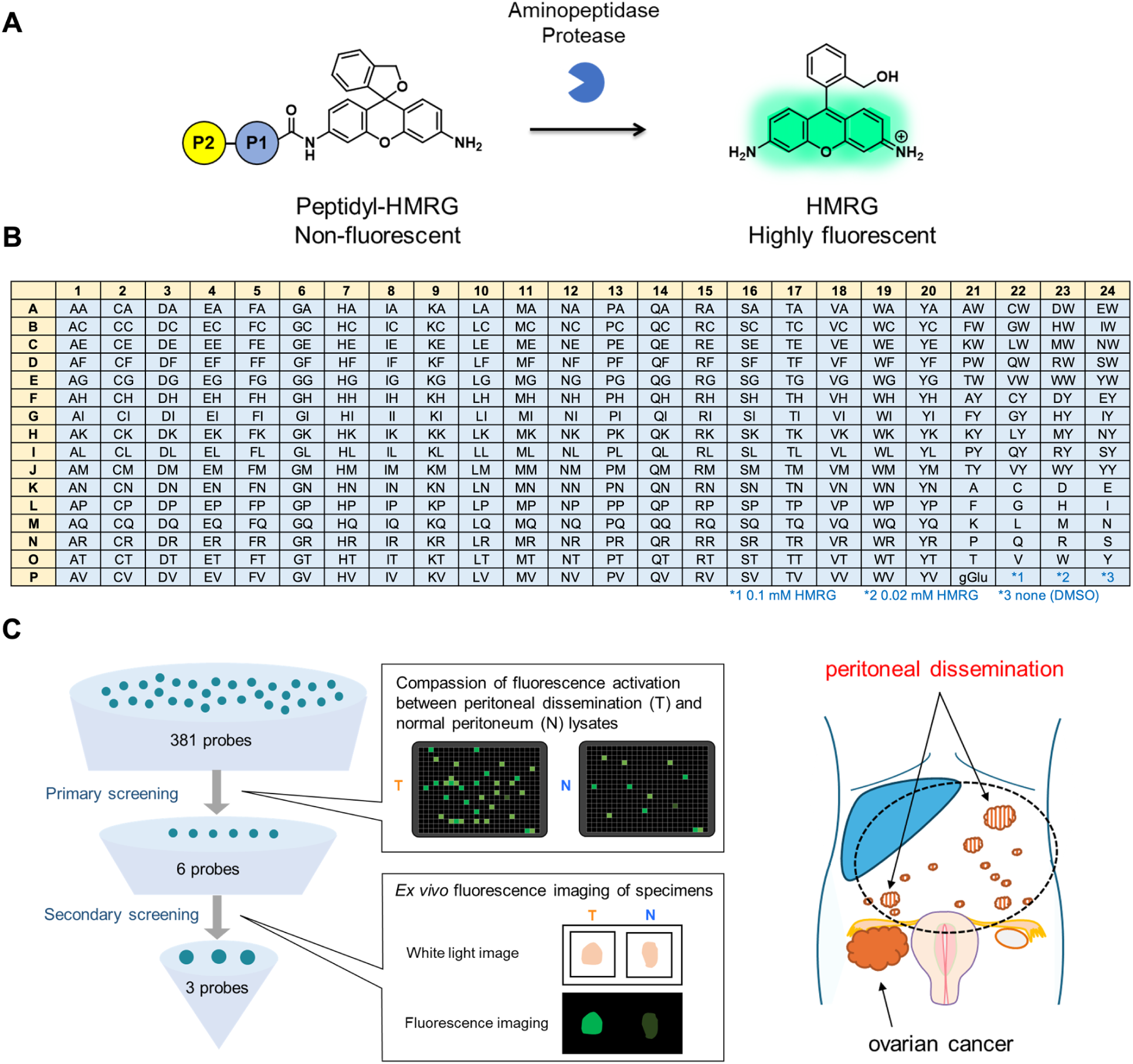
HMRG-based fluorescence probe library and flowchart of screening for peritoneal dissemination of HGSC. (**A**) Reaction scheme of peptidyl-HMRG with aminopeptidases or proteases. (**B**) List of the fluorescence probes in the library. Columns and rows represent amino acids at the P1 and P2 positions, respectively. (**C**) Flowchart of screening using human surgical peritoneal dissemination and adjacent normal peritoneum.

In the primary screening, we screened our aminopeptidases and proteases-reactive fluorescence probe library consisting of 381 probes with lysates of surgically resected human HGSG tumor and adjacent non-tumor tissue lysates from five different patients. For this study, we partly refined the reported probe library^(34)^ (**Fig. 1A** and **B**; Supplementary Table S2). Fluorescence intensity was measured over 3 h after the probe application (Supplementary Fig. S1). For the selection of cancer imaging probes, it is desirable to select probes that exhibit a large fluorescence increase in tumor tissues with high T/N. To identify such probes, we selected candidate probes using the following two criteria: CVR_3 h – 0 h_ of tumor > 0.1 and T/N > 2.0. Among the screened probes, 14 probes satisfied both criteria in at least three of the five patients. EK-HMRG, NA-HMRG and DA-HMRG, which met the criteria in all five patients, were selected as the principal candidate probes. NK-HMRG and KQ-HMRG, which met the criteria in four patients, and KV-HMRG, which met the criteria in three patients, were also selected for further evaluation to ensure diversity in amino acid sequences (**Table 1**).

**Table 1.** Candidate probes in primary screening. Probes that met the predefined criteria— CVR_3 h – 0 h_ of tumor > 0.1 and (2) T/N > 2.0—in at least three of five patients were selected. T/N, tumor/normal ratio. CVR, conversion rate of probes.

| Amino acid sequence<br>(P2P1) | Patient<br>No.1 |  | Patient<br>No.2 |  | Patient<br>No.3 |  | Patient<br>No.4 |  | Patient<br>No.5 |  |
| --- | --- | --- | --- | --- | --- | --- | --- | --- | --- | --- |
|  | T/N | CVR | T/N | CVR | T/N | CVR | T/N | CVR | T/N | CVR |
| EK | 2.2 | 0.13 | 2.9 | 0.14 | 3.2 | 0.33 | 3.4 | 0.4 | 7.4 | 0.69 |
| NA | 2.1 | 0.16 | 3.5 | 0.15 | 2.5 | 0.21 | 2.8 | 0.35 | 6.1 | 0.66 |
| DA | 2.1 | 0.12 | 2.9 | 0.15 | 3.2 | 0.25 | 3.0 | 0.38 | 5.8 | 0.47 |
| NK |  |  | 2.8 | 0.28 | 2.2 | 0.71 | 2.2 | 0.71 | 3.1 | 0.77 |
| KQ |  |  | 3.9 | 0.17 | 2.2 | 0.13 | 3.3 | 0.25 | 4.7 | 0.27 |
| KV |  |  |  |  | 3.9 | 0.16 | 4.1 | 0.22 | 5.4 | 0.26 |

In the secondary screening, these six candidates were evaluated by *ex vivo* fluorescence imaging using 21 freshly resected paired tumor and peritumoral tissues from 19 HGSC patients. Fluorescence imaging was performed over 30 min, and fluorescence intensity was quantified by drawing regions of interest (ROIs) (**Fig. 2A**). These probes mostly exhibited a markedly higher fluorescence increase in tumor tissues than in normal tissues (**Fig. 2B** and **C**). To evaluate the diagnostic performance of the six selected probes, we calculated sensitivity and specificity using receiver operating characteristic curve analysis based on the fluorescence intensity values.^(38)^ Among the 6 candidates, EK-HMRG, NA-HMRG and DA-HMRG exhibited especially high detection performance at 30 min (**Fig. 3**; Supplementary Fig. S2 and S3). The sensitivity and specificity of EK-HMRG were calculated to be 92.3% and 100%, respectively, with an AUC of 0.992 (95% confidence interval [CI], 0.923-1.000). Both NA-HMRG and DA-HMRG exhibited a sensitivity and specificity of 100% and an AUC of 1.000 (95% CI: 0.946–1.000 and 0.881–1.000, respectively). Thus, all three probes showed excellent diagnostic performance in differentiating peritoneal tumor lesions from adjacent tissue and were considered promising candidates for the intraoperative imaging of HGSC dissemination.

**Figure 2.**
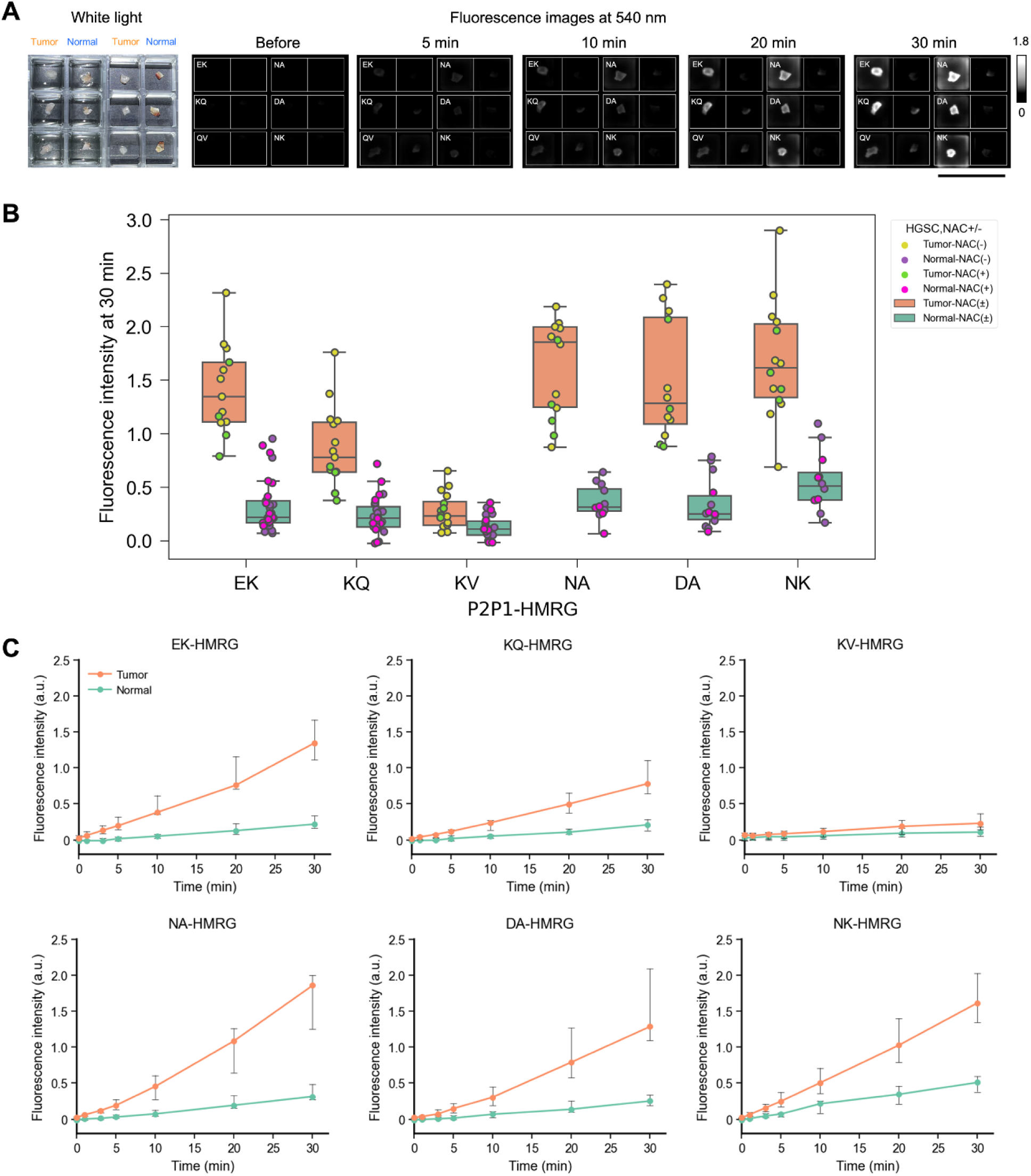
Comparison of fluorescence increase in tumor and adjacent normal peritoneal tissues in the secondary screening. (**A**) Example of screening using surgically resected peritoneal dissemination lesions and adjacent normal peritoneum. Probe solution was prepared with PBS (−) containing 0.5% v/v DMSO as a cosolvent. [fluorescence probe] = 50 μM. Scale bar, 2 cm. (**B**) Analysis of fluorescence intensity in tumor and adjacent normal peritoneal tissues using 6 fluorescence probes (N = 12−16 for Tumor, N = 12−29 for Normal). Increase in fluorescence intensity was measured at 30 min after addition of fluorescence probes. Pink, purple, green, and yellow dots represent fluorescence increases in tumor tissues without neoadjuvant chemotherapy (NAC), tumor tissues with NAC, normal tissues without NAC, and tumor tissues with NAC, respectively. (**C**) Time-dependent fluorescence intensity of six candidate probes (EK-, KQ-, KV-, NA-, DA- and NK-HMRG) in peritoneal dissemination lesions and adjacent normal peritoneum. Orange lines represent fluorescence increases in peritoneal dissemination lesions. Green lines represent fluorescence increases in normal peritoneum. Data are presented as median values with error bars indicating the interquartile range (25th–75th percentiles). Outliers were excluded according to Tukey’s rule. [fluorescence probe] = 50 μM.

**Figure 3.**
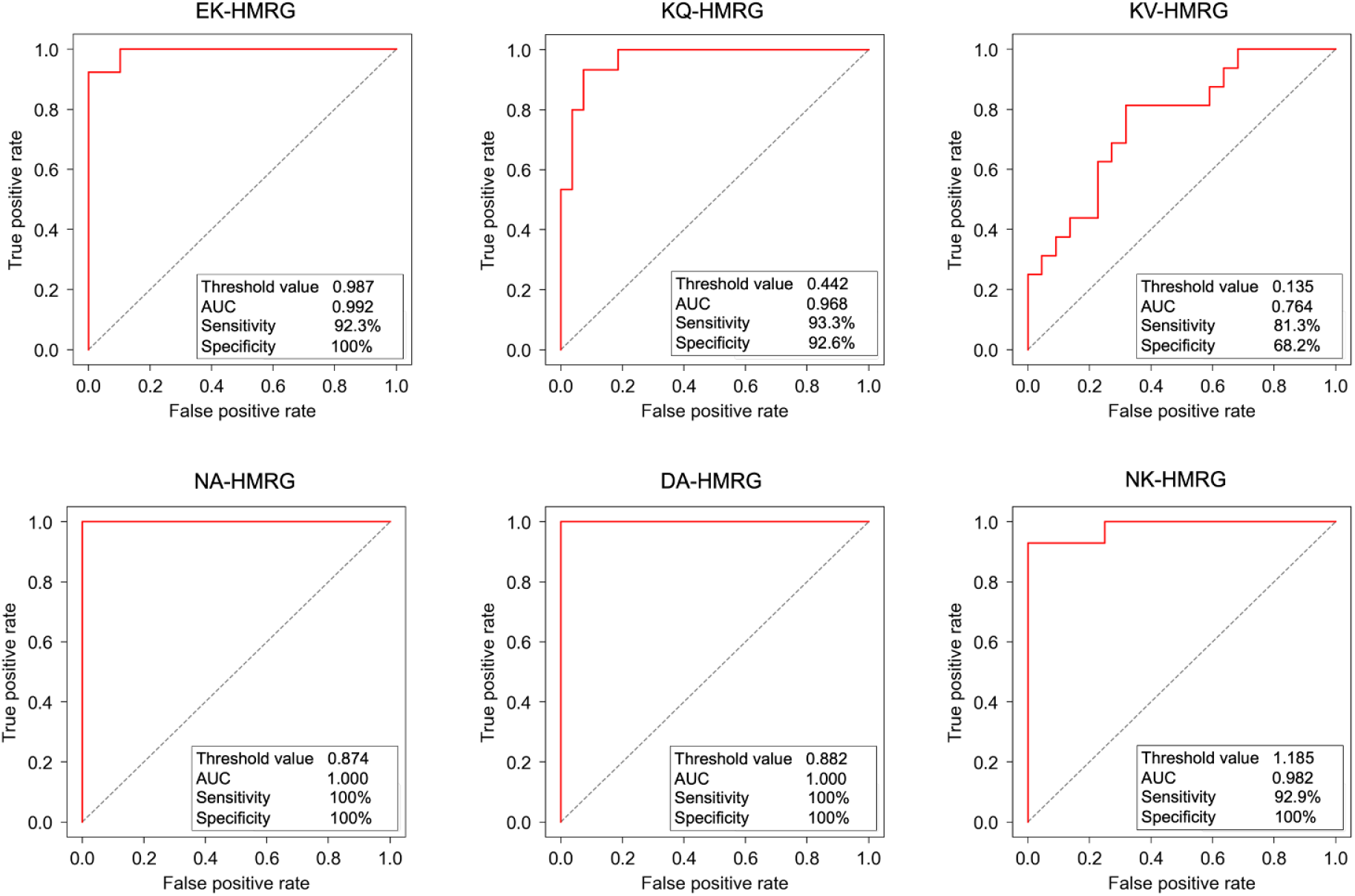
Detection performance of hit fluorescence probes. ROC curves of hit fluorescence probes. Threshold value, sensitivity, specificity and AUC of each probe were evaluated from the ROC curves. Tumor (N = 13) and Normal (N = 29) tissues were examined with EK-HMRG. Tumor (N = 15) and Normal (N = 27) tissues were examined with KQ-HMRG. Tumor (N = 16) and Normal (N = 22) tissues were examined with KV-HMRG. Tumor (N = 14) and Normal (N = 12) tissues were examined with NA-HMRG. Tumor (N = 12) and Normal (N = 14) tissues were examined with DA-HMRG. Tumor (N = 14) and Normal (N = 12) tissues were examined with NK-HMRG.

### Target identification of the hit fluorescence probes

In order to identify the target enzyme of the hit probes, we carried out a diced electrophoresis gel assay, which is a combination analysis of 2D-gel fluorometric assay and peptide mass fingerprinting,^(39)^ using EK-HMRG and lysates from peritoneal dissemination lesions of HGSC. The lysates were placed in gels and electrophoresed based on isoelectric point and molecular weight. After incubation with EK-HMRG, we observed a single fluorescent spot on the gel, and this was identified as puromycin-sensitive aminopeptidase (PSA) by peptide mass fingerprinting analysis (**Fig. 4A**; Supplementary Fig. S4). This enzyme was consistently identified in multiple independent samples. All other proteins identified in fluorescent regions of the gels are summarized in Supplementary (Supplementary Table S3). These findings suggested that PSA is predominantly responsible for the activation of EK-HMRG in peritoneal dissemination lesions of HGSC.

**Figure 4.**
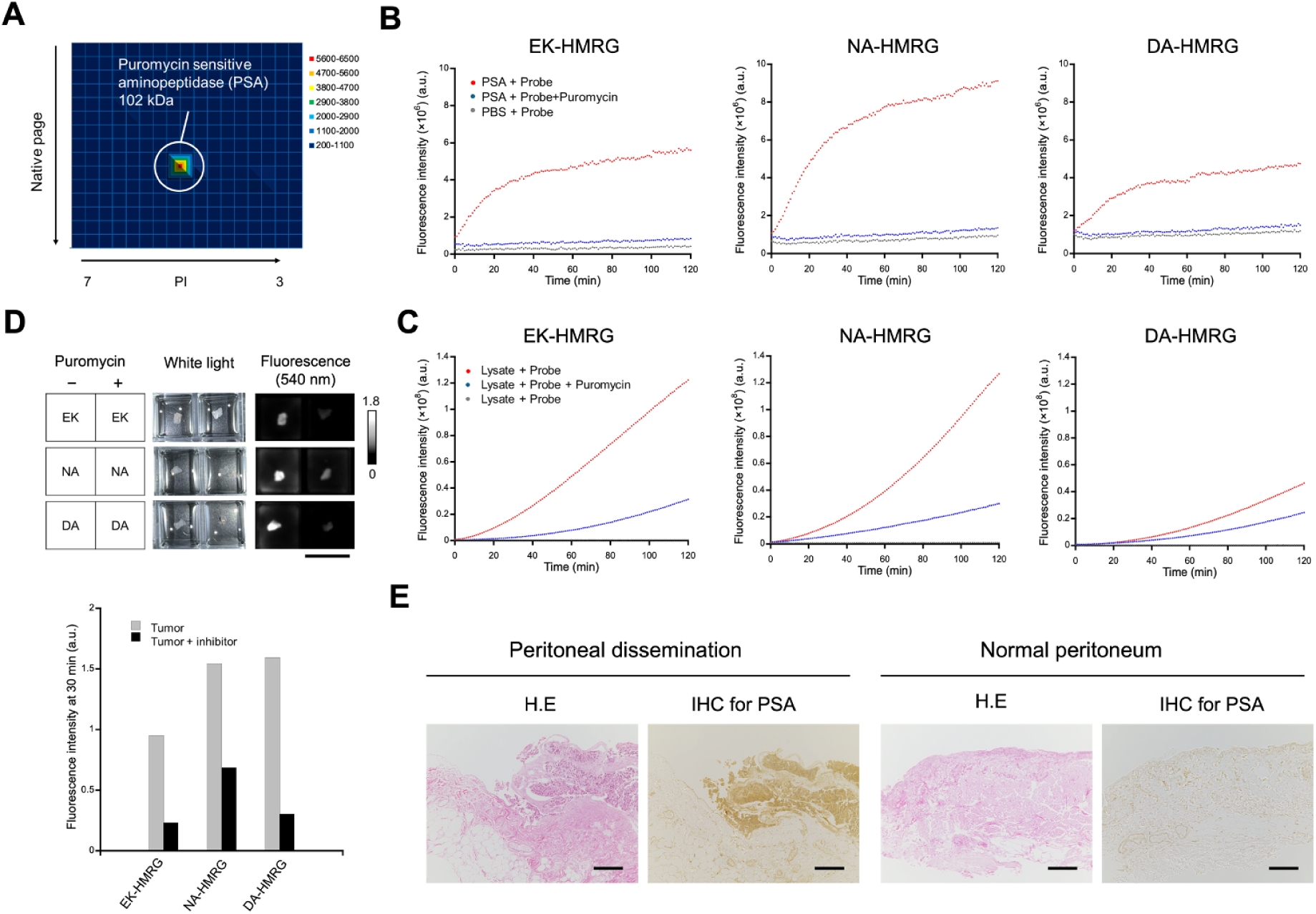
Target identification of hit fluorescence probes. (**A**) DEG assay for peritoneal dissemination lesions using EK-HMRG probe. PSA (102 kDa) was identified by peptide mass fingerprinting analysis from the fluorescent spot on 2D gel. (**B**) Time-dependent fluorescence intensity of each probe with 0.25 µg/mL PSA in the presence or absence of puromycin. Puromycin suppressed the fluorescence increase of EK-HMRG, NA-HMRG and DA-HMRG. [fluorescence probes] = 1 µM, [puromycin] = 30 µM. (**C**) Fluorescence intensity of each probe incubated with 0.1 mg/mL of tissue lysates prepared from peritoneal dissemination lesions in the presence or absence of puromycin. Puromycin suppressed the fluorescence increase of EK-HMRG, NA-HMRG and DA-HMRG. [fluorescence probes] = 1 µM, [puromycin] = 30 µM. (**D**) *Ex vivo* fluorescence images of peritoneal dissemination tissues 30 min after application of fluorescence probes in the presence and absence of puromycin (upper). Fluorescence intensity at 30 min in peritoneal dissemination tissues in the presence and absence of puromycin (bottom). Gray bars: fluorescence increase in the absence of inhibitor. Black bars: fluorescence increase in the presence of inhibitor. [fluorescence probe] = 50 μM, [puromycin] = 500 μM. Scale bars, 2 cm. (**E**) Representative images of IHC staining for PSA in peritoneal dissemination lesions and adjacent normal peritoneum. Cancer cells exhibited strong staining of PSA. Scale bars, 200 μm.

To validate the reactivity of EK-HMRG with PSA, we performed *in vitro* fluorescence assays using recombinant human PSA. EK-HMRG exhibited a rapid, large fluorescence increase in the presence of PSA, while this reaction was markedly suppressed by puromycin, a known PSA inhibitor. We also performed inhibition assays to evaluate the reactivity of NA-HMRG and DA-HMRG with PSA. As in the case of EK-HMRG, these probes exhibited a fluorescence increase in the presence of PSA, and the observed increases were strongly inhibited by puromycin, indicating that three hit probes are able to react with PSA (**Fig. 4B**).

Furthermore, we examined the fluorescence increase of these hit probes in HGSC lysates with and without puromycin. Pre-incubation with puromycin markedly suppressed fluorescence activation in a dose-dependent manner for EK-HMRG, NA-HMRG and DA-HMRG (**Fig. 4C**), supporting the view that these probes react with PSA in HGSC clinical samples. We also performed fluorescence imaging of HGSC tumor specimens using EK-HMRG, NA-HMRG and DA-HMRG in the presence and absence of puromycin. Puromycin markedly suppressed the fluorescence increase of each probe, further suggesting that PSA activity mediates the fluorescence increase (**Fig. 4D**).

For additional validation, we evaluated the expression levels of PSA in the specimens by immunohistochemical (IHC) staining. Tumor cells from peritoneal dissemination lesions of HGSC showed strong cytoplasmic PSA staining, whereas mesothelial and stromal cells in non-tumor areas exhibited weak or focal staining (**Fig. 4E**; Supplementary Fig. S5), indicating that PSA is preferentially expressed in HGSC tumor tissue. Overall, these results demonstrate that PSA is the target enzyme of EK-HMRG, NA-HMRG and DA-HMRG. To our knowledge, this enzyme has not previously been reported as a biomarker enzyme in peritoneal dissemination of HGSC.

### *Ex vivo* fluorescence imaging of peritoneal dissemination lesions in clinical specimens using the hit fluorescence probes

To assess the clinical applicability of the selected probes, we performed *ex vivo* fluorescence imaging using clinical specimens from patients with HGSC that are expected to contain both normal peritoneum and peritoneal dissemination lesions. A solution of EK-HMRG, NA-HMRG or DA-HMRG was topically applied to each specimen, and fluorescence images were acquired. The fluorescence intensity of tumor regions began to increase over 3-10 min (**Fig. 5**). All probes were able to clearly visualize tumor regions less than 1 mm in a diameter within 10 min. Histopathological examination confirmed that the highly fluorescent regions coincided with pathological tumor regions, while non-fluorescent regions were diagnosed as normal tissues. These findings indicated that all three probes can rapidly and sensitively visualize peritoneal dissemination lesions of HGSC just a few millimeters in size within an intraoperative time scale.

**Figure 5.**
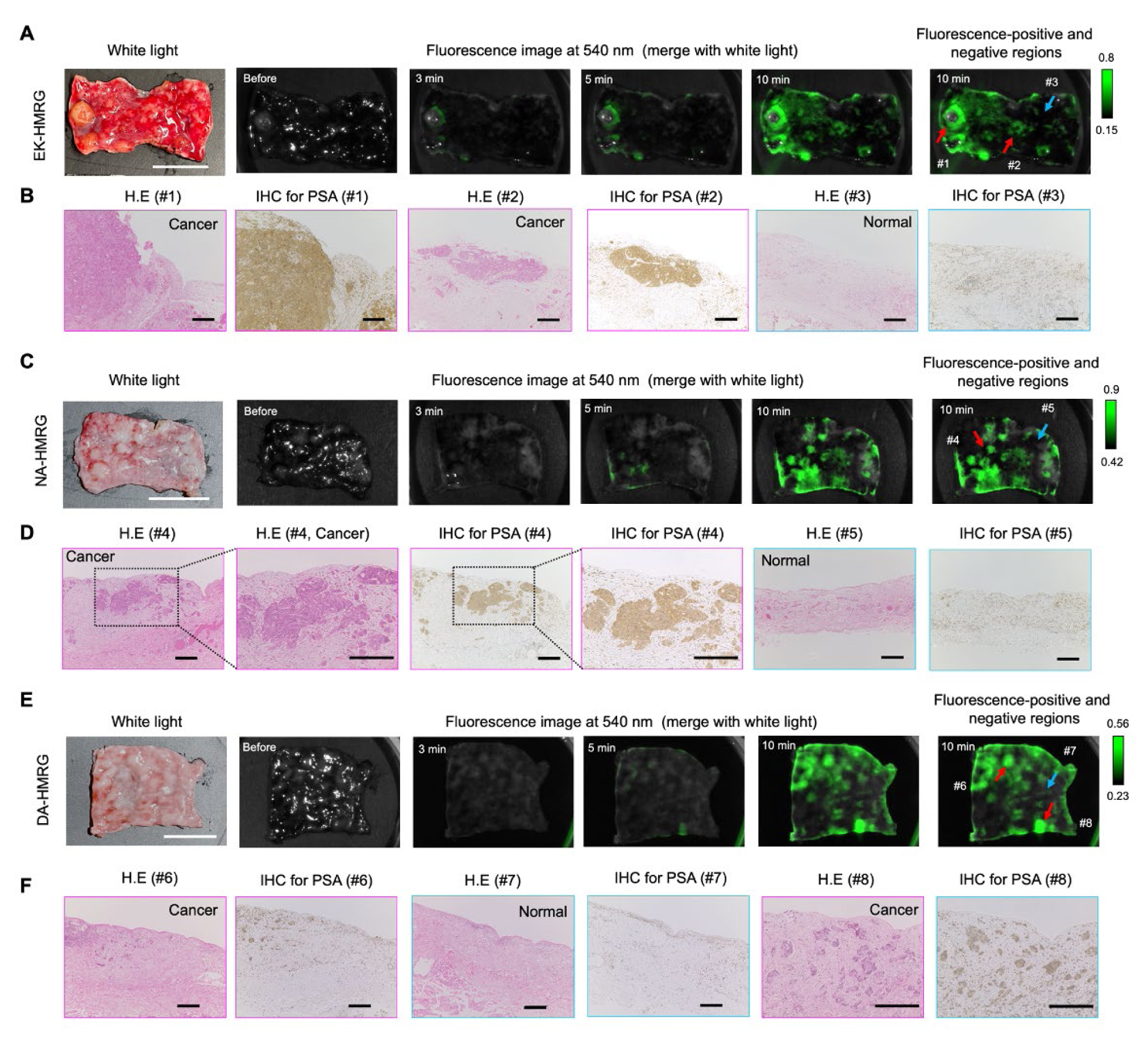
*Ex vivo* fluorescence imaging of HGSC peritoneal dissemination in clinical specimens. (**A**) Time-dependent fluorescence images of surgically resected fresh human specimens containing both peritoneal dissemination lesions and adjacent normal peritoneum after administration of EK-HMRG. Probe solution was prepared with PBS (−) containing 0.5% v/v DMSO as a cosolvent. [fluorescence probe] = 50 μM. Scale bar, 2 cm. (**B**) Histological analysis and IHC analysis for PSA of ROI #1 and #2 with strong fluorescence activation (pink box) or ROI #3 with almost no fluorescence activation (blue box). Scale bars, 200 μm. (**C**) Time-dependent fluorescence images of surgically resected fresh human specimens containing both peritoneal dissemination lesions and adjacent normal peritoneum after administration of NA-HMRG. Probe solution was prepared with PBS (−) containing 0.5% v/v DMSO as a cosolvent. [fluorescence probe] = 50 μM. Scale bar, 2 cm. (**D**) Histological analysis and IHC analysis for PSA of ROI #4 with strong fluorescence activation (pink box) or ROI #5 with almost no fluorescence activation (blue box). Scale bars, 200 μm. (**E**) Time-dependent fluorescence image of surgically resected fresh human specimens containing both peritoneal dissemination lesions and adjacent normal peritoneum after administration of DA-HMRG. Probe solution was prepared with PBS (−) containing 0.5% v/v DMSO as a cosolvent. [fluorescence probe] = 50 μM. Scale bar, 2 cm. (**F**) Histological analysis and IHC analysis for PSA of ROI #6 and #8 with strong fluorescence activation (pink box) or ROI #7 with almost no fluorescence activation (blue box). Scale bars, 200 μm.

### *In vivo* fluorescence imaging of peritoneal metastases in a mouse model using the hit probes

In clinical applications, these probes are expected to be topically applied to the peritoneal cavity. To evaluate the efficiencies of the three hit probes in such applications, we performed *in vivo* fluorescence imaging using a SKOV-3 peritoneal dissemination mouse model. EK-HMRG, NA-HMRG and DA-HMRG (100 µM, 300 µL each) were intraperitoneally administered. At 15 min after probe application, mice were sacrificed, the abdominal cavity was exposed and fluorescence imaging was performed. For all three probes, strong fluorescence signals were observed in tumor nodules scattered on the peritoneal surfaces, with minimal background fluorescence in normal tissues, such as intestines (**Fig. 6**). These findings demonstrate that PSA-targeting probes can selectively visualize peritoneal dissemination lesions of HGSC *in vivo*. Notably, these probes can visualize lesions just a few millimeters in size, which are difficult to identify with the naked eye. These results support the potential clinical utility of EK-HMRG, NA-HMRG and DA-HMRG for rapid and sensitive intraoperative fluorescence detection of peritoneal metastases of HGSC.

**Figure 6.**
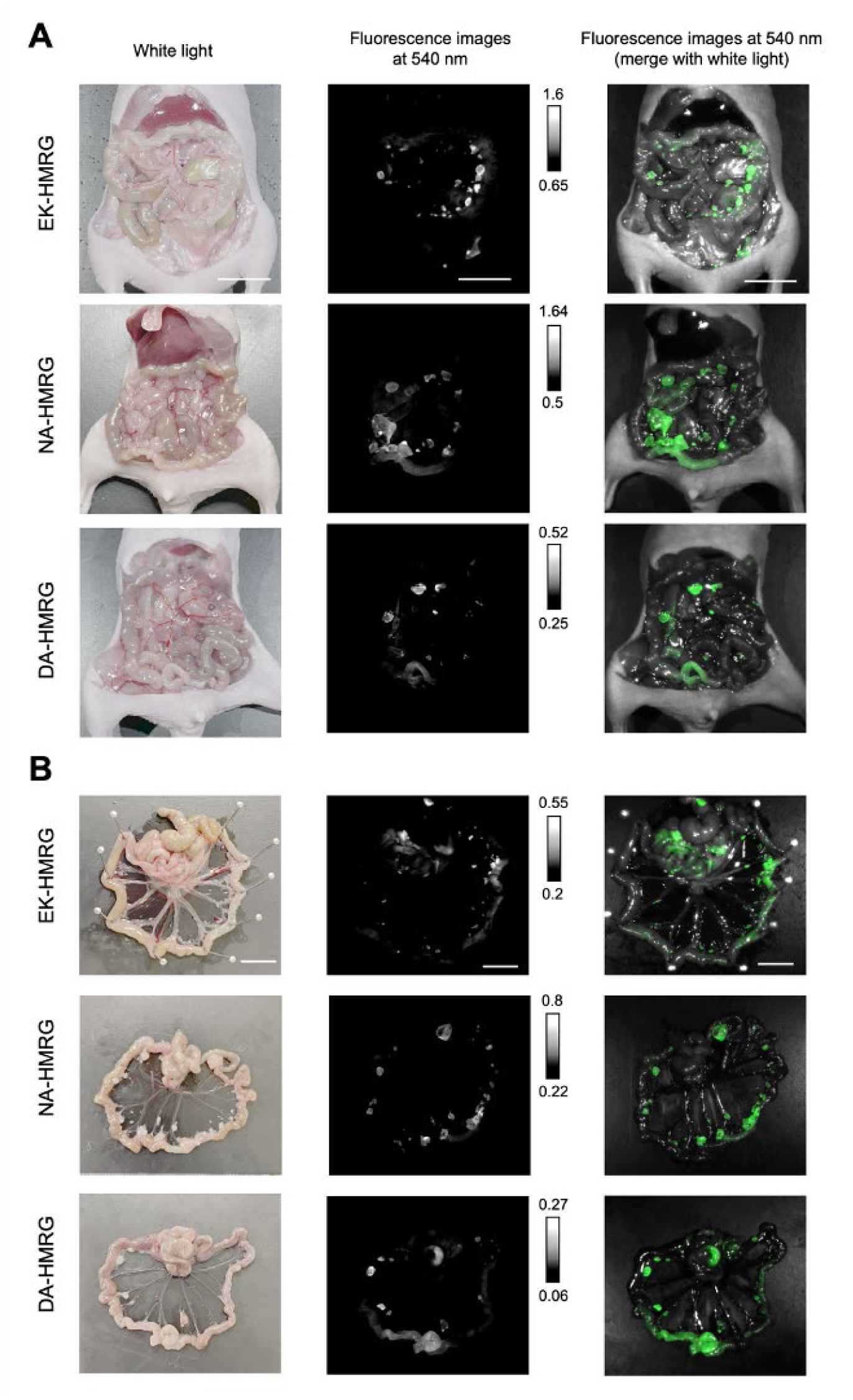
*In vivo* fluorescence imaging of peritoneal dissemination in mouse model. (**a**) Representative white light, fluorescence and composite images of peritoneum 15 min after intraperitoneal injection of EK-HMRG, NA-HMRG or DA-HMRG in the SKOV3 ovarian cancer metastasis mouse model. 300 µL of probe solution was prepared with PBS (−) containing 0.5% *v/v* DMSO as a cosolvent. [fluorescence probes] = 100 μM. Scale bars, 2 cm. (**b**) *Ex vivo* fluorescence imaging of peritoneal dissemination using EK-HMRG, NA-HMRG or DA-HMRG in SKOV3 ovarian cancer metastasis mouse model. Guts and peritoneum were resected from (**a**). SKOV3 cancer lesions were clearly visualized. Probe solution was prepared with PBS (−) containing 0.5% v/v DMSO as a cosolvent. [fluorescence probe] = 100 μM. Scale bars, 2 cm.

### *Ex vivo* fluorescence imaging of large clinical specimens

To evaluate intraoperative applicability, we evaluated EK-HMRG using relatively large, fresh specimens immediately after surgical resection in an operating room. The specimens were obtained from patients who had received neoadjuvant chemotherapy with carboplatin, paclitaxel and bevacizumab. Post-chemotherapy CT imaging showed therapeutic responses, including reduced peritoneal thickening, disappearance of ascites, and shrinkage of omental dissemination and peritoneal dissemination nodules, while no obvious residual peritoneal dissemination was detected on the vesicouterine peritoneum (**Fig. 7A** and **B**). EK-HMRG solution was topically applied to the specimens, which were expected to contain both tumor and normal peritoneum, as well as to uterus, ovary and sigmoid colon, and fluorescence images were acquired (**Fig. 7C** and **D**). EK-HMRG clearly exhibited three fluorescent regions on the peritoneum over 10-20 min (**Fig. 7D** and **E**). Histological analysis confirmed that these fluorescent regions contained cancer tissues and represented peritoneal dissemination lesions (**Fig. 7F**). These results demonstrate that EK-HMRG can sensitively detect minute peritoneal dissemination lesions that were not clearly detectable by CT imaging within an intraoperative time scale.

**Figure 7.**
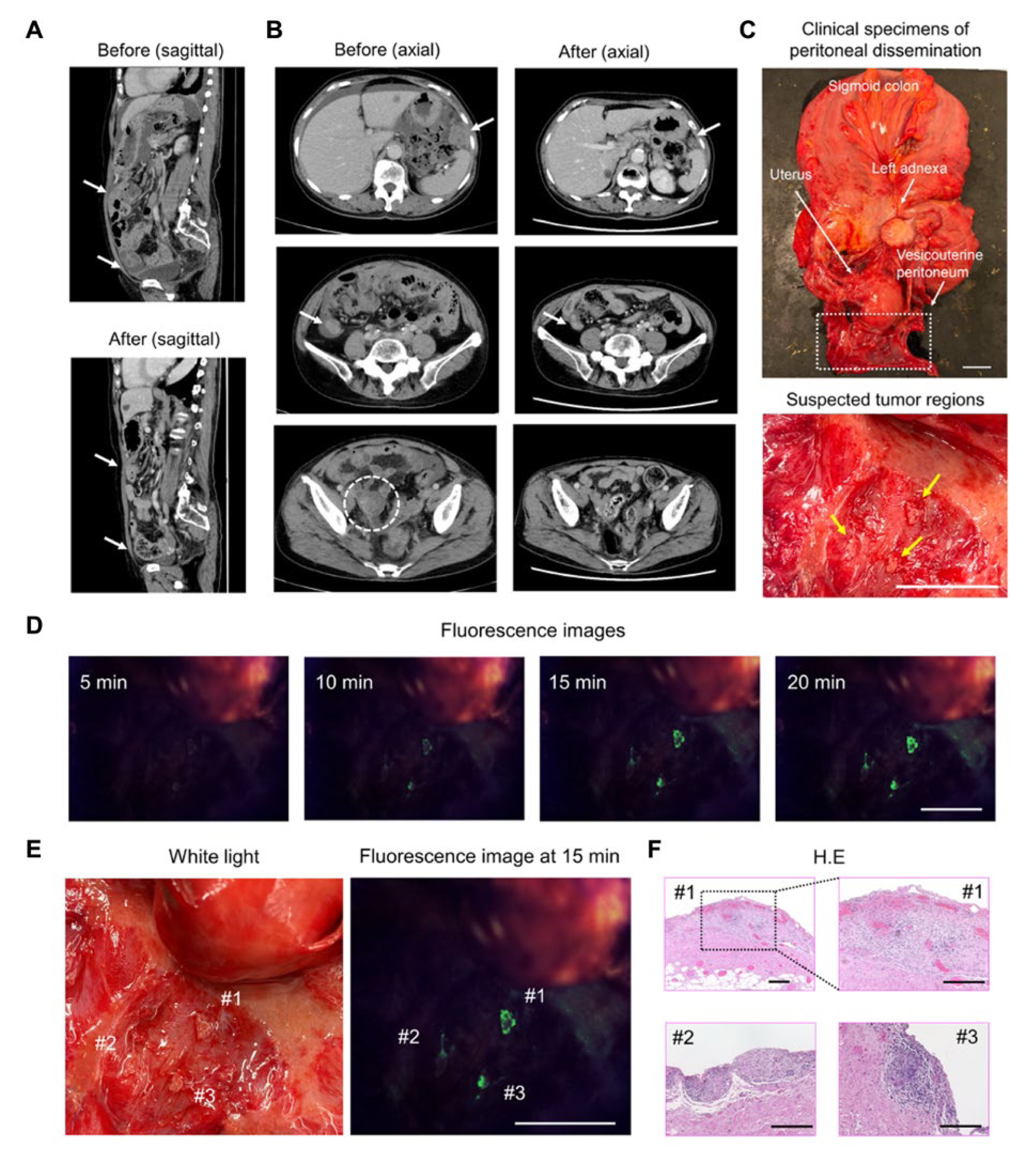
Fluorescence imaging of metastatic regions in the resected HGSC specimen from a patient who underwent interval debulking surgery. (**A**) Sagittal CT images before and after neoadjuvant chemotherapy. Upper white arrows indicate omental dissemination. Lower white arrows indicate peritoneal thickening and ascites. These findings were reduced, and no obvious nodules were identified in the vesicouterine peritoneum. (**B**) Axial CT images before and after neoadjuvant chemotherapy. White arrows indicate peritoneal dissemination nodules, which became smaller after chemotherapy. Dotted white circles indicate the right ovarian tumor and its absence after staging laparoscopy. (**C**) Macroscopic findings of the resected specimen. The right adnexa was removed during staging laparoscopy. Neoadjuvant chemotherapy was subsequently administered, followed by interval debulking surgery. The uterus, sigmoid colon, and left adnexa were adherent due to tumor invasion and were resected en bloc. Scale bars, 2 cm. (**D**) Time-dependent fluorescence images of surgically resected fresh human specimens containing both peritoneal dissemination lesions and adjacent normal peritoneum after administration of EK-HMRG. Probe solution was prepared with PBS (−) containing 0.5% v/v DMSO as a cosolvent. [fluorescence probe] = 50 μM. Scale bar, 2 cm. (**E**) Magnified white light and fluorescence images of the fluorescent regions 15 min after probe application. Scale bar, 2 cm. (**F**) Histological analysis of fluorescent regions in the evaluated specimen. Fluorescent regions (#1-#3) contained HGSC cancer tissues. Scale bars, 200 µm.

## Discussion

High-grade serous ovarian cancer (HGSC) is typically diagnosed at an advanced stage, when it is often accompanied with extensive peritoneal dissemination.^(1–4)^ Complete cytoreductive surgery is the most important prognostic factor,^(5–8)^ but reliable intraoperative detection of minute peritoneal lesions remains challenging. Conventional preoperative imaging modalities such as CT, MRI, and PET/CT have limited spatial resolution, with CT failing to detect implants smaller than approximately 10 mm, and MRI and PET-CT showing reduced sensitivity for subcentimeter nodules.^(10–15)^ Intraoperatively, white-light inspection and indocyanine green (ICG) fluorescence visualization also lack sufficient specificity, and microscopic lesions are frequently missed.^(21–23)^

In this study, we developed novel enzyme-activatable fluorescence probes, EK-HMRG, NA-HMRG and DA-HMRG, targeting elevated PSA activity in HGSC. These probes were able to visualize cancer lesions in clinical specimens *ex vivo* within approximately 10 min with a high T/N ratio and excellent diagnostic accuracy. Importantly, our probes enabled highly sensitive visualization of nodules a few millimeters in size that are often undetectable by conventional imaging or intraoperative inspection. This capability represents a major advantage over current modalities and in the clinical context has the potential for enabling complete cytoreduction. Given that residual disease may persist despite apparently complete gross resection^(9)^, improved visualization of minute lesions may facilitate more accurate assessment of residual disease and support surgical decision-making during cytoreductive surgery.

Agents such as ICG and Cytalux are clinically approved for fluorescence-guided detection of ovarian cancer lesions, but both remain limited by nonspecific accumulation in normal tissues, resulting in inevitable background fluorescence or false-positive signals.^(21–23,26,40)^. Cytalux targets folate receptor-α (FRα), which is overexpressed in more than 90% of epithelial ovarian cancers, but it still showed a patient-level false-positive rate of 24.8%.^(24–26)^ Although pH-activatable probes and structure-inherent targeting probes have also been developed to enhance tumor specificity in ovarian cancer mouse models, their performance in human ovarian cancer specimens remains largely unexplored. ^(41–43)^

Compared with these previously reported probes, our PSA-reactive activatable probes exhibited extremely high sensitivity and specificity in human surgical specimens after topical administration. From a clinical perspective, the ability to visualize peritoneal dissemination rapidly and specifically without systemic administration represents a major advantage. Such probes could facilitate real-time intraoperative decision-making, shorten operative time, and improve the completeness of cytoreduction. In this context, since the probes are expected to be used for real-time fluorescence imaging during surgery, it is necessary to achieve high brightness and T/N on an intraoperative time scale. For these reasons, probes with high fluorescence quantum yield and high visibility are preferred.^(31)^ Our probes utilizing the bright HMRG fluorophore, in addition to the rapid catalytic turnover by the target PSA, meet these criteria for intraoperative fluorescence imaging.

Through target validation analyses, we identified PSA as the main target enzyme for all three probes. This enzyme, which cleaves N-terminal amino acids from peptides, is predominantly localized in the cytosol and shows high activity in proliferating tumor cells.^(44,45)^ Although PSA has been reported to be overexpressed in other malignancies, to our knowledge this is the first study to demonstrate that PSA can work as an efficient biomarker enzyme for HGSC peritoneal dissemination.^(34,37,46,47)^ Notably, a clear fluorescence contrast was also observed in specimens obtained from patients who had received neoadjuvant chemotherapy, such as carboplatin, paclitaxel, and bevacizumab, indicating that PSA activity remains detectable even after these treatments. This finding indicated that PSA-targeting fluorescence imaging can be applied to both lesions without neoadjuvant chemotherapy and residual lesions after chemotherapy, suggesting its utility as robust biomarker in HGSC. Probes possessing KK or QA substrates, which we previously reported as efficient PSA-reactive probes,^(34,37,46)^ did not emerge as top hits in this screening, probably due to unspecific activation in non-tumor peritoneum tissues mediated by other proteases and aminopeptidases.

Further validation will be important to support clinical translation. A larger multicenter study is needed to confirm diagnostic performance and generalizability, and prospective intraoperative studies will be required to establish feasibility and safety in humans. In addition, broader profiling of PSA expression and activity across normal organs will further define the specificity and clinical applicability of PSA-targeting probes. Together, these studies are expected to provide a basis for future first-in-human intraoperative imaging trials. In conclusion, we have identified PSA as a promising biomarker enzyme in HGSC and demonstrated the utility of topical activatable fluorescence probes targeting PSA for rapid and highly sensitive detection of HGSC peritoneal dissemination. These findings support the potential utility of PSA-targeted activatable fluorescence probes for fluorescence-guided detection of HGSC peritoneal dissemination and may contribute to more accurate intraoperative assessment of residual disease during cytoreductive surgery.

### Translational Relevance

Complete cytoreductive surgery is a major determinant of survival in patients with advanced high-grade serous ovarian carcinoma (HGSC), yet minute peritoneal disseminated lesions are often difficult to identify intraoperatively. This study identifies puromycin-sensitive aminopeptidase (PSA) as an activity-elevated biomarker enzyme of HGSC peritoneal dissemination and demonstrates the translational potential of three PSA-activatable fluorescence probes, EK-HMRG, NA-HMRG and DA-HMRG. Upon topical application, these probes enabled rapid and highly sensitive visualization of small peritoneal lesions in human clinical specimens, as well as in a peritoneal dissemination mouse model, on an intraoperative time scale. Because this approach does not require systemic administration and provides real-time lesion contrast, PSA-targeting activatable fluorescence imaging may help surgeons detect otherwise occult peritoneal dissemination and support more complete cytoreductive surgery. These findings strongly support the further clinical development of topical enzyme-activatable fluorescence imaging as an intraoperative strategy to improve cytoreductive surgery in advanced HGSC.

## Acknowledgements

This work was supported by Grants-in-Aid for Scientific Research (KAKENHI) (JP19H05632 and JP24H00050 to Y. U., JP22K20528 and JP23K14317 to K. F.), Japan Agency for Medical Research and Development (AMED) Project for Promotion of Cancer Research and Therapeutic Evolution (P-PROMOTE) (JP26ama221159h0001 to K. F.), Japan Science and Technology Agency (JST) Mirai Program (JPMJMI24G2 to Y. U.), and JST Moonshot Research and Development Program (JPMJMS2022 to Y. U.). We are grateful to Atsuki Abe, Yuko Hirata, Aika Nanjo, Kayo Hasegawa, Reiko Tsuchiya, Akemi Matsumoto, Kazunari Fujino and Yosuke Ito for their technical assistance in the screening experiments. We thank the members of the Laboratory of Morphology and Image Analysis, Biomedical Research Core Facilities, Juntendo University Graduate School of Medicine, for their technical assistance with microscopy. We thank the members of the Laboratory of Molecular and Biochemical Research, Biomedical Research Core Facilities, Juntendo University Graduate School of Medicine, for their technical assistance.

## Author contributions

H.S., K.F., E.Y. and Y.U. designed the study. E.Y., D.O. and Y.T. performed surgical resection and extracted sample specimens. H.S., K.F. and Y.S. performed fluorescence imaging. H.S. performed immunohistochemical staining. H.S., K.F., E.Y., T.U., T.K. and Y.U. analyzed the data. H.S., K.F. and Y.U. wrote the manuscript with inputs from all the authors. T.H. made the pathological diagnoses. K.F., E.Y., Y.U. and T.Y. supervised the study.

