## Supplementary Information for "Topical Activatable Fluorescence Probes for Rapid Intraoperative Detection of Peritoneal Dissemination in High-Grade Serous Ovarian Carcinoma"

### Supplementary Figures.

Tumor conversion rate
3 h

Patient No. 1

| | 1 | 2 | 3 | 4 | 5 | 6 | 7 | 8 | 9 | 10 | 11 | 12 | 13 | 14 | 15 | 16 | 17 | 18 | 19 | 20 | 21 | 22 | 23 | 24 |
|---|---|---|---|---|---|---|---|---|---|---|---|---|---|---|---|---|---|---|---|---|---|---|---|---|
| A | 0.32 | 0.02 | 0.12 | 0.10 | 0.11 | 0.20 | 0.35 | 0.18 | 0.33 | 0.22 | 0.25 | 0.16 | 0.23 | 0.31 | 0.35 | 0.25 | 0.21 | 0.31 | 0.13 | 0.18 | 0.01 | 0.01 | 0.00 | 0.01 |
| B | 0.01 | 0.00 | 0.01 | 0.01 | 0.00 | 0.04 | 0.01 | 0.01 | 0.02 | 0.01 | 0.01 | 0.01 | 0.02 | 0.01 | 0.00 | 0.01 | 0.01 | 0.01 | 0.00 | 0.00 | 0.01 | 0.04 | 0.01 | 0.01 |
| C | 0.10 | 0.00 | 0.00 | 0.00 | 0.02 | 0.03 | 0.02 | 0.01 | 0.01 | 0.02 | 0.02 | 0.01 | 0.03 | 0.01 | 0.02 | 0.03 | 0.02 | 0.05 | 0.01 | 0.01 | 0.01 | 0.00 | 0.00 | 0.01 |
| D | 0.04 | 0.00 | 0.01 | 0.01 | 0.01 | 0.05 | 0.04 | 0.01 | 0.05 | 0.03 | 0.01 | 0.02 | 0.03 | 0.03 | 0.05 | 0.03 | 0.03 | 0.01 | 0.01 | 0.02 | 0.03 | 0.02 | 0.01 | 0.02 |
| E | 0.09 | 0.00 | 0.01 | 0.01 | 0.02 | 0.09 | 0.12 | 0.03 | 0.03 | 0.02 | 0.02 | 0.02 | 0.05 | 0.05 | 0.04 | 0.06 | 0.03 | 0.09 | 0.01 | 0.02 | 0.01 | 0.02 | 0.00 | 0.01 |
| F | 0.10 | 0.00 | 0.01 | 0.01 | 0.01 | 0.14 | 0.06 | 0.01 | 0.02 | 0.01 | 0.01 | 0.03 | 0.16 | 0.02 | 0.02 | 0.12 | 0.10 | 0.10 | 0.00 | 0.01 | 0.03 | 0.00 | 0.03 | 0.04 |
| G | 0.06 | 0.02 | 0.01 | 0.01 | 0.01 | 0.04 | 0.06 | 0.03 | 0.08 | 0.02 | 0.02 | 0.01 | 0.02 | 0.03 | 0.10 | 0.04 | 0.03 | 0.02 | 0.01 | 0.03 | 0.01 | 0.03 | 0.04 | 0.03 |
| H | 0.10 | 0.01 | 0.09 | 0.13 | 0.24 | 0.10 | 0.32 | 0.21 | 0.44 | 0.24 | 0.19 | 0.25 | 0.15 | 0.33 | 0.33 | 0.26 | 0.32 | 0.28 | 0.15 | 0.20 | 0.09 | 0.02 | 0.01 | 0.09 |
| I | 0.10 | 0.00 | 0.03 | 0.05 | 0.13 | 0.02 | 0.09 | 0.06 | 0.22 | 0.02 | 0.08 | 0.06 | 0.07 | 0.07 | 0.13 | 0.10 | 0.04 | 0.05 | 0.03 | 0.06 | 0.06 | 0.04 | 0.04 | 0.03 |
| J | 0.09 | 0.00 | 0.07 | 0.07 | 0.09 | 0.08 | 0.14 | 0.08 | 0.19 | 0.14 | 0.08 | 0.09 | 0.16 | 0.09 | 0.17 | 0.14 | 0.18 | 0.13 | 0.11 | 0.07 | 0.05 | 0.03 | 0.03 | 0.03 |
| K | 0.12 | 0.00 | 0.00 | 0.00 | 0.01 | 0.11 | 0.07 | 0.03 | 0.03 | 0.01 | 0.01 | 0.01 | 0.12 | 0.02 | 0.02 | 0.10 | 0.05 | 0.11 | 0.00 | 0.01 | 0.08 | 0.00 | 0.01 | 0.00 |
| L | 0.21 | 0.00 | 0.10 | 0.12 | 0.05 | 0.21 | 0.20 | 0.13 | 0.27 | 0.10 | 0.11 | 0.15 | 0.24 | 0.08 | 0.24 | 0.20 | 0.17 | 0.16 | 0.05 | 0.05 | 0.01 | 0.02 | 0.00 | 0.03 |
| M | 0.14 | 0.00 | 0.01 | 0.04 | 0.03 | 0.18 | 0.12 | 0.05 | 0.09 | 0.03 | 0.07 | 0.03 | 0.23 | 0.10 | 0.09 | 0.17 | 0.08 | 0.12 | 0.03 | 0.03 | 0.18 | 0.03 | 0.09 | 0.01 |
| N | 0.06 | 0.01 | 0.09 | 0.10 | 0.20 | 0.04 | 0.15 | 0.18 | 0.33 | 0.17 | 0.26 | 0.14 | 0.03 | 0.23 | 0.18 | 0.15 | 0.21 | 0.21 | 0.10 | 0.19 | 0.01 | 0.05 | 0.23 | 0.01 |
| O | 0.15 | 0.00 | 0.00 | 0.00 | 0.01 | 0.15 | 0.05 | 0.03 | 0.02 | 0.01 | 0.02 | 0.02 | 0.15 | 0.02 | 0.02 | 0.13 | 0.09 | 0.06 | 0.01 | 0.01 | 0.01 | 0.04 | 0.01 | 0.06 |
| P | 0.10 | 0.00 | 0.02 | 0.02 | 0.02 | 0.02 | 0.06 | 0.02 | 0.08 | 0.03 | 0.04 | 0.01 | 0.01 | 0.03 | 0.06 | 0.03 | 0.03 | 0.03 | 0.02 | 0.04 | 0.03 | 0.00 | 0.00 | 0.00 |

Patient No. 2

| | 1 | 2 | 3 | 4 | 5 | 6 | 7 | 8 | 9 | 10 | 11 | 12 | 13 | 14 | 15 | 16 | 17 | 18 | 19 | 20 | 21 | 22 | 23 | 24 |
|---|---|---|---|---|---|---|---|---|---|---|---|---|---|---|---|---|---|---|---|---|---|---|---|---|
| A | 0.33 | 0.02 | 0.15 | 0.10 | 0.11 | 0.23 | 0.29 | 0.17 | 0.34 | 0.24 | 0.27 | 0.15 | 0.15 | 0.30 | 0.34 | 0.28 | 0.25 | 0.30 | 0.11 | 0.20 | 0.01 | 0.01 | 0.00 | 0.01 |
| B | 0.01 | 0.00 | 0.01 | 0.01 | 0.00 | 0.04 | 0.00 | 0.01 | 0.03 | 0.00 | 0.01 | 0.00 | 0.00 | 0.00 | 0.00 | 0.00 | 0.00 | 0.00 | 0.00 | 0.00 | 0.00 | 0.01 | 0.00 | 0.00 |
| C | 0.05 | 0.00 | 0.00 | 0.00 | 0.02 | 0.01 | 0.01 | 0.01 | 0.02 | 0.02 | 0.02 | 0.01 | 0.01 | 0.01 | 0.03 | 0.01 | 0.01 | 0.03 | 0.01 | 0.02 | 0.01 | 0.00 | 0.00 | 0.00 |
| D | 0.05 | 0.00 | 0.01 | 0.01 | 0.01 | 0.03 | 0.03 | 0.01 | 0.06 | 0.02 | 0.01 | 0.02 | 0.02 | 0.02 | 0.06 | 0.03 | 0.03 | 0.01 | 0.00 | 0.01 | 0.01 | 0.00 | 0.01 | 0.01 |
| E | 0.08 | 0.00 | 0.01 | 0.01 | 0.01 | 0.05 | 0.07 | 0.02 | 0.05 | 0.02 | 0.03 | 0.02 | 0.03 | 0.04 | 0.06 | 0.04 | 0.02 | 0.06 | 0.01 | 0.03 | 0.00 | 0.00 | 0.00 | 0.00 |
| F | 0.05 | 0.00 | 0.01 | 0.01 | 0.01 | 0.06 | 0.02 | 0.01 | 0.02 | 0.01 | 0.01 | 0.01 | 0.09 | 0.01 | 0.02 | 0.05 | 0.03 | 0.03 | 0.00 | 0.01 | 0.04 | 0.00 | 0.04 | 0.04 |
| G | 0.08 | 0.03 | 0.01 | 0.02 | 0.02 | 0.05 | 0.06 | 0.04 | 0.14 | 0.05 | 0.02 | 0.01 | 0.02 | 0.04 | 0.11 | 0.05 | 0.04 | 0.04 | 0.01 | 0.03 | 0.01 | 0.03 | 0.04 | 0.04 |
| H | 0.15 | 0.01 | 0.10 | 0.14 | 0.24 | 0.28 | 0.37 | 0.22 | 0.49 | 0.21 | 0.19 | 0.28 | 0.27 | 0.38 | 0.34 | 0.31 | 0.33 | 0.35 | 0.14 | 0.20 | 0.10 | 0.01 | 0.01 | 0.06 |
| I | 0.17 | 0.00 | 0.03 | 0.05 | 0.15 | 0.02 | 0.11 | 0.09 | 0.21 | 0.03 | 0.06 | 0.03 | 0.03 | 0.07 | 0.16 | 0.12 | 0.04 | 0.06 | 0.04 | 0.04 | 0.05 | 0.04 | 0.05 | 0.04 |
| J | 0.11 | 0.00 | 0.08 | 0.07 | 0.05 | 0.08 | 0.13 | 0.09 | 0.20 | 0.11 | 0.05 | 0.06 | 0.12 | 0.08 | 0.14 | 0.15 | 0.13 | 0.11 | 0.04 | 0.06 | 0.05 | 0.02 | 0.01 | 0.03 |
| K | 0.06 | 0.00 | 0.00 | 0.00 | 0.01 | 0.06 | 0.03 | 0.01 | 0.04 | 0.01 | 0.01 | 0.01 | 0.07 | 0.01 | 0.03 | 0.04 | 0.03 | 0.06 | 0.00 | 0.02 | 0.10 | 0.00 | 0.01 | 0.00 |
| L | 0.17 | 0.00 | 0.06 | 0.07 | 0.03 | 0.16 | 0.11 | 0.12 | 0.32 | 0.10 | 0.09 | 0.10 | 0.21 | 0.06 | 0.25 | 0.17 | 0.15 | 0.12 | 0.01 | 0.03 | 0.01 | 0.03 | 0.00 | 0.04 |
| M | 0.13 | 0.00 | 0.02 | 0.05 | 0.02 | 0.18 | 0.09 | 0.05 | 0.17 | 0.05 | 0.08 | 0.02 | 0.30 | 0.12 | 0.14 | 0.15 | 0.07 | 0.09 | 0.04 | 0.05 | 0.18 | 0.05 | 0.09 | 0.01 |
| N | 0.23 | 0.00 | 0.08 | 0.12 | 0.21 | 0.16 | 0.16 | 0.18 | 0.33 | 0.21 | 0.23 | 0.14 | 0.07 | 0.29 | 0.20 | 0.28 | 0.20 | 0.25 | 0.10 | 0.19 | 0.01 | 0.06 | 0.20 | 0.02 |
| O | 0.10 | 0.00 | 0.00 | 0.01 | 0.00 | 0.11 | 0.02 | 0.02 | 0.02 | 0.01 | 0.02 | 0.01 | 0.10 | 0.02 | 0.02 | 0.09 | 0.03 | 0.03 | 0.01 | 0.01 | 0.01 | 0.04 | 0.01 | 0.06 |
| P | 0.09 | 0.00 | 0.02 | 0.02 | 0.01 | 0.03 | 0.04 | 0.02 | 0.09 | 0.03 | 0.04 | 0.01 | 0.01 | 0.03 | 0.07 | 0.02 | 0.02 | 0.03 | 0.01 | 0.04 | 0.06 | 0.00 | 0.00 | 0.00 |

Patient No. 3

| | 1 | 2 | 3 | 4 | 5 | 6 | 7 | 8 | 9 | 10 | 11 | 12 | 13 | 14 | 15 | 16 | 17 | 18 | 19 | 20 | 21 | 22 | 23 | 24 |
|---|---|---|---|---|---|---|---|---|---|---|---|---|---|---|---|---|---|---|---|---|---|---|---|---|
| A | 0.47 | 0.04 | 0.25 | 0.26 | 0.25 | 0.37 | 0.42 | 0.28 | 0.57 | 0.37 | 0.34 | 0.21 | 0.24 | 0.44 | 0.51 | 0.42 | 0.31 | 0.34 | 0.20 | 0.32 | 0.03 | 0.01 | 0.00 | 0.01 |
| B | 0.03 | 0.03 | 0.02 | 0.02 | 0.01 | 0.04 | 0.02 | 0.03 | 0.03 | 0.01 | 0.02 | 0.03 | 0.02 | 0.02 | 0.02 | 0.02 | 0.02 | 0.03 | 0.01 | 0.01 | 0.01 | 0.06 | 0.02 | 0.01 |
| C | 0.07 | 0.01 | 0.01 | 0.01 | 0.02 | 0.03 | 0.02 | 0.01 | 0.02 | 0.03 | 0.02 | 0.01 | 0.02 | 0.02 | 0.03 | 0.04 | 0.02 | 0.03 | 0.01 | 0.02 | 0.02 | 0.01 | 0.01 | 0.02 |
| D | 0.09 | 0.01 | 0.05 | 0.06 | 0.03 | 0.10 | 0.06 | 0.03 | 0.13 | 0.07 | 0.04 | 0.09 | 0.09 | 0.07 | 0.12 | 0.10 | 0.11 | 0.06 | 0.03 | 0.06 | 0.03 | 0.03 | 0.03 | 0.03 |
| E | 0.08 | 0.00 | 0.01 | 0.01 | 0.02 | 0.06 | 0.07 | 0.02 | 0.05 | 0.03 | 0.03 | 0.01 | 0.04 | 0.04 | 0.04 | 0.06 | 0.04 | 0.07 | 0.01 | 0.03 | 0.03 | 0.02 | 0.00 | 0.01 |
| F | 0.08 | 0.00 | 0.01 | 0.01 | 0.01 | 0.13 | 0.03 | 0.02 | 0.02 | 0.02 | 0.02 | 0.02 | 0.15 | 0.02 | 0.02 | 0.08 | 0.06 | 0.06 | 0.01 | 0.01 | 0.17 | 0.02 | 0.07 | 0.07 |
| G | 0.16 | 0.01 | 0.03 | 0.05 | 0.06 | 0.09 | 0.13 | 0.08 | 0.19 | 0.09 | 0.12 | 0.06 | 0.07 | 0.12 | 0.19 | 0.10 | 0.09 | 0.10 | 0.04 | 0.10 | 0.06 | 0.18 | 0.11 | 0.05 |
| H | 0.42 | 0.12 | 0.24 | 0.33 | 0.35 | 0.51 | 0.63 | 0.54 | 0.84 | 0.62 | 0.66 | 0.71 | 0.67 | 0.78 | 0.71 | 0.66 | 0.68 | 0.58 | 0.32 | 0.50 | 0.20 | 0.11 | 0.10 | 0.13 |
| I | 0.38 | 0.04 | 0.10 | 0.15 | 0.18 | 0.28 | 0.34 | 0.28 | 0.48 | 0.30 | 0.28 | 0.17 | 0.30 | 0.43 | 0.44 | 0.36 | 0.27 | 0.22 | 0.09 | 0.17 | 0.14 | 0.14 | 0.14 | 0.15 |
| J | 0.32 | 0.06 | 0.16 | 0.23 | 0.20 | 0.44 | 0.35 | 0.21 | 0.39 | 0.29 | 0.32 | 0.37 | 0.47 | 0.37 | 0.43 | 0.35 | 0.40 | 0.32 | 0.11 | 0.21 | 0.15 | 0.12 | 0.06 | 0.10 |
| K | 0.17 | 0.00 | 0.01 | 0.01 | 0.01 | 0.13 | 0.05 | 0.02 | 0.02 | 0.01 | 0.03 | 0.02 | 0.13 | 0.02 | 0.04 | 0.12 | 0.04 | 0.08 | 0.01 | 0.02 | 0.54 | 0.01 | 0.02 | 0.04 |
| L | 0.31 | 0.01 | 0.23 | 0.30 | 0.10 | 0.32 | 0.19 | 0.23 | 0.43 | 0.25 | 0.26 | 0.17 | 0.39 | 0.24 | 0.39 | 0.33 | 0.26 | 0.23 | 0.07 | 0.13 | 0.12 | 0.06 | 0.02 | 0.14 |
| M | 0.26 | 0.01 | 0.03 | 0.03 | 0.05 | 0.30 | 0.08 | 0.04 | 0.13 | 0.08 | 0.10 | 0.03 | 0.30 | 0.05 | 0.16 | 0.20 | 0.10 | 0.11 | 0.03 | 0.09 | 0.79 | 0.29 | 0.26 | 0.02 |
| N | 0.36 | 0.02 | 0.23 | 0.30 | 0.24 | 0.39 | 0.43 | 0.26 | 0.74 | 0.49 | 0.39 | 0.30 | 0.45 | 0.47 | 0.64 | 0.49 | 0.40 | 0.44 | 0.15 | 0.36 | 0.02 | 0.41 | 0.66 | 0.06 |
| O | 0.14 | 0.01 | 0.02 | 0.01 | 0.02 | 0.22 | 0.05 | 0.03 | 0.06 | 0.03 | 0.03 | 0.02 | 0.19 | 0.03 | 0.05 | 0.12 | 0.06 | 0.07 | 0.02 | 0.03 | 0.04 | 0.13 | 0.05 | 0.12 |
| P | 0.12 | 0.02 | 0.05 | 0.05 | 0.08 | 0.04 | 0.07 | 0.04 | 0.16 | 0.10 | 0.10 | 0.04 | 0.02 | 0.08 | 0.14 | 0.08 | 0.06 | 0.06 | 0.06 | 0.11 | 0.05 | -0.01 | 0.00 | 0.00 |

Patient No. 4

| | 1 | 2 | 3 | 4 | 5 | 6 | 7 | 8 | 9 | 10 | 11 | 12 | 13 | 14 | 15 | 16 | 17 | 18 | 19 | 20 | 21 | 22 | 23 | 24 |
|---|---|---|---|---|---|---|---|---|---|---|---|---|---|---|---|---|---|---|---|---|---|---|---|---|
| A | 0.41 | 0.04 | 0.38 | 0.44 | 0.27 | 0.51 | 0.54 | 0.47 | 0.66 | 0.46 | 0.39 | 0.35 | 0.36 | 0.63 | 0.63 | 0.55 | 0.47 | 0.51 | 0.21 | 0.29 | 0.03 | 0.04 | 0.00 | 0.04 |
| B | 0.01 | 0.02 | 0.01 | 0.01 | 0.01 | 0.02 | 0.01 | 0.01 | 0.02 | 0.01 | 0.01 | 0.01 | 0.01 | 0.01 | 0.02 | 0.01 | 0.01 | 0.01 | 0.00 | 0.00 | 0.02 | 0.07 | 0.08 | 0.02 |
| C | 0.16 | 0.01 | 0.01 | 0.02 | 0.03 | 0.08 | 0.04 | 0.02 | 0.04 | 0.05 | 0.05 | 0.02 | 0.07 | 0.03 | 0.06 | 0.08 | 0.05 | 0.07 | 0.02 | 0.04 | 0.06 | 0.02 | 0.02 | 0.04 |
| D | 0.12 | 0.01 | 0.11 | 0.13 | 0.04 | 0.12 | 0.09 | 0.06 | 0.21 | 0.07 | 0.07 | 0.12 | 0.10 | 0.11 | 0.16 | 0.14 | 0.13 | 0.08 | 0.02 | 0.08 | 0.07 | 0.04 | 0.06 | 0.05 |
| E | 0.16 | 0.01 | 0.04 | 0.03 | 0.04 | 0.13 | 0.14 | 0.06 | 0.09 | 0.07 | 0.07 | 0.04 | 0.10 | 0.08 | 0.09 | 0.11 | 0.07 | 0.10 | 0.02 | 0.06 | 0.04 | 0.04 | 0.02 | 0.02 |
| F | 0.11 | 0.00 | 0.01 | 0.01 | 0.01 | 0.13 | 0.05 | 0.02 | 0.04 | 0.02 | 0.02 | 0.04 | 0.23 | 0.03 | 0.04 | 0.11 | 0.09 | 0.07 | 0.01 | 0.02 | 0.16 | 0.01 | 0.18 | 0.20 |
| G | 0.26 | 0.01 | 0.07 | 0.09 | 0.06 | 0.14 | 0.10 | 0.08 | 0.25 | 0.08 | 0.11 | 0.07 | 0.10 | 0.14 | 0.19 | 0.14 | 0.15 | 0.10 | 0.03 | 0.06 | 0.09 | 0.20 | 0.14 | 0.11 |
| H | 0.30 | 0.10 | 0.27 | 0.40 | 0.31 | 0.28 | 0.49 | 0.52 | 0.75 | 0.46 | 0.45 | 0.71 | 0.36 | 0.73 | 0.72 | 0.43 | 0.55 | 0.45 | 0.21 | 0.39 | 0.22 | 0.13 | 0.11 | 0.21 |
| I | 0.27 | 0.02 | 0.15 | 0.22 | 0.07 | 0.26 | 0.16 | 0.29 | 0.42 | 0.20 | 0.14 | 0.21 | 0.22 | 0.31 | 0.28 | 0.21 | 0.22 | 0.21 | 0.05 | 0.11 | 0.14 | 0.19 | 0.18 | 0.16 |
| J | 0.31 | 0.01 | 0.21 | 0.32 | 0.13 | 0.32 | 0.28 | 0.12 | 0.40 | 0.17 | 0.21 | 0.32 | 0.34 | 0.23 | 0.27 | 0.27 | 0.33 | 0.27 | 0.06 | 0.17 | 0.20 | 0.20 | 0.08 | 0.14 |
| K | 0.19 | 0.01 | 0.01 | 0.01 | 0.02 | 0.20 | 0.07 | 0.03 | 0.03 | 0.02 | 0.02 | 0.04 | 0.25 | 0.03 | 0.05 | 0.18 | 0.09 | 0.16 | 0.01 | 0.02 | 0.47 | 0.00 | 0.03 | 0.06 |
| L | 0.26 | 0.01 | 0.16 | 0.21 | 0.07 | 0.26 | 0.10 | 0.24 | 0.47 | 0.23 | 0.19 | 0.12 | 0.32 | 0.20 | 0.31 | 0.21 | 0.29 | 0.19 | 0.05 | 0.13 | 0.12 | 0.12 | 0.04 | 0.21 |
| M | 0.28 | 0.01 | 0.07 | 0.08 | 0.10 | 0.29 | 0.17 | 0.11 | 0.25 | 0.15 | 0.14 | 0.08 | 0.27 | 0.12 | 0.25 | 0.26 | 0.16 | 0.20 | 0.05 | 0.13 | 0.71 | 0.28 | 0.25 | 0.03 |
| N | 0.18 | 0.02 | 0.27 | 0.39 | 0.23 | 0.13 | 0.37 | 0.36 | 0.67 | 0.37 | 0.31 | 0.30 | 0.18 | 0.42 | 0.48 | 0.24 | 0.40 | 0.42 | 0.16 | 0.29 | 0.03 | 0.48 | 0.58 | 0.10 |
| O | 0.17 | 0.01 | 0.03 | 0.03 | 0.03 | 0.23 | 0.11 | 0.06 | 0.09 | 0.04 | 0.04 | 0.04 | 0.31 | 0.06 | 0.06 | 0.19 | 0.14 | 0.12 | 0.02 | 0.04 | 0.06 | 0.19 | 0.07 | 0.17 |
| P | 0.18 | 0.01 | 0.08 | 0.09 | 0.09 | 0.10 | 0.12 | 0.08 | 0.22 | 0.13 | 0.14 | 0.07 | 0.04 | 0.14 | 0.20 | 0.14 | 0.11 | 0.09 | 0.09 | 0.09 | 0.17 | 0.02 | 0.00 | 0.00 |

Patient No. 5

| | 1 | 2 | 3 | 4 | 5 | 6 | 7 | 8 | 9 | 10 | 11 | 12 | 13 | 14 | 15 | 16 | 17 | 18 | 19 | 20 | 21 | 22 | 23 | 24 |
|---|---|---|---|---|---|---|---|---|---|---|---|---|---|---|---|---|---|---|---|---|---|---|---|---|
| A | 0.50 | 0.32 | 0.47 | 0.62 | 0.34 | 0.64 | 0.66 | 0.68 | 0.57 | 0.55 | 0.43 | 0.66 | 0.58 | 0.53 | 0.54 | 0.67 | 0.64 | 0.46 | 0.29 | 0.29 | 0.12 | 0.10 | 0.02 | 0.08 |
| B | 0.18 | 0.06 | 0.19 | 0.22 | 0.13 | 0.22 | 0.17 | 0.25 | 0.19 | 0.15 | 0.17 | 0.14 | 0.13 | 0.13 | 0.16 | 0.11 | 0.15 | 0.13 | 0.05 | 0.11 | 0.10 | 0.09 | 0.10 | 0.05 |
| C | 0.10 | 0.05 | 0.02 | 0.03 | 0.05 | 0.05 | 0.06 | 0.05 | 0.09 | 0.08 | 0.07 | 0.04 | 0.03 | 0.07 | 0.09 | 0.06 | 0.05 | 0.06 | 0.04 | 0.06 | 0.15 | 0.20 | 0.18 | 0.15 |
| D | 0.29 | 0.08 | 0.31 | 0.30 | 0.45 | 0.30 | 0.44 | 0.47 | 0.45 | 0.53 | 0.50 | 0.44 | 0.39 | 0.32 | 0.39 | 0.42 | 0.37 | 0.34 | 0.34 | 0.53 | 0.21 | 0.20 | 0.17 | 0.15 |
| E | 0.15 | 0.04 | 0.08 | 0.06 | 0.09 | 0.14 | 0.12 | 0.11 | 0.17 | 0.14 | 0.12 | 0.07 | 0.09 | 0.12 | 0.15 | 0.13 | 0.10 | 0.14 | 0.07 | 0.10 | 0.17 | 0.16 | 0.14 | 0.19 |
| F | 0.09 | 0.04 | 0.04 | 0.05 | 0.05 | 0.17 | 0.08 | 0.06 | 0.08 | 0.07 | 0.07 | 0.08 | 0.17 | 0.06 | 0.08 | 0.11 | 0.08 | 0.07 | 0.07 | 0.06 | 0.39 | 0.13 | 0.32 | 0.38 |
| G | 0.36 | 0.13 | 0.22 | 0.21 | 0.54 | 0.31 | 0.39 | 0.52 | 0.53 | 0.37 | 0.46 | 0.32 | 0.36 | 0.35 | 0.50 | 0.49 | 0.37 | 0.51 | 0.48 | 0.40 | 0.51 | 0.31 | 0.27 | 0.38 |
| H | 0.39 | 0.48 | 0.47 | 0.69 | 0.31 | 0.63 | 0.78 | 0.84 | 0.70 | 0.51 | 0.48 | 0.77 | 0.84 | 0.66 | 0.66 | 0.57 | 0.92 | 0.75 | 0.33 | 0.36 | 0.36 | 0.42 | 0.38 | 0.46 |
| I | 0.30 | 0.29 | 0.31 | 0.38 | 0.45 | 0.49 | 0.40 | 0.56 | 0.65 | 0.60 | 0.47 | 0.57 | 0.52 | 0.41 | 0.32 | 0.59 | 0.48 | 0.40 | 0.43 | 0.37 | 0.31 | 0.34 | 0.28 | 0.27 |
| J | 0.28 | 0.33 | 0.34 | 0.38 | 0.46 | 0.38 | 0.25 | 0.36 | 0.47 | 0.50 | 0.33 | 0.45 | 0.49 | 0.32 | 0.36 | 0.36 | 0.54 | 0.46 | 0.44 | 0.26 | 0.36 | 0.35 | 0.41 | 0.39 |
| K | 0.13 | 0.04 | 0.05 | 0.04 | 0.04 | 0.14 | 0.08 | 0.03 | 0.08 | 0.06 | 0.06 | 0.05 | 0.15 | 0.06 | 0.10 | 0.13 | 0.06 | 0.11 | 0.06 | 0.06 | 0.35 | 0.18 | 0.05 | 0.07 |
| L | 0.14 | 0.05 | 0.14 | 0.21 | 0.15 | 0.25 | 0.13 | 0.42 | 0.42 | 0.27 | 0.13 | 0.12 | 0.32 | 0.15 | 0.26 | 0.22 | 0.25 | 0.16 | 0.22 | 0.15 | 0.19 | 0.13 | 0.04 | 0.25 |
| M | 0.33 | 0.15 | 0.11 | 0.10 | 0.15 | 0.34 | 0.17 | 0.13 | 0.27 | 0.19 | 0.27 | 0.10 | 0.28 | 0.17 | 0.36 | 0.24 | 0.16 | 0.19 | 0.19 | 0.22 | 0.40 | 0.13 | 0.24 | 0.06 |
| N | 0.26 | 0.26 | 0.39 | 0.45 | 0.18 | 0.34 | 0.46 | 0.62 | 0.55 | 0.30 | 0.25 | 0.48 | 0.53 | 0.47 | 0.40 | 0.49 | 0.53 | 0.53 | 0.28 | 0.14 | 0.06 | 0.38 | 0.29 | 0.14 |
| O | 0.16 | 0.08 | 0.08 | 0.08 | 0.09 | 0.27 | 0.13 | 0.09 | 0.17 | 0.10 | 0.13 | 0.08 | 0.17 | 0.16 | 0.13 | 0.18 | 0.14 | 0.12 | 0.08 | 0.11 | 0.12 | 0.25 | 0.20 | 0.22 |
| P | 0.21 | 0.16 | 0.14 | 0.20 | 0.21 | 0.17 | 0.19 | 0.23 | 0.26 | 0.22 | 0.23 | 0.17 | 0.10 | 0.35 | 0.33 | 0.25 | 0.27 | 0.25 | 0.34 | 0.18 | 0.14 | 0.01 | 0.00 | 0.00 |

**Normal conversion rate**
**3 h**

**Patient No. 1**

| | 1 | 2 | 3 | 4 | 5 | 6 | 7 | 8 | 9 | 10 | 11 | 12 | 13 | 14 | 15 | 16 | 17 | 18 | 19 | 20 | 21 | 22 | 23 | 24 |
|---|---|---|---|---|---|---|---|---|---|---|---|---|---|---|---|---|---|---|---|---|---|---|---|---|
| A | 0.33 | 0.02 | 0.06 | 0.04 | 0.08 | 0.15 | 0.21 | 0.11 | 0.26 | 0.17 | 0.20 | 0.08 | 0.14 | 0.20 | 0.29 | 0.18 | 0.12 | 0.22 | 0.11 | 0.14 | 0.02 | 0.00 | 0.00 | 0.01 |
| B | 0.01 | 0.00 | 0.01 | 0.01 | 0.00 | 0.03 | 0.00 | 0.00 | 0.02 | 0.00 | 0.00 | 0.00 | 0.01 | 0.00 | 0.00 | 0.01 | 0.00 | 0.01 | 0.00 | 0.00 | 0.00 | 0.03 | 0.00 | 0.01 |
| C | 0.06 | 0.00 | 0.00 | 0.00 | 0.01 | 0.01 | 0.01 | 0.00 | 0.00 | 0.01 | 0.01 | 0.00 | 0.01 | 0.00 | 0.01 | 0.02 | 0.01 | 0.02 | 0.00 | 0.01 | 0.01 | 0.00 | 0.00 | 0.00 |
| D | 0.04 | 0.00 | 0.00 | 0.00 | 0.01 | 0.03 | 0.02 | 0.01 | 0.03 | 0.02 | 0.01 | 0.01 | 0.02 | 0.02 | 0.03 | 0.02 | 0.02 | 0.01 | 0.01 | 0.01 | 0.02 | 0.01 | 0.01 | 0.01 |
| E | 0.09 | 0.00 | 0.00 | 0.01 | 0.02 | 0.05 | 0.08 | 0.02 | 0.02 | 0.01 | 0.02 | 0.01 | 0.03 | 0.03 | 0.03 | 0.04 | 0.02 | 0.07 | 0.01 | 0.01 | 0.01 | 0.01 | 0.00 | 0.00 |
| F | 0.14 | 0.00 | 0.00 | 0.01 | 0.01 | 0.11 | 0.04 | 0.01 | 0.01 | 0.01 | 0.01 | 0.01 | 0.11 | 0.01 | 0.01 | 0.08 | 0.06 | 0.07 | 0.00 | 0.00 | 0.03 | 0.00 | 0.01 | 0.02 |
| G | 0.04 | 0.02 | 0.00 | 0.00 | 0.01 | 0.02 | 0.03 | 0.01 | 0.04 | 0.01 | 0.01 | 0.00 | 0.01 | 0.01 | 0.06 | 0.02 | 0.01 | 0.01 | 0.01 | 0.01 | 0.01 | 0.03 | 0.03 | 0.02 |
| H | 0.18 | 0.01 | 0.04 | 0.06 | 0.21 | 0.21 | 0.25 | 0.14 | 0.32 | 0.18 | 0.15 | 0.14 | 0.21 | 0.22 | 0.25 | 0.26 | 0.24 | 0.22 | 0.11 | 0.15 | 0.06 | 0.01 | 0.01 | 0.05 |
| I | 0.12 | 0.00 | 0.01 | 0.02 | 0.10 | 0.02 | 0.05 | 0.04 | 0.18 | 0.03 | 0.07 | 0.03 | 0.05 | 0.05 | 0.13 | 0.08 | 0.03 | 0.04 | 0.03 | 0.05 | 0.05 | 0.02 | 0.03 | 0.03 |
| J | 0.11 | 0.00 | 0.03 | 0.03 | 0.08 | 0.08 | 0.10 | 0.06 | 0.17 | 0.12 | 0.07 | 0.06 | 0.15 | 0.07 | 0.15 | 0.13 | 0.13 | 0.11 | 0.09 | 0.06 | 0.04 | 0.02 | 0.02 | 0.02 |
| K | 0.12 | 0.00 | 0.00 | 0.00 | 0.00 | 0.08 | 0.04 | 0.02 | 0.02 | 0.01 | 0.01 | 0.01 | 0.08 | 0.01 | 0.03 | 0.07 | 0.03 | 0.07 | 0.00 | 0.01 | 0.08 | 0.00 | 0.00 | 0.00 |
| L | 0.22 | 0.00 | 0.08 | 0.09 | 0.04 | 0.17 | 0.16 | 0.10 | 0.22 | 0.08 | 0.09 | 0.11 | 0.20 | 0.06 | 0.24 | 0.16 | 0.12 | 0.13 | 0.03 | 0.04 | 0.00 | 0.02 | 0.00 | 0.02 |
| M | 0.20 | 0.00 | 0.00 | 0.03 | 0.02 | 0.19 | 0.07 | 0.03 | 0.06 | 0.03 | 0.05 | 0.02 | 0.27 | 0.08 | 0.07 | 0.15 | 0.06 | 0.08 | 0.02 | 0.03 | 0.17 | 0.03 | 0.09 | 0.01 |
| N | 0.20 | 0.01 | 0.04 | 0.06 | 0.18 | 0.15 | 0.14 | 0.11 | 0.27 | 0.16 | 0.23 | 0.10 | 0.06 | 0.18 | 0.18 | 0.20 | 0.18 | 0.18 | 0.08 | 0.18 | 0.00 | 0.04 | 0.24 | 0.01 |
| O | 0.22 | 0.00 | 0.00 | 0.00 | 0.00 | 0.15 | 0.03 | 0.02 | 0.01 | 0.01 | 0.02 | 0.01 | 0.13 | 0.01 | 0.01 | 0.13 | 0.07 | 0.05 | 0.01 | 0.01 | 0.01 | 0.02 | 0.01 | 0.06 |
| P | 0.07 | 0.00 | 0.01 | 0.01 | 0.01 | 0.01 | 0.03 | 0.01 | 0.03 | 0.02 | 0.02 | 0.00 | 0.00 | 0.01 | 0.03 | 0.02 | 0.01 | 0.01 | 0.01 | 0.02 | 0.02 | 0.02 | 0.00 | 0.00 |

**Patient No. 2**

| | 1 | 2 | 3 | 4 | 5 | 6 | 7 | 8 | 9 | 10 | 11 | 12 | 13 | 14 | 15 | 16 | 17 | 18 | 19 | 20 | 21 | 22 | 23 | 24 |
|---|---|---|---|---|---|---|---|---|---|---|---|---|---|---|---|---|---|---|---|---|---|---|---|---|
| A | 0.26 | 0.01 | 0.05 | 0.04 | 0.06 | 0.10 | 0.13 | 0.07 | 0.22 | 0.15 | 0.18 | 0.04 | 0.07 | 0.12 | 0.23 | 0.12 | 0.09 | 0.13 | 0.07 | 0.13 | 0.01 | 0.00 | 0.00 | 0.00 |
| B | 0.01 | 0.00 | 0.00 | 0.00 | 0.00 | 0.02 | 0.00 | 0.00 | 0.02 | 0.00 | 0.00 | 0.00 | 0.00 | 0.00 | 0.00 | 0.00 | 0.00 | 0.00 | 0.00 | 0.00 | 0.00 | 0.01 | 0.00 | 0.00 |
| C | 0.03 | 0.00 | 0.00 | 0.00 | 0.01 | 0.01 | 0.01 | 0.00 | 0.00 | 0.01 | 0.01 | 0.00 | 0.00 | 0.00 | 0.01 | 0.01 | 0.00 | 0.01 | 0.00 | 0.01 | 0.00 | 0.00 | 0.00 | 0.00 |
| D | 0.03 | 0.00 | 0.00 | 0.00 | 0.00 | 0.01 | 0.01 | 0.00 | 0.02 | 0.01 | 0.00 | 0.01 | 0.01 | 0.01 | 0.02 | 0.01 | 0.01 | 0.00 | 0.00 | 0.00 | 0.00 | 0.00 | 0.00 | 0.01 |
| E | 0.05 | 0.00 | 0.00 | 0.00 | 0.01 | 0.02 | 0.03 | 0.01 | 0.02 | 0.01 | 0.01 | 0.00 | 0.01 | 0.01 | 0.02 | 0.01 | 0.01 | 0.03 | 0.00 | 0.01 | 0.00 | 0.00 | 0.00 | 0.00 |
| F | 0.07 | 0.00 | 0.00 | 0.00 | 0.00 | 0.05 | 0.01 | 0.00 | 0.01 | 0.00 | 0.00 | 0.01 | 0.04 | 0.01 | 0.01 | 0.04 | 0.02 | 0.02 | 0.00 | 0.00 | 0.02 | 0.00 | 0.01 | 0.01 |
| G | 0.03 | 0.03 | 0.00 | 0.01 | 0.01 | 0.01 | 0.02 | 0.01 | 0.05 | 0.02 | 0.01 | 0.00 | 0.00 | 0.01 | 0.05 | 0.01 | 0.01 | 0.01 | 0.00 | 0.01 | 0.00 | 0.01 | 0.01 | 0.01 |
| H | 0.15 | 0.01 | 0.03 | 0.05 | 0.16 | 0.25 | 0.20 | 0.11 | 0.29 | 0.13 | 0.14 | 0.10 | 0.23 | 0.19 | 0.23 | 0.24 | 0.16 | 0.18 | 0.08 | 0.13 | 0.03 | 0.01 | 0.00 | 0.02 |
| I | 0.12 | 0.00 | 0.01 | 0.02 | 0.08 | 0.01 | 0.04 | 0.04 | 0.16 | 0.02 | 0.04 | 0.01 | 0.02 | 0.03 | 0.10 | 0.05 | 0.01 | 0.03 | 0.01 | 0.02 | 0.02 | 0.01 | 0.02 | 0.02 |
| J | 0.10 | 0.00 | 0.03 | 0.03 | 0.03 | 0.05 | 0.07 | 0.04 | 0.15 | 0.09 | 0.04 | 0.03 | 0.07 | 0.05 | 0.13 | 0.10 | 0.08 | 0.06 | 0.03 | 0.04 | 0.02 | 0.01 | 0.00 | 0.01 |
| K | 0.08 | 0.00 | 0.00 | 0.00 | 0.00 | 0.03 | 0.01 | 0.01 | 0.02 | 0.01 | 0.01 | 0.00 | 0.03 | 0.01 | 0.03 | 0.03 | 0.01 | 0.03 | 0.00 | 0.01 | 0.07 | 0.00 | 0.00 | 0.00 |
| L | 0.17 | 0.00 | 0.05 | 0.06 | 0.02 | 0.13 | 0.10 | 0.07 | 0.20 | 0.06 | 0.06 | 0.08 | 0.14 | 0.04 | 0.20 | 0.13 | 0.09 | 0.10 | 0.01 | 0.02 | 0.00 | 0.01 | 0.00 | 0.02 |
| M | 0.10 | 0.00 | 0.00 | 0.02 | 0.01 | 0.10 | 0.03 | 0.01 | 0.04 | 0.02 | 0.03 | 0.01 | 0.24 | 0.05 | 0.05 | 0.07 | 0.02 | 0.04 | 0.01 | 0.02 | 0.15 | 0.04 | 0.09 | 0.01 |
| N | 0.28 | 0.00 | 0.04 | 0.05 | 0.13 | 0.22 | 0.09 | 0.06 | 0.22 | 0.15 | 0.17 | 0.06 | 0.08 | 0.14 | 0.17 | 0.21 | 0.10 | 0.12 | 0.06 | 0.15 | 0.00 | 0.03 | 0.20 | 0.01 |
| O | 0.12 | 0.00 | 0.00 | 0.00 | 0.00 | 0.08 | 0.01 | 0.01 | 0.01 | 0.00 | 0.01 | 0.01 | 0.06 | 0.01 | 0.01 | 0.06 | 0.02 | 0.02 | 0.00 | 0.01 | 0.01 | 0.02 | 0.00 | 0.04 |
| P | 0.04 | 0.00 | 0.00 | 0.01 | 0.00 | 0.01 | 0.02 | 0.01 | 0.03 | 0.01 | 0.01 | 0.00 | 0.00 | 0.01 | 0.03 | 0.01 | 0.01 | 0.01 | 0.00 | 0.01 | 0.02 | 0.01 | 0.00 | 0.00 |

**Patient No. 3**

| | 1 | 2 | 3 | 4 | 5 | 6 | 7 | 8 | 9 | 10 | 11 | 12 | 13 | 14 | 15 | 16 | 17 | 18 | 19 | 20 | 21 | 22 | 23 | 24 |
|---|---|---|---|---|---|---|---|---|---|---|---|---|---|---|---|---|---|---|---|---|---|---|---|---|
| A | 0.36 | 0.03 | 0.08 | 0.09 | 0.17 | 0.30 | 0.25 | 0.15 | 0.36 | 0.24 | 0.22 | 0.08 | 0.25 | 0.21 | 0.34 | 0.32 | 0.19 | 0.26 | 0.12 | 0.22 | 0.05 | 0.00 | 0.00 | 0.00 |
| B | 0.03 | 0.02 | 0.01 | 0.01 | 0.00 | 0.03 | 0.01 | 0.02 | 0.01 | 0.01 | 0.01 | 0.01 | 0.02 | 0.01 | 0.01 | 0.02 | 0.01 | 0.02 | 0.00 | 0.00 | 0.00 | 0.06 | 0.01 | 0.01 |
| C | 0.13 | 0.00 | 0.00 | 0.00 | 0.01 | 0.06 | 0.02 | 0.01 | 0.01 | 0.02 | 0.02 | 0.01 | 0.04 | 0.01 | 0.02 | 0.07 | 0.03 | 0.04 | 0.01 | 0.01 | 0.01 | 0.01 | 0.01 | 0.01 |
| D | 0.07 | 0.00 | 0.01 | 0.02 | 0.02 | 0.08 | 0.03 | 0.01 | 0.05 | 0.03 | 0.02 | 0.02 | 0.07 | 0.03 | 0.05 | 0.07 | 0.05 | 0.05 | 0.01 | 0.03 | 0.05 | 0.01 | 0.02 | 0.05 |
| E | 0.17 | 0.01 | 0.01 | 0.00 | 0.02 | 0.12 | 0.15 | 0.03 | 0.03 | 0.03 | 0.04 | 0.01 | 0.08 | 0.05 | 0.03 | 0.10 | 0.05 | 0.16 | 0.01 | 0.02 | 0.04 | 0.04 | 0.00 | 0.00 |
| F | 0.20 | 0.00 | 0.00 | 0.00 | 0.01 | 0.19 | 0.08 | 0.03 | 0.01 | 0.01 | 0.03 | 0.03 | 0.24 | 0.02 | 0.01 | 0.17 | 0.13 | 0.15 | 0.00 | 0.01 | 0.14 | 0.01 | 0.02 | 0.02 |
| G | 0.14 | 0.00 | 0.01 | 0.02 | 0.05 | 0.05 | 0.09 | 0.05 | 0.17 | 0.09 | 0.09 | 0.02 | 0.03 | 0.09 | 0.19 | 0.06 | 0.05 | 0.07 | 0.03 | 0.08 | 0.04 | 0.16 | 0.06 | 0.03 |
| H | 0.27 | 0.06 | 0.09 | 0.10 | 0.21 | 0.37 | 0.41 | 0.27 | 0.47 | 0.33 | 0.37 | 0.33 | 0.55 | 0.37 | 0.48 | 0.47 | 0.47 | 0.46 | 0.17 | 0.28 | 0.08 | 0.04 | 0.05 | 0.05 |
| I | 0.30 | 0.02 | 0.03 | 0.05 | 0.12 | 0.25 | 0.15 | 0.11 | 0.27 | 0.21 | 0.21 | 0.07 | 0.31 | 0.19 | 0.33 | 0.26 | 0.14 | 0.11 | 0.08 | 0.14 | 0.15 | 0.05 | 0.07 | 0.13 |
| J | 0.28 | 0.02 | 0.05 | 0.08 | 0.18 | 0.40 | 0.25 | 0.13 | 0.27 | 0.19 | 0.25 | 0.19 | 0.44 | 0.24 | 0.32 | 0.29 | 0.28 | 0.22 | 0.10 | 0.17 | 0.10 | 0.11 | 0.02 | 0.05 |
| K | 0.27 | 0.00 | 0.00 | 0.00 | 0.01 | 0.23 | 0.09 | 0.04 | 0.01 | 0.01 | 0.04 | 0.03 | 0.25 | 0.03 | 0.03 | 0.22 | 0.10 | 0.15 | 0.01 | 0.01 | 0.41 | 0.00 | 0.01 | 0.03 |
| L | 0.32 | 0.01 | 0.21 | 0.26 | 0.08 | 0.27 | 0.16 | 0.18 | 0.37 | 0.19 | 0.24 | 0.16 | 0.36 | 0.24 | 0.34 | 0.30 | 0.20 | 0.24 | 0.06 | 0.08 | 0.05 | 0.04 | 0.01 | 0.16 |
| M | 0.36 | 0.01 | 0.01 | 0.01 | 0.04 | 0.44 | 0.12 | 0.07 | 0.06 | 0.06 | 0.11 | 0.04 | 0.51 | 0.05 | 0.11 | 0.39 | 0.21 | 0.17 | 0.02 | 0.06 | 0.50 | 0.24 | 0.25 | 0.01 |
| N | 0.27 | 0.01 | 0.10 | 0.13 | 0.18 | 0.33 | 0.32 | 0.17 | 0.43 | 0.37 | 0.33 | 0.25 | 0.41 | 0.32 | 0.51 | 0.47 | 0.40 | 0.39 | 0.09 | 0.28 | 0.01 | 0.20 | 0.56 | 0.03 |
| O | 0.34 | 0.00 | 0.00 | 0.00 | 0.01 | 0.38 | 0.11 | 0.05 | 0.02 | 0.02 | 0.02 | 0.02 | 0.33 | 0.02 | 0.02 | 0.27 | 0.12 | 0.17 | 0.01 | 0.01 | 0.02 | 0.05 | 0.03 | 0.06 |
| P | 0.07 | 0.01 | 0.01 | 0.01 | 0.02 | 0.02 | 0.07 | 0.01 | 0.04 | 0.03 | 0.03 | 0.01 | 0.01 | 0.02 | 0.04 | 0.04 | 0.02 | 0.04 | 0.02 | 0.03 | 0.06 | 0.01 | 0.00 | 0.00 |

**Patient No. 4**

| | 1 | 2 | 3 | 4 | 5 | 6 | 7 | 8 | 9 | 10 | 11 | 12 | 13 | 14 | 15 | 16 | 17 | 18 | 19 | 20 | 21 | 22 | 23 | 24 |
|---|---|---|---|---|---|---|---|---|---|---|---|---|---|---|---|---|---|---|---|---|---|---|---|---|
| A | 0.45 | 0.03 | 0.12 | 0.14 | 0.22 | 0.36 | 0.37 | 0.21 | 0.51 | 0.35 | 0.29 | 0.12 | 0.25 | 0.34 | 0.54 | 0.41 | 0.24 | 0.35 | 0.16 | 0.24 | 0.03 | 0.01 | 0.00 | 0.01 |
| B | 0.01 | 0.02 | 0.00 | 0.00 | 0.00 | 0.02 | 0.01 | 0.01 | 0.01 | 0.00 | 0.00 | 0.00 | 0.01 | 0.00 | 0.01 | 0.00 | 0.00 | 0.01 | 0.00 | 0.00 | 0.01 | 0.05 | 0.02 | 0.01 |
| C | 0.15 | 0.01 | 0.00 | 0.00 | 0.01 | 0.05 | 0.02 | 0.01 | 0.01 | 0.03 | 0.03 | 0.01 | 0.05 | 0.02 | 0.02 | 0.07 | 0.03 | 0.05 | 0.01 | 0.02 | 0.02 | 0.01 | 0.01 | 0.01 |
| D | 0.09 | 0.00 | 0.02 | 0.02 | 0.01 | 0.08 | 0.04 | 0.02 | 0.08 | 0.03 | 0.04 | 0.04 | 0.06 | 0.04 | 0.08 | 0.09 | 0.07 | 0.05 | 0.01 | 0.04 | 0.05 | 0.01 | 0.02 | 0.04 |
| E | 0.18 | 0.00 | 0.01 | 0.01 | 0.02 | 0.09 | 0.13 | 0.03 | 0.04 | 0.04 | 0.04 | 0.02 | 0.07 | 0.05 | 0.04 | 0.08 | 0.04 | 0.13 | 0.01 | 0.03 | 0.02 | 0.03 | 0.01 | 0.01 |
| F | 0.27 | 0.00 | 0.01 | 0.00 | 0.01 | 0.15 | 0.06 | 0.02 | 0.01 | 0.01 | 0.01 | 0.02 | 0.23 | 0.02 | 0.01 | 0.14 | 0.09 | 0.10 | 0.01 | 0.01 | 0.11 | 0.00 | 0.03 | 0.04 |
| G | 0.16 | 0.00 | 0.01 | 0.02 | 0.03 | 0.05 | 0.04 | 0.04 | 0.15 | 0.05 | 0.07 | 0.02 | 0.03 | 0.07 | 0.11 | 0.06 | 0.06 | 0.04 | 0.02 | 0.04 | 0.04 | 0.15 | 0.07 | 0.03 |
| H | 0.37 | 0.08 | 0.09 | 0.12 | 0.23 | 0.41 | 0.38 | 0.29 | 0.51 | 0.31 | 0.32 | 0.32 | 0.54 | 0.48 | 0.51 | 0.52 | 0.42 | 0.37 | 0.15 | 0.29 | 0.09 | 0.05 | 0.04 | 0.07 |
| I | 0.26 | 0.01 | 0.04 | 0.06 | 0.06 | 0.24 | 0.09 | 0.14 | 0.33 | 0.19 | 0.12 | 0.08 | 0.18 | 0.18 | 0.23 | 0.17 | 0.12 | 0.12 | 0.04 | 0.10 | 0.11 | 0.07 | 0.08 | 0.10 |
| J | 0.37 | 0.01 | 0.06 | 0.09 | 0.11 | 0.34 | 0.23 | 0.07 | 0.32 | 0.16 | 0.20 | 0.16 | 0.32 | 0.17 | 0.24 | 0.24 | 0.25 | 0.21 | 0.05 | 0.14 | 0.10 | 0.13 | 0.03 | 0.06 |
| K | 0.29 | 0.01 | 0.00 | 0.00 | 0.01 | 0.17 | 0.08 | 0.02 | 0.01 | 0.02 | 0.02 | 0.02 | 0.23 | 0.03 | 0.03 | 0.17 | 0.09 | 0.14 | 0.01 | 0.01 | 0.44 | 0.00 | 0.01 | 0.02 |
| L | 0.29 | 0.00 | 0.14 | 0.17 | 0.05 | 0.19 | 0.09 | 0.16 | 0.35 | 0.14 | 0.18 | 0.09 | 0.22 | 0.18 | 0.28 | 0.17 | 0.19 | 0.15 | 0.03 | 0.09 | 0.05 | 0.05 | 0.02 | 0.13 |
| M | 0.38 | 0.01 | 0.01 | 0.01 | 0.05 | 0.32 | 0.10 | 0.07 | 0.08 | 0.07 | 0.07 | 0.04 | 0.38 | 0.04 | 0.10 | 0.26 | 0.13 | 0.17 | 0.02 | 0.06 | 0.56 | 0.26 | 0.28 | 0.02 |
| N | 0.35 | 0.02 | 0.11 | 0.14 | 0.23 | 0.30 | 0.34 | 0.19 | 0.48 | 0.34 | 0.27 | 0.17 | 0.34 | 0.33 | 0.46 | 0.34 | 0.39 | 0.39 | 0.14 | 0.29 | 0.01 | 0.11 | 0.64 | 0.04 |
| O | 0.35 | 0.01 | 0.01 | 0.00 | 0.01 | 0.36 | 0.09 | 0.04 | 0.02 | 0.02 | 0.03 | 0.02 | 0.30 | 0.03 | 0.02 | 0.23 | 0.13 | 0.15 | 0.01 | 0.02 | 0.02 | 0.06 | 0.03 | 0.08 |
| P | 0.08 | 0.00 | 0.01 | 0.01 | 0.02 | 0.03 | 0.06 | 0.01 | 0.05 | 0.04 | 0.04 | 0.01 | 0.01 | 0.04 | 0.06 | 0.04 | 0.02 | 0.03 | 0.02 | 0.03 | 0.02 | 0.02 | 0.00 | 0.00 |

**Patient No. 5**

| | 1 | 2 | 3 | 4 | 5 | 6 | 7 | 8 | 9 | 10 | 11 | 12 | 13 | 14 | 15 | 16 | 17 | 18 | 19 | 20 | 21 | 22 | 23 | 24 |
|---|---|---|---|---|---|---|---|---|---|---|---|---|---|---|---|---|---|---|---|---|---|---|---|---|
| A | 0.38 | 0.06 | 0.08 | 0.11 | 0.60 | 0.31 | 0.33 | 0.38 | 0.41 | 0.55 | 0.54 | 0.11 | 0.24 | 0.21 | 0.49 | 0.28 | 0.21 | 0.31 | 0.53 | 0.46 | 0.23 | 0.04 | 0.01 | 0.04 |
| B | 0.09 | 0.02 | 0.02 | 0.02 | 0.02 | 0.11 | 0.03 | 0.06 | 0.05 | 0.04 | 0.06 | 0.04 | 0.10 | 0.04 | 0.04 | 0.08 | 0.07 | 0.08 | 0.01 | 0.03 | 0.05 | 0.24 | 0.10 | 0.06 |
| C | 0.08 | 0.01 | 0.00 | 0.00 | 0.03 | 0.03 | 0.03 | 0.02 | 0.02 | 0.04 | 0.03 | 0.01 | 0.03 | 0.02 | 0.02 | 0.03 | 0.02 | 0.03 | 0.03 | 0.02 | 0.09 | 0.10 | 0.10 | 0.05 |
| D | 0.38 | 0.02 | 0.08 | 0.10 | 0.18 | 0.35 | 0.27 | 0.12 | 0.20 | 0.21 | 0.28 | 0.17 | 0.33 | 0.16 | 0.27 | 0.36 | 0.22 | 0.25 | 0.12 | 0.24 | 0.28 | 0.08 | 0.13 | 0.16 |
| E | 0.14 | 0.01 | 0.01 | 0.01 | 0.08 | 0.08 | 0.11 | 0.05 | 0.03 | 0.07 | 0.10 | 0.02 | 0.07 | 0.05 | 0.04 | 0.06 | 0.03 | 0.17 | 0.06 | 0.05 | 0.15 | 0.16 | 0.03 | 0.08 |
| F | 0.23 | 0.01 | 0.01 | 0.01 | 0.03 | 0.20 | 0.08 | 0.05 | 0.02 | 0.04 | 0.07 | 0.03 | 0.25 | 0.02 | 0.02 | 0.18 | 0.09 | 0.14 | 0.03 | 0.03 | 0.38 | 0.01 | 0.06 | 0.07 |
| G | 0.44 | 0.03 | 0.09 | 0.08 | 0.41 | 0.17 | 0.22 | 0.24 | 0.44 | 0.47 | 0.45 | 0.11 | 0.10 | 0.25 | 0.61 | 0.28 | 0.18 | 0.24 | 0.34 | 0.45 | 0.22 | 0.30 | 0.18 | 0.14 |
| H | 0.36 | 0.09 | 0.08 | 0.09 | 0.39 | 0.57 | 0.47 | 0.36 | 0.39 | 0.46 | 0.46 | 0.25 | 0.72 | 0.31 | 0.42 | 0.45 | 0.46 | 0.41 | 0.42 | 0.38 | 0.15 | 0.23 | 0.27 | 0.14 |
| I | 0.76 | 0.06 | 0.14 | 0.15 | 0.65 | 0.50 | 0.37 | 0.38 | 0.67 | 0.92 | 0.85 | 0.23 | 0.44 | 0.37 | 0.74 | 0.60 | 0.39 | 0.36 | 0.48 | 0.71 | 0.30 | 0.12 | 0.20 | 0.21 |
| J | 0.69 | 0.05 | 0.15 | 0.18 | 0.84 | 0.61 | 0.47 | 0.44 | 0.73 | 1.00 | 0.95 | 0.32 | 0.68 | 0.51 | 0.81 | 0.70 | 0.61 | 0.55 | 0.64 | 0.87 | 0.20 | 0.24 | 0.16 | 0.22 |
| K | 0.14 | 0.01 | 0.01 | 0.00 | 0.03 | 0.12 | 0.09 | 0.03 | 0.01 | 0.03 | 0.04 | 0.02 | 0.15 | 0.02 | 0.04 | 0.10 | 0.04 | 0.12 | 0.02 | 0.02 | 0.32 | 0.03 | 0.01 | 0.03 |
| L | 0.45 | 0.04 | 0.25 | 0.37 | 0.52 | 0.42 | 0.41 | 0.67 | 0.38 | 0.60 | 0.53 | 0.22 | 0.47 | 0.36 | 0.41 | 0.44 | 0.33 | 0.48 | 0.42 | 0.53 | 0.26 | 0.04 | 0.01 | 0.48 |
| M | 0.31 | 0.02 | 0.01 | 0.01 | 0.14 | 0.24 | 0.07 | 0.10 | 0.06 | 0.13 | 0.16 | 0.03 | 0.30 | 0.04 | 0.12 | 0.18 | 0.09 | 0.16 | 0.12 | 0.12 | 0.33 | 0.53 | 0.62 | 0.01 |
| N | 0.35 | 0.03 | 0.10 | 0.12 | 0.59 | 0.42 | 0.32 | 0.35 | 0.35 | 0.62 | 0.59 | 0.16 | 0.63 | 0.32 | 0.50 | 0.53 | 0.34 | 0.41 | 0.50 | 0.42 | 0.01 | 0.16 | 0.54 | 0.03 |
| O | 0.24 | 0.01 | 0.00 | 0.00 | 0.03 | 0.22 | 0.06 | 0.04 | 0.01 | 0.03 | 0.06 | 0.01 | 0.18 | 0.02 | 0.01 | 0.15 | 0.07 | 0.11 | 0.02 | 0.02 | 0.01 | 0.05 | 0.18 | 0.21 |
| P | 0.05 | 0.01 | 0.01 | 0.01 | 0.07 | 0.02 | 0.05 | 0.02 | 0.05 | 0.07 | 0.09 | 0.01 | 0.01 | 0.03 | 0.07 | 0.03 | 0.02 | 0.04 | 0.06 | 0.06 | 0.05 | 0.01 | 0.00 | 0.00 |

**T/N 3 h**

| Patient | | 1 | 2 | 3 | 4 | 5 | 6 | 7 | 8 | 9 | 10 | 11 | 12 | 13 | 14 | 15 | 16 | 17 | 18 | 19 | 20 | 21 | 22 | 23 | 24 |
|---|---|---|---|---|---|---|---|---|---|---|---|---|---|---|---|---|---|---|---|---|---|---|---|---|---|
| No. 1 | A | 0.95 | 1.23 | 2.07 | 2.17 | 1.34 | 1.37 | 1.64 | 1.64 | 1.27 | 1.34 | 1.26 | 2.07 | 1.64 | 1.59 | 1.22 | 1.38 | 1.78 | 1.42 | 1.28 | 1.30 | 0.97 | 2.74 | 1.16 | 2.05 |
| | B | 1.16 | 1.70 | 2.02 | 2.01 | 1.34 | 1.39 | 2.20 | 1.58 | 1.33 | 1.45 | 1.94 | 2.57 | 1.96 | 1.92 | 1.71 | 1.82 | 1.61 | 1.53 | 1.93 | 1.65 | 1.47 | 1.39 | 1.76 | 1.65 |
| | C | 1.88 | 2.09 | 3.64 | 3.52 | 1.92 | 2.02 | 2.23 | 2.27 | 2.49 | 1.72 | 1.71 | 2.22 | 2.10 | 2.17 | 1.97 | 1.82 | 2.09 | 1.96 | 2.09 | 1.87 | 1.65 | 1.26 | 1.51 | 1.58 |
| | D | 1.06 | 1.79 | 3.07 | 2.75 | 1.66 | 1.41 | 1.83 | 1.86 | 1.79 | 2.05 | 1.61 | 2.10 | 1.52 | 1.81 | 1.61 | 1.49 | 1.79 | 1.43 | 1.64 | 1.56 | 1.49 | 1.66 | 1.46 | 1.16 |
| | E | 1.09 | 1.00 | 2.18 | 1.74 | 1.31 | 1.80 | 1.46 | 1.46 | 1.47 | 1.28 | 1.18 | 1.79 | 1.86 | 1.64 | 1.35 | 1.63 | 1.90 | 1.23 | 1.24 | 1.14 | 1.35 | 1.33 | 0.99 | 1.20 |
| | F | 0.73 | 3.29 | 1.65 | 1.83 | 1.38 | 1.27 | 1.36 | 1.51 | 1.69 | 1.42 | 1.24 | 2.08 | 1.50 | 1.60 | 1.71 | 1.38 | 1.61 | 1.45 | 1.41 | 1.56 | 1.08 | 5.02 | 3.64 | 2.38 |
| | G | 1.55 | 0.77 | 2.48 | 3.24 | 1.62 | 2.39 | 2.22 | 2.28 | 2.07 | 1.48 | 1.85 | 3.04 | 2.45 | 2.22 | 1.77 | 2.03 | 2.27 | 2.00 | 2.02 | 1.80 | 1.37 | 1.08 | 1.40 | 1.74 |
| | H | 0.56 | 1.16 | 2.25 | 2.24 | 1.13 | 0.46 | 1.31 | 1.47 | 1.35 | 1.33 | 1.27 | 1.79 | 0.69 | 1.51 | 1.36 | 1.00 | 1.33 | 1.24 | 1.33 | 1.29 | 1.59 | 1.45 | 1.55 | 1.86 |
| | I | 0.89 | 0.85 | 2.14 | 2.33 | 1.24 | 1.14 | 1.67 | 1.46 | 1.19 | 0.85 | 1.17 | 1.84 | 1.35 | 1.48 | 1.05 | 1.28 | 1.57 | 1.36 | 1.23 | 1.18 | 1.20 | 1.55 | 1.32 | 1.01 |
| | J | 0.80 | 2.03 | 2.11 | 2.29 | 1.17 | 1.04 | 1.35 | 1.44 | 1.14 | 1.18 | 1.10 | 1.66 | 1.03 | 1.32 | 1.09 | 1.07 | 1.44 | 1.22 | 1.18 | 1.12 | 1.52 | 1.39 | 1.67 | 1.36 |
| | K | 1.05 | 1.25 | 1.57 | 1.50 | 1.52 | 1.41 | 1.86 | 1.80 | 1.20 | 1.30 | 1.09 | 2.01 | 1.53 | 1.53 | 0.95 | 1.34 | 1.81 | 1.54 | 1.19 | 1.22 | 1.08 | 1.50 | 1.83 | 1.64 |
| | L | 0.92 | 1.20 | 1.29 | 1.31 | 1.20 | 1.25 | 1.20 | 1.29 | 1.22 | 1.26 | 1.23 | 1.36 | 1.22 | 1.25 | 0.99 | 1.24 | 1.42 | 1.18 | 1.45 | 1.34 | 1.48 | 1.28 | 1.62 | 1.61 |
| | M | 0.71 | 1.88 | 2.31 | 1.42 | 1.30 | 0.97 | 1.74 | 1.80 | 1.58 | 1.22 | 1.27 | 1.51 | 0.87 | 1.24 | 1.20 | 1.14 | 1.49 | 1.43 | 1.33 | 1.21 | 1.07 | 0.93 | 0.94 | 1.17 |
| | N | 0.31 | 1.19 | 1.94 | 1.90 | 1.12 | 0.24 | 1.10 | 1.61 | 1.23 | 1.07 | 1.14 | 1.41 | 0.52 | 1.26 | 1.04 | 0.75 | 1.16 | 1.15 | 1.19 | 1.09 | 1.97 | 1.28 | 0.95 | 1.20 |
| | O | 0.66 | 0.77 | 1.99 | 2.62 | 1.55 | 1.01 | 1.51 | 1.21 | 1.81 | 1.46 | 1.12 | 1.83 | 1.17 | 1.75 | 1.64 | 1.08 | 1.23 | 1.10 | 1.66 | 1.84 | 1.40 | 1.59 | 1.28 | 1.00 |
| | P | 1.52 | 1.77 | 2.83 | 2.95 | 1.87 | 2.06 | 1.92 | 2.98 | 2.26 | 1.95 | 2.00 | 2.80 | 2.16 | 2.74 | 1.94 | 1.89 | 2.55 | 1.98 | 2.27 | 2.12 | 1.19 | | | |

| Patient | | 1 | 2 | 3 | 4 | 5 | 6 | 7 | 8 | 9 | 10 | 11 | 12 | 13 | 14 | 15 | 16 | 17 | 18 | 19 | 20 | 21 | 22 | 23 | 24 |
|---|---|---|---|---|---|---|---|---|---|---|---|---|---|---|---|---|---|---|---|---|---|---|---|---|---|
| No. 2 | A | 1.27 | 1.60 | 2.90 | 2.84 | 1.74 | 2.40 | 2.24 | 2.31 | 1.57 | 1.63 | 1.55 | 3.54 | 2.30 | 2.54 | 1.43 | 2.42 | 2.78 | 2.22 | 1.62 | 1.58 | 1.60 | 3.11 | 0.17 | 2.69 |
| | B | 1.20 | 1.64 | 2.41 | 2.94 | 3.13 | 2.36 | 1.91 | 1.90 | 1.70 | 4.77 | 2.35 | 1.84 | 1.80 | 1.55 | 2.34 | 2.11 | 2.41 | 1.96 | 3.06 | 2.22 | 3.17 | 1.81 | 2.09 | 2.69 |
| | C | 1.76 | 1.95 | 2.83 | 3.05 | 2.56 | 2.14 | 2.51 | 3.00 | 3.75 | 1.75 | 1.98 | 3.45 | 2.62 | 2.77 | 2.79 | 2.09 | 2.53 | 2.37 | 2.36 | 2.44 | 2.66 | 2.34 | 2.72 | 4.54 |
| | D | 1.75 | 3.26 | 4.42 | 4.94 | 1.80 | 2.22 | 3.18 | 4.05 | 2.78 | 2.03 | 2.59 | 3.58 | 2.83 | 2.79 | 2.92 | 2.40 | 3.34 | 3.16 | 2.53 | 2.71 | 2.11 | 2.69 | 2.64 | 2.01 |
| | E | 1.52 | 4.20 | 3.90 | 4.11 | 2.27 | 2.54 | 2.40 | 3.31 | 3.39 | 2.82 | 2.40 | 3.19 | 2.46 | 3.41 | 3.15 | 2.53 | 3.67 | 2.15 | 3.30 | 3.01 | 2.64 | 1.97 | 1.26 | 2.49 |
| | F | 0.71 | 1.57 | 1.78 | 2.48 | 2.04 | 1.26 | 1.76 | 2.50 | 2.59 | 2.59 | 2.37 | 2.36 | 2.10 | 2.16 | 2.56 | 1.31 | 2.03 | 1.29 | 1.65 | 3.26 | 2.19 | 1.09 | 3.73 | 4.10 |
| | G | 2.37 | 1.13 | 3.75 | 4.42 | 2.30 | 5.03 | 3.57 | 3.44 | 2.96 | 2.75 | 2.82 | 3.80 | 4.82 | 3.32 | 2.44 | 4.30 | 3.75 | 3.70 | 3.27 | 2.82 | 1.97 | 1.89 | 2.77 | 4.02 |
| | H | 1.03 | 1.50 | 3.02 | 2.91 | 1.49 | 1.09 | 1.87 | 2.09 | 1.68 | 1.61 | 1.43 | 2.75 | 1.20 | 1.99 | 1.48 | 1.29 | 2.02 | 1.94 | 1.80 | 1.58 | 2.80 | 2.52 | 2.64 | 2.79 |
| | I | 1.36 | 1.45 | 2.41 | 2.93 | 1.83 | 1.88 | 2.84 | 2.60 | 1.34 | 1.52 | 1.28 | 3.02 | 2.03 | 2.15 | 1.63 | 2.43 | 2.81 | 2.38 | 3.46 | 1.64 | 2.39 | 3.10 | 2.39 | 2.24 |
| | J | 1.10 | 2.72 | 2.77 | 2.73 | 1.56 | 1.67 | 1.93 | 2.35 | 1.31 | 1.29 | 1.30 | 2.11 | 1.66 | 1.49 | 1.13 | 1.54 | 1.65 | 1.77 | 1.42 | 1.46 | 2.73 | 2.52 | 2.70 | 2.27 |
| | K | 0.83 | 2.08 | 1.97 | 1.61 | 1.75 | 1.71 | 1.90 | 2.15 | 1.92 | 1.70 | 1.53 | 1.92 | 2.20 | 2.10 | 1.28 | 1.54 | 2.19 | 2.14 | 2.15 | 2.07 | 1.37 | 1.26 | 2.19 | 1.86 |
| | L | 1.02 | 1.32 | 1.12 | 1.17 | 1.45 | 1.26 | 1.15 | 1.77 | 1.56 | 1.70 | 1.45 | 1.26 | 1.44 | 1.36 | 1.28 | 1.30 | 1.71 | 1.24 | 1.33 | 1.54 | 1.65 | 2.50 | 1.69 | 2.06 |
| | M | 1.27 | 1.50 | 3.92 | 2.12 | 1.52 | 1.78 | 3.39 | 3.84 | 3.85 | 2.62 | 2.62 | 3.63 | 1.23 | 2.12 | 2.58 | 2.06 | 2.81 | 2.40 | 3.22 | 2.57 | 1.23 | 1.18 | 1.02 | 1.93 |
| | N | 0.83 | 1.52 | 2.19 | 2.41 | 1.57 | 0.76 | 1.76 | 2.82 | 1.52 | 1.40 | 1.33 | 2.31 | 0.97 | 2.16 | 1.22 | 1.29 | 2.03 | 2.05 | 1.66 | 1.31 | 2.71 | 2.21 | 1.03 | 2.12 |
| | O | 0.85 | 0.87 | 3.15 | 3.22 | 1.97 | 1.34 | 1.97 | 1.70 | 2.28 | 1.99 | 1.66 | 2.03 | 1.55 | 2.50 | 2.17 | 1.36 | 1.65 | 1.30 | 2.21 | 2.61 | 2.08 | 2.14 | 2.02 | 1.62 |
| | P | 2.51 | 1.59 | 3.36 | 3.92 | 2.61 | 4.38 | 2.77 | 3.54 | 3.11 | 2.75 | 2.85 | 3.63 | 2.24 | 4.11 | 2.73 | 3.65 | 3.64 | 3.86 | 2.85 | 2.75 | 2.73 | | | |

| Patient | | 1 | 2 | 3 | 4 | 5 | 6 | 7 | 8 | 9 | 10 | 11 | 12 | 13 | 14 | 15 | 16 | 17 | 18 | 19 | 20 | 21 | 22 | 23 | 24 |
|---|---|---|---|---|---|---|---|---|---|---|---|---|---|---|---|---|---|---|---|---|---|---|---|---|---|
| No. 3 | A | 1.30 | 1.39 | 3.23 | 2.99 | 1.46 | 1.25 | 1.66 | 1.79 | 1.61 | 1.57 | 1.54 | 2.54 | 0.97 | 2.11 | 1.49 | 1.32 | 1.64 | 1.32 | 1.63 | 1.42 | 0.69 | 2.79 | 2.04 | 2.94 |
| | B | 0.85 | 1.28 | 2.95 | 3.78 | 3.02 | 1.11 | 1.62 | 1.86 | 2.52 | 2.45 | 2.33 | 2.61 | 1.00 | 2.39 | 2.29 | 1.07 | 1.33 | 1.13 | 5.15 | 3.54 | 1.65 | 0.94 | 1.39 | 0.72 |
| | C | 0.56 | 1.78 | 2.38 | 2.40 | 1.70 | 0.54 | 0.70 | 1.02 | 2.52 | 1.49 | 1.15 | 1.25 | 0.49 | 1.69 | 1.71 | 0.55 | 0.56 | 0.65 | 1.81 | 1.73 | 2.19 | 1.63 | 1.54 | 1.96 |
| | D | 1.27 | 2.33 | 4.34 | 4.01 | 2.08 | 1.31 | 2.31 | 2.88 | 2.52 | 2.42 | 2.49 | 3.89 | 1.33 | 2.65 | 2.58 | 1.38 | 2.07 | 1.25 | 2.55 | 2.24 | 0.68 | 2.30 | 2.04 | 0.63 |
| | E | 0.48 | 0.78 | 2.48 | 2.44 | 1.15 | 0.56 | 0.48 | 0.97 | 1.73 | 1.24 | 0.83 | 1.26 | 0.56 | 0.78 | 1.65 | 0.57 | 0.79 | 0.44 | 1.44 | 1.35 | 0.78 | 0.56 | 2.38 | 1.98 |
| | F | 0.39 | 1.12 | 1.56 | 2.15 | 1.41 | 0.69 | 0.43 | 0.69 | 2.24 | 1.19 | 0.58 | 0.75 | 0.62 | 0.81 | 2.28 | 0.47 | 0.50 | 0.39 | 2.53 | 1.63 | 1.27 | 2.71 | 4.75 | 3.90 |
| | G | 1.12 | 1.89 | 2.35 | 2.21 | 1.18 | 1.82 | 1.44 | 1.56 | 1.11 | 1.04 | 1.37 | 2.72 | 2.23 | 1.46 | 1.05 | 1.65 | 1.89 | 1.40 | 1.48 | 1.18 | 1.77 | 1.15 | 1.80 | 1.57 |
| | H | 1.57 | 1.96 | 2.78 | 3.19 | 1.62 | 1.37 | 1.56 | 1.99 | 1.81 | 1.89 | 1.76 | 2.15 | 1.21 | 2.09 | 1.49 | 1.41 | 1.47 | 1.26 | 1.86 | 1.77 | 2.33 | 2.54 | 2.04 | 2.81 |
| | I | 1.25 | 2.24 | 3.08 | 2.86 | 1.45 | 1.10 | 2.32 | 2.54 | 1.77 | 1.40 | 1.30 | 2.55 | 0.95 | 2.23 | 1.33 | 1.36 | 1.86 | 1.95 | 1.02 | 1.22 | 0.91 | 2.56 | 1.98 | 1.14 |
| | J | 1.12 | 3.26 | 3.10 | 2.74 | 1.12 | 1.08 | 1.40 | 1.55 | 1.45 | 1.51 | 1.27 | 1.97 | 1.06 | 1.54 | 1.37 | 1.18 | 1.41 | 1.47 | 1.13 | 1.24 | 1.41 | 1.03 | 2.40 | 2.15 |
| | K | 0.61 | 1.17 | 2.10 | 2.50 | 1.38 | 0.55 | 0.55 | 0.49 | 2.42 | 1.15 | 0.62 | 0.72 | 0.53 | 0.71 | 1.44 | 0.54 | 0.45 | 0.52 | 1.98 | 1.39 | 1.31 | 2.14 | 2.14 | 1.27 |
| | L | 0.96 | 1.22 | 1.11 | 1.15 | 1.23 | 1.17 | 1.17 | 1.27 | 1.16 | 1.30 | 1.08 | 1.12 | 1.08 | 1.00 | 1.13 | 1.09 | 1.35 | 0.99 | 1.19 | 1.54 | 2.40 | 1.49 | 2.42 | 0.88 |
| | M | 0.73 | 1.17 | 2.51 | 2.67 | 1.48 | 0.68 | 0.65 | 0.61 | 2.21 | 1.26 | 0.96 | 0.84 | 0.60 | 1.11 | 1.41 | 0.52 | 0.47 | 0.61 | 1.65 | 1.45 | 1.59 | 1.19 | 1.04 | 1.77 |
| | N | 1.36 | 1.49 | 2.38 | 2.26 | 1.36 | 1.19 | 1.34 | 1.54 | 1.72 | 1.33 | 1.21 | 1.20 | 1.10 | 1.47 | 1.25 | 1.04 | 0.98 | 1.11 | 1.61 | 1.27 | 3.84 | 2.06 | 1.18 | 1.91 |
| | O | 0.42 | 1.73 | 3.85 | 5.02 | 1.95 | 0.59 | 0.46 | 0.63 | 3.59 | 1.77 | 1.15 | 0.92 | 0.58 | 1.54 | 3.18 | 0.44 | 0.50 | 0.41 | 2.67 | 2.54 | 2.74 | 2.60 | 1.78 | 1.90 |
| | P | 1.74 | 2.76 | 6.13 | 4.92 | 3.61 | 2.27 | 1.12 | 4.23 | 3.85 | 3.25 | 2.91 | 5.51 | 2.00 | 4.13 | 3.29 | 2.18 | 3.56 | 1.61 | 3.83 | 3.59 | 0.92 | | | |

| Patient | | 1 | 2 | 3 | 4 | 5 | 6 | 7 | 8 | 9 | 10 | 11 | 12 | 13 | 14 | 15 | 16 | 17 | 18 | 19 | 20 | 21 | 22 | 23 | 24 |
|---|---|---|---|---|---|---|---|---|---|---|---|---|---|---|---|---|---|---|---|---|---|---|---|---|---|
| No. 4 | A | 0.92 | 1.17 | 3.04 | 3.06 | 1.23 | 1.42 | 1.48 | 2.25 | 1.28 | 1.29 | 1.31 | 2.82 | 1.44 | 1.83 | 1.16 | 1.34 | 1.99 | 1.43 | 1.35 | 1.22 | 1.02 | 5.85 | 1.84 | 5.02 |
| | B | 0.95 | 1.09 | 2.60 | 3.62 | 3.16 | 1.22 | 1.84 | 1.73 | 1.58 | 2.48 | 2.03 | 3.18 | 1.43 | 2.04 | 2.52 | 1.36 | 1.74 | 1.49 | 7.25 | 3.07 | 2.61 | 1.23 | 3.64 | 2.27 |
| | C | 1.13 | 2.09 | 5.07 | 5.50 | 2.52 | 1.46 | 1.68 | 2.20 | 5.49 | 1.69 | 1.90 | 2.23 | 1.43 | 2.11 | 3.08 | 1.22 | 1.65 | 1.33 | 2.76 | 2.32 | 2.45 | 2.33 | 2.39 | 3.25 |
| | D | 1.40 | 2.44 | 5.68 | 5.46 | 2.74 | 1.48 | 2.30 | 3.28 | 2.78 | 2.22 | 2.05 | 2.67 | 1.67 | 2.47 | 2.07 | 1.54 | 1.96 | 1.72 | 2.47 | 2.35 | 1.41 | 3.02 | 2.58 | 1.27 |
| | E | 0.89 | 1.68 | 4.37 | 4.82 | 2.10 | 1.47 | 1.11 | 2.29 | 2.59 | 1.92 | 1.56 | 2.06 | 1.42 | 1.53 | 2.43 | 1.30 | 1.77 | 0.83 | 2.22 | 2.10 | 1.66 | 1.27 | 2.65 | 2.15 |
| | F | 0.42 | 1.23 | 2.81 | 3.41 | 2.36 | 0.82 | 0.90 | 1.13 | 2.95 | 1.66 | 1.46 | 1.50 | 1.01 | 1.57 | 2.52 | 0.81 | 1.00 | 0.67 | 2.46 | 2.28 | 1.45 | 3.45 | 5.81 | 5.33 |
| | G | 1.63 | 1.79 | 5.01 | 4.84 | 1.80 | 2.98 | 2.18 | 2.32 | 1.71 | 1.60 | 1.69 | 3.75 | 3.36 | 2.07 | 1.73 | 2.36 | 2.66 | 2.36 | 1.62 | 1.81 | 2.24 | 1.33 | 1.99 | 3.47 |
| | H | 0.80 | 1.22 | 2.93 | 3.41 | 1.33 | 0.69 | 1.31 | 1.77 | 1.47 | 1.51 | 1.41 | 2.22 | 0.66 | 1.52 | 1.39 | 0.83 | 1.33 | 1.21 | 1.36 | 1.35 | 2.55 | 2.39 | 2.43 | 2.93 |
| | I | 1.05 | 1.54 | 3.59 | 3.81 | 1.11 | 1.09 | 1.79 | 2.07 | 1.25 | 1.06 | 1.18 | 2.69 | 1.25 | 1.72 | 1.22 | 1.25 | 1.80 | 1.71 | 1.22 | 1.06 | 1.25 | 2.69 | 2.29 | 1.54 |
| | J | 0.85 | 1.74 | 3.70 | 3.48 | 1.15 | 0.95 | 1.24 | 1.58 | 1.24 | 1.12 | 1.04 | 2.04 | 1.05 | 1.34 | 1.14 | 1.12 | 1.34 | 1.29 | 1.29 | 1.20 | 1.96 | 1.57 | 2.40 | 2.26 |
| | K | 0.66 | 1.15 | 3.08 | 3.26 | 1.52 | 1.17 | 0.87 | 1.41 | 2.69 | 1.25 | 1.04 | 1.47 | 1.11 | 1.08 | 1.55 | 1.04 | 1.08 | 1.09 | 2.02 | 1.79 | 1.09 | 2.21 | 2.86 | 2.76 |
| | L | 0.90 | 1.12 | 1.15 | 1.24 | 1.48 | 1.35 | 1.10 | 1.47 | 1.35 | 1.59 | 1.07 | 1.31 | 1.45 | 1.11 | 1.10 | 1.26 | 1.54 | 1.22 | 1.52 | 1.42 | 2.15 | 2.23 | 2.60 | 1.56 |
| | M | 0.74 | 2.53 | 5.34 | 5.26 | 2.13 | 0.93 | 1.67 | 1.67 | 3.32 | 2.18 | 2.09 | 2.15 | 0.71 | 2.84 | 2.38 | 1.02 | 1.23 | 1.17 | 2.26 | 2.26 | 1.27 | 1.07 | 0.91 | 1.97 |
| | N | 0.53 | 1.32 | 2.61 | 2.75 | 1.00 | 0.44 | 1.10 | 1.86 | 1.39 | 1.07 | 1.15 | 1.81 | 0.55 | 1.27 | 1.05 | 0.71 | 1.03 | 1.07 | 1.21 | 1.02 | 3.62 | 4.22 | 0.90 | 2.50 |
| | O | 0.49 | 1.53 | 5.44 | 8.01 | 2.50 | 0.64 | 1.24 | 1.40 | 3.61 | 2.63 | 1.35 | 1.79 | 1.04 | 2.28 | 3.17 | 0.84 | 1.05 | 0.77 | 3.00 | 2.83 | 3.38 | 2.95 | 2.18 | 2.06 |
| | P | 2.41 | 3.40 | 6.86 | 6.83 | 3.79 | 3.88 | 2.06 | 8.26 | 4.10 | 3.24 | 3.46 | 6.15 | 3.67 | 4.08 | 3.61 | 3.33 | 4.78 | 3.35 | 3.96 | 2.87 | 8.32 | | | |

| Patient | | 1 | 2 | 3 | 4 | 5 | 6 | 7 | 8 | 9 | 10 | 11 | 12 | 13 | 14 | 15 | 16 | 17 | 18 | 19 | 20 | 21 | 22 | 23 | 24 |
|---|---|---|---|---|---|---|---|---|---|---|---|---|---|---|---|---|---|---|---|---|---|---|---|---|---|
| No. 5 | A | 1.33 | 5.87 | 5.75 | 5.46 | 0.57 | 2.08 | 2.04 | 1.77 | 1.39 | 1.00 | 0.81 | 6.09 | 2.42 | 2.51 | 1.09 | 2.41 | 3.10 | 1.50 | 0.55 | 0.64 | 0.51 | 2.88 | 1.94 | 2.22 |
| | B | 2.14 | 2.79 | 10.76 | 8.94 | 5.13 | 1.91 | 5.51 | 4.12 | 3.73 | 3.45 | 2.67 | 3.90 | 1.33 | 3.06 | 4.43 | 1.38 | 2.26 | 1.54 | 4.42 | 3.21 | 1.81 | 0.38 | 0.99 | 0.85 |
| | C | 1.32 | 5.05 | 7.52 | 9.23 | 1.71 | 1.52 | 2.21 | 2.33 | 5.51 | 1.74 | 2.08 | 3.97 | 1.16 | 4.39 | 3.83 | 1.85 | 2.04 | 1.89 | 1.69 | 2.70 | 1.59 | 2.05 | 1.76 | 2.81 |
| | D | 0.75 | 3.33 | 3.95 | 2.96 | 2.55 | 0.87 | 1.65 | 3.76 | 2.28 | 2.52 | 1.78 | 2.64 | 1.21 | 1.99 | 1.45 | 1.19 | 1.64 | 1.39 | 2.88 | 2.19 | 0.76 | 2.50 | 1.33 | 0.92 |
| | E | 1.06 | 3.15 | 8.56 | 5.95 | 1.17 | 1.65 | 1.10 | 2.31 | 5.03 | 1.89 | 1.14 | 4.21 | 1.33 | 2.52 | 3.30 | 2.05 | 4.05 | 0.84 | 1.24 | 1.80 | 1.10 | 0.99 | 4.46 | 2.34 |
| | F | 0.41 | 3.10 | 4.06 | 3.44 | 1.77 | 0.88 | 0.97 | 1.20 | 5.18 | 1.83 | 1.01 | 2.25 | 0.65 | 2.69 | 4.04 | 0.63 | 0.96 | 0.50 | 2.56 | 2.01 | 1.02 | 9.67 | 5.41 | 5.71 |
| | G | 0.81 | 4.22 | 2.44 | 2.52 | 1.33 | 1.79 | 1.77 | 2.16 | 1.20 | 0.79 | 1.02 | 3.05 | 3.68 | 1.41 | 0.82 | 1.78 | 2.03 | 2.13 | 1.42 | 0.91 | 2.32 | 1.05 | 1.52 | 2.69 |
| | H | 1.10 | 5.56 | 6.13 | 7.37 | 0.79 | 1.11 | 1.64 | 2.34 | 1.81 | 1.11 | 1.04 | 3.11 | 1.16 | 2.12 | 1.56 | 1.26 | 1.98 | 1.82 | 0.79 | 0.94 | 2.34 | 1.78 | 1.43 | 3.28 |
| | I | 0.40 | 4.71 | 2.28 | 2.53 | 0.70 | 0.98 | 1.09 | 1.49 | 0.97 | 0.65 | 0.55 | 2.49 | 1.18 | 1.09 | 0.43 | 0.99 | 1.24 | 1.11 | 0.90 | 0.52 | 1.03 | 2.73 | 1.43 | 1.25 |
| | J | 0.40 | 7.11 | 2.18 | 2.09 | 0.55 | 0.62 | 0.52 | 0.82 | 0.64 | 0.50 | 0.35 | 1.38 | 0.72 | 0.63 | 0.45 | 0.50 | 0.88 | 0.84 | 0.68 | 0.30 | 1.78 | 1.47 | 2.56 | 1.76 |
| | K | 0.94 | 4.69 | 6.29 | 8.40 | 1.54 | 1.23 | 0.97 | 1.10 | 7.22 | 2.02 | 1.45 | 2.24 | 1.02 | 2.60 | 2.54 | 1.29 | 1.48 | 0.93 | 2.78 | 2.69 | 1.11 | 5.54 | 4.95 | 2.59 |
| | L | 0.31 | 1.06 | 0.55 | 0.58 | 0.29 | 0.58 | 0.32 | 0.62 | 1.10 | 0.46 | 0.24 | 0.57 | 0.69 | 0.41 | 0.62 | 0.51 | 0.77 | 0.33 | 0.54 | 0.29 | 0.75 | 3.09 | 4.07 | 0.52 |
| | M | 1.06 | 7.57 | 8.48 | 7.48 | 1.04 | 1.41 | 2.51 | 1.34 | 4.71 | 1.52 | 1.64 | 3.98 | 0.93 | 4.84 | 2.95 | 1.35 | 1.79 | 1.21 | 1.52 | 1.80 | 1.20 | 0.25 | 0.40 | 3.99 |
| | N | 0.74 | 8.37 | 3.76 | 3.72 | 0.31 | 0.80 | 1.42 | 1.78 | 1.57 | 0.49 | 0.43 | 2.92 | 0.84 | 1.49 | 0.80 | 0.93 | 1.57 | 1.29 | 0.56 | 0.33 | 8.27 | 2.39 | 0.54 | 5.27 |
| | O | 0.66 | 8.77 | 27.53 | 38.69 | 3.09 | 1.20 | 2.15 | 2.11 | 12.59 | 3.62 | 2.15 | 7.58 | 0.98 | 9.26 | 10.21 | 1.17 | 2.03 | 1.06 | 4.56 | 5.77 | 9.49 | 4.62 | 1.12 | 1.02 |
| | P | 4.20 | 21.36 | 12.15 | 14.41 | 3.16 | 7.32 | 4.02 | 10.24 | 5.43 | 3.01 | 2.55 | 12.58 | 11.54 | 10.27 | 4.88 | 8.10 | 12.38 | 6.94 | 5.82 | 3.20 | 2.80 | | | |

**Supplementary Figure S1. The results of primary screening with tissue lysates from five patients with high-grade serous carcinoma (HGSC).** The conversion rate (CVR) of 0.9 μM probes in the library after incubation with 0.05 mg/mL tissue lysate for 3 h. All assays were performed at 37°C in 10 μL total volume of phosphate-buffered saline (pH 7.4) containing 100 mg/L $CaCl_2$ and $MgCl_2 \cdot 6H_2O$, and 0.5 % (v/v) DMSO as a cosolvent. Fluorescence intensity at 0 h or 3 h was calculated as an average of three measurements. The tumor-to-normal (T/N) ratio was calculated by dividing the conversion rate (CVR) of each probe in tumor lysate by that in paired normal lysate. Excitation/emission wavelengths were 485/535 nm. The conversion rate (CVR) was calculated as follows.

$$\text{Conversion rate} = \frac{(\text{F.I. of probe at 3 h} - \text{F.I. of probe at 0 h})}{(\text{F.I. of HMRG at 3 h} - \text{F.I. of HMRG at 0 h})}$$

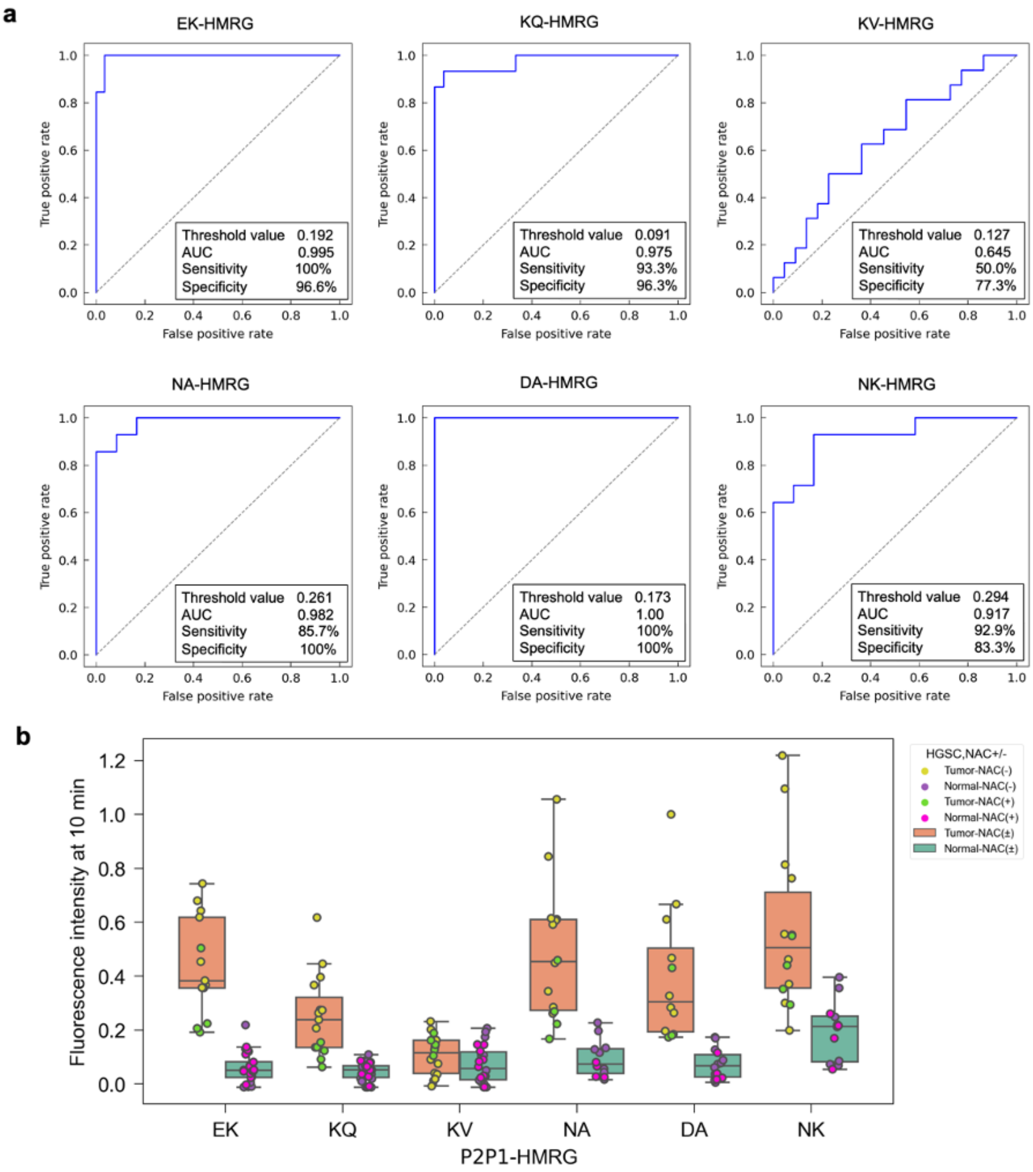


**Supplementary Figure S2. Comparison of fluorescence increase in tumor and adjacent normal peritoneal tissues at 10 min.** (**a**) Threshold value, sensitivity, specificity and AUC of each probe were evaluated from the receiver operating characteristic. Tumor (N = 13) and Normal (N = 29) tissues were examined with EK-HMRG. Tumor (N = 15) and Normal (N = 27) tissues were examined with KQ-HMRG. Tumor (N = 16) and Normal (N = 22) tissues were examined with KQ-HMRG. Tumor (N = 14) and Normal (N = 12) tissues were examined with NA-HMRG. Tumor (N = 12) and Normal (N = 14) tissues were examined with DA-HMRG. Tumor (N = 14) and Normal (N = 12) tissues were examined with NK-HMRG. (**b**) Comprehensive analysis of fluorescence intensity in tumor and adjacent normal peritoneal tissues using 6 fluorescent probes (N = 12−16 for Tumor, N = 12−29 for Normal). Fluorescence increase represents the increase at 10 min after addition of fluorescent probes. Pink, purple, green, and yellow dots represent fluorescence increases in tumor tissues without neoadjuvant chemotherapy (NAC), tumor tissues with NAC, normal tissues without NAC, and tumor tissues with NAC, respectively.

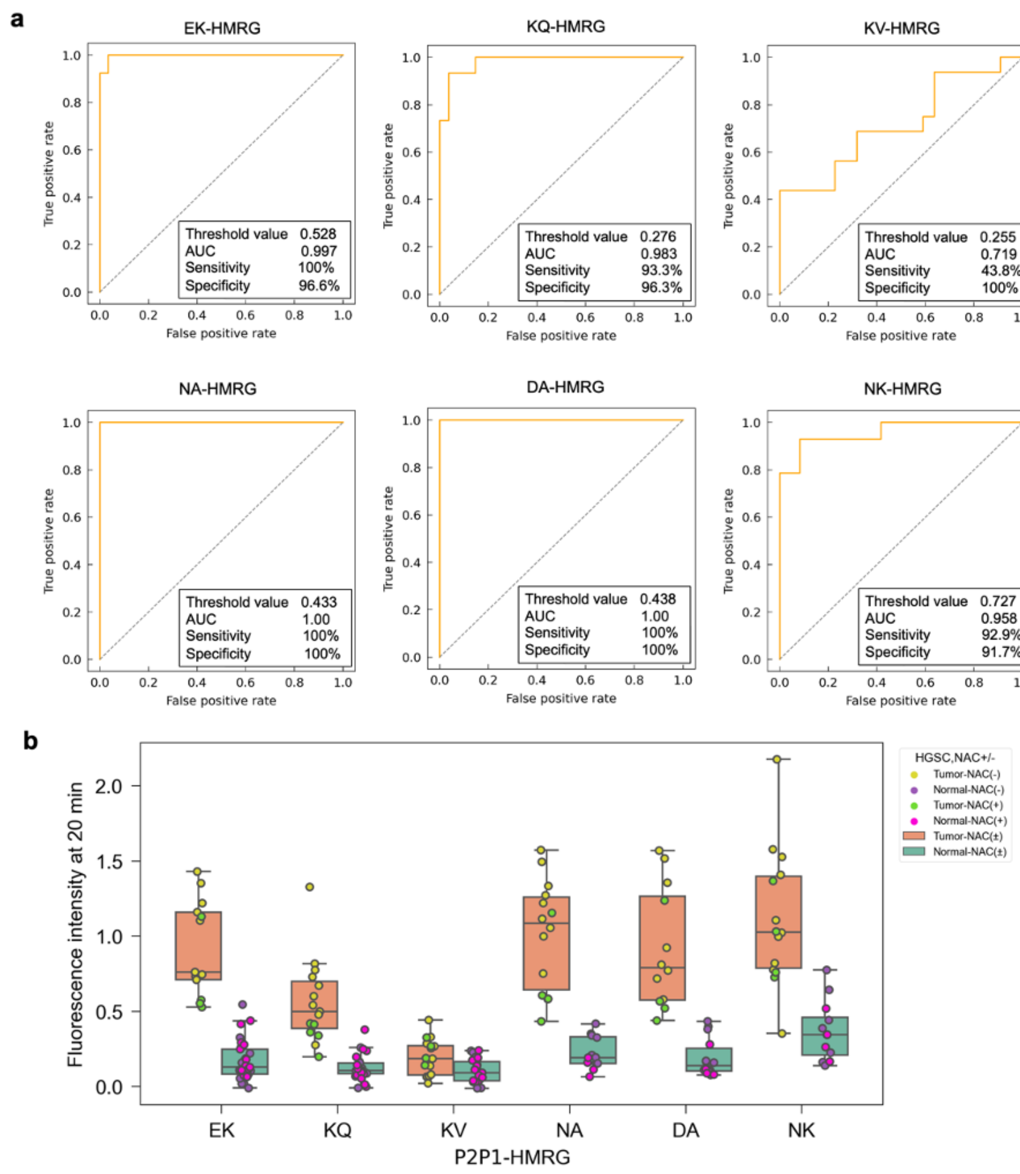


**Supplementary Figure S3. Comparison of fluorescence increases in tumor and adjacent normal peritoneal tissues at 20 min after probe application.** (**a**) Threshold value, sensitivity, specificity and AUC of each probe were evaluated from the receiver operating characteristic. Tumor (N = 13) and Normal (N = 29) tissues were examined with EK-HMRG. Tumor (N = 15) and Normal (N = 27) tissues were examined with KQ-HMRG. Tumor (N = 16) and Normal (N = 22) tissues were examined with KQ-HMRG. Tumor (N = 14) and Normal (N = 12) tissues were examined with NA-HMRG. Tumor (N = 12) and Normal (N = 14) tissues were examined with DA-HMRG. Tumor (N = 14) and Normal (N = 12) tissues were examined with NK-HMRG. (**b**) Comprehensive analysis of fluorescence intensity in tumor and adjacent normal peritoneal tissues using 6 fluorescent probes (N = 12−16 for Tumor, N = 12−29 for Normal). Fluorescence increase represent increase at 20 min after addition of fluorescent probes. Pink, purple, green, and yellow dots represent fluorescence increases in tumor tissues without neoadjuvant chemotherapy (NAC), tumor tissues with NAC, normal tissues without NAC, and tumor tissues with NAC, respectively.

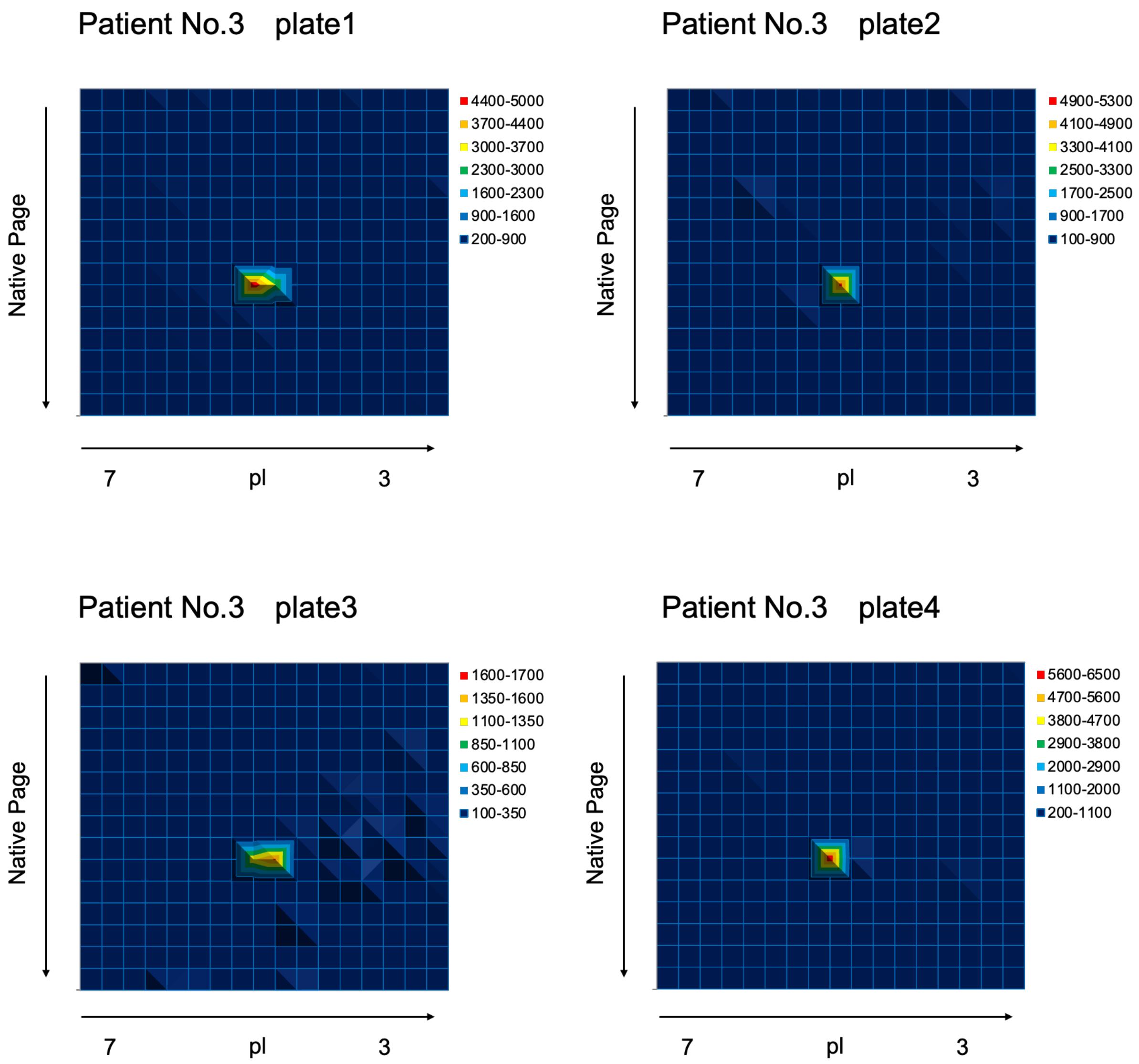


**Supplementary Figure S4. DEG assay using EK-HMRG with peritoneal dissemination lysate.** The assays were carried out as reported.(1) A single fluorescent spot was observed on 2D gels and puromycin-sensitive aminopeptidase (PSA) was identified by peptide mass fingerprinting analysis. The lysate sample was evaluated on four gels. Gels were incubated with the probe at 37°C for 1 h. Proteins detected by peptide mass fingerprinting analysis are summarized in Tables S4, and the clinical information of Patient No. 3, from whom the analyzed lysate was derived, is provided in Table S1. [EK-HMRG] = 70 μM.

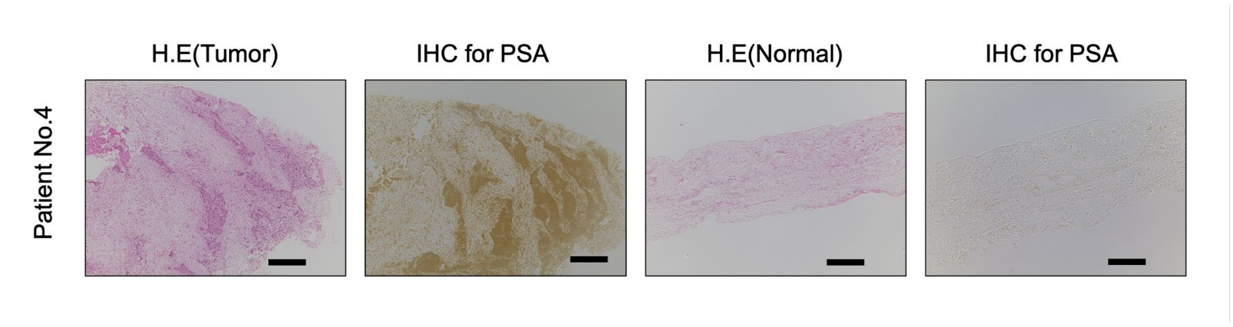


**Supplementary Figure S5. HE and IHC staining of ovarian cancer peritoneal dissemination tissue from Patient No.4.** PSA was overexpressed in tumor tissues compared to normal tissues. Scale bar, 200 μm.

.

#### Supplementary Tables

**Supplementary Table S1. Patient characteristics of 20 cases with high-grade serous carcinoma.** Nine patients had received neoadjuvant chemotherapy (NAC) prior to surgery. Age values represent age at surgery. Samples from Patients No.16, 17, 18, and 20 were subjected to fluorescence imaging in a fresh condition.

| Number | Age (years) | FIGO stage | NAC | Specimen |
|---|---|---|---|---|
| 1 | 66 | IIIC | No | peritoneum |
| 2 | 55 | IIIB | No | peritoneum, omentum |
| 3 | 59 | IIIC | No | peritoneum |
| 4 | 50 | IIIC | No | peritoneum |
| 5 | 38 | IIIB | No | peritoneum |
| 6 | 62 | IIIC | Yes | peritoneum |
| 7 | 49 | IIIC | Yes | omentum |
| 8 | 42 | IIIC | Yes | peritoneum, omentum |
| 9 | 35 | IIIA | No | peritoneum |
| 10 | 67 | IVA | Yes | peritoneum |
| 11 | 55 | IIIB | No | peritoneum |
| 12 | 69 | IIIC | Yes | peritoneum |
| 13 | 77 | IIIC | Yes | peritoneum |
| 14 | 52 | IVB | No | peritoneum |
| 15 | 47 | IIIC | Yes | omentum |
| 16 | 48 | IIIC | No | omentum |
| 17 | 85 | IIIC | No | omentum |
| 18 | 48 | IIIC | Yes | peritoneum |
| 19 | 68 | IIIA | No | peritoneum |
| 20 | 76 | IIIC | Yes | peritoneum |

FIGO, International Federation of Gynecology and Obstetrics.

**Supplementary Table S2. Abbreviations of amino acids.**

| Abbreviation | Amino acid |
|---|---|
| A | Alanine |
| C | Cysteine |
| D | Aspartic Acid |
| E | Glutamic Acid |
| F | Phenylalanine |
| G | Glycine |
| H | Histidine |
| I | Isoleucine |
| K | Lysine |
| L | Leucine |
| M | Methionine |
| N | Asparagine |
| P | Proline |
| Q | Glutamine |
| R | Arginine |
| S | Serine |
| T | Threonine |
| U | Valine |
| W | Tryptophan |
| Y | Tyrosine |
| gGlu | gamma-glutamyl |

**Supplementary Table S3. List of detected proteins in DEG assay of peritoneal dissemination specimen.** 190 proteins were identified by peptide mass fingerprinting analysis. Puromycin sensitive aminopeptidase is highlighted in yellow.

| | Identified Proteins (190) | Molecular Weight | Protein Identification Probability |
|---|---|---|---|
| 1 | Puromycin-sensitive aminopeptidase OS=Homo sapiens OX=9606 GN=NPEPPS PE=1 SV=2 | 103 kDa | 100% |
| 2 | Albumin OS=Homo sapiens OX=9606 GN=ALB PE=1 SV=2 | 69 kDa | 100% |
| 3 | Vitamin D-binding protein OS=Homo sapiens OX=9606 GN=GC PE=1 SV=2 | 53 kDa | 100% |
| 4 | Cluster of Actin, cytoplasmic 1 OS=Homo sapiens OX=9606 GN=ACTB PE=1 SV=1 (P60709) | 42 kDa | 100% |
| 5 | Keratin, type II cytoskeletal 1 OS=Homo sapiens OX=9606 GN=KRT1 PE=1 SV=6 | 66 kDa | 100% |
| 6 | Cluster of Heat shock protein HSP 90-beta OS=Homo sapiens OX=9606 GN=HSP90AB1 PE=1 SV=4 (P08238) | 83 kDa | 100% |
| 7 | Endoplasmic reticulum chaperone BiP OS=Homo sapiens OX=9606 GN=HSPA5 PE=1 SV=2 | 72 kDa | 100% |
| 8 | Importin subunit beta-1 OS=Homo sapiens OX=9606 GN=KPNB1 PE=1 SV=2 | 97 kDa | 100% |
| 9 | Keratin, type I cytoskeletal 10 OS=Homo sapiens OX=9606 GN=KRT10 PE=1 SV=6 | 59 kDa | 100% |
| 10 | Keratin, type I cytoskeletal 9 OS=Homo sapiens OX=9606 GN=KRT9 PE=1 SV=3 | 62 kDa | 100% |
| 11 | Ubiquitin-like modifier-activating enzyme 1 OS=Homo sapiens OX=9606 GN=UBA1 PE=1 SV=3 | 118 kDa | 100% |
| 12 | Lumican OS=Homo sapiens OX=9606 GN=LUM PE=1 SV=2 | 38 kDa | 100% |
| 13 | Keratin, type II cytoskeletal 2 epidermal OS=Homo sapiens OX=9606 GN=KRT2 PE=1 SV=2 | 65 kDa | 100% |
| 14 | Neutral alpha-glucosidase AB OS=Homo sapiens OX=9606 GN=GANAB PE=1 SV=3 | 107 kDa | 100% |
| 15 | Endoplasmin OS=Homo sapiens OX=9606 GN=HSP90B1 PE=1 SV=1 | 92 kDa | 100% |
| 16 | Isocitrate dehydrogenase [NADP] cytoplasmic OS=Homo sapiens OX=9606 GN=IDH1 PE=1 SV=2 | 47 kDa | 100% |
| 17 | Heat shock cognate 71 kDa protein OS=Homo sapiens OX=9606 GN=HSPA8 PE=1 SV=1 | 71 kDa | 100% |
| 18 | Protein disulfide-isomerase A4 OS=Homo sapiens OX=9606 GN=PDIA4 PE=1 SV=2 | 73 kDa | 100% |
| 19 | SUMO-activating enzyme subunit 2 OS=Homo sapiens OX=9606 GN=UBA2 PE=1 SV=2 | 71 kDa | 100% |
| 20 | Plastin-2 OS=Homo sapiens OX=9606 GN=LCP1 PE=1 SV=6 | 70 kDa | 100% |
| 21 | Antithrombin-III OS=Homo sapiens OX=9606 GN=SERPINC1 PE=1 SV=1 | 53 kDa | 100% |
| 22 | Cluster of L-lactate dehydrogenase B chain OS=Homo sapiens OX=9606 GN=LDHB PE=1 SV=2 (P07195) | 37 kDa | 100% |
| 23 | Cluster of cAMP-dependent protein kinase type II-beta regulatory subunit OS=Homo sapiens OX=9606 GN=PRKAR2B PE=1 SV=3 (P31323) | 46 kDa | 100% |
| 24 | Beta-galactosidase OS=Homo sapiens OX=9606 GN=GLB1 PE=1 SV=2 | 76 kDa | 100% |
| 25 | Serine/threonine-protein phosphatase 2A 65 kDa regulatory subunit A alpha isoform OS=Homo sapiens OX=9606 GN=PPP2R1A PE=1 SV=4 | 65 kDa | 100% |
| 26 | Nucleolin OS=Homo sapiens OX=9606 GN=NCL PE=1 SV=3 | 77 kDa | 100% |
| 27 | SHC1_HUMAN-DECOY | ? | 84% |
| 28 | Cluster of Heat shock 70 kDa protein 1A OS=Homo sapiens OX=9606 GN=HSPA1A PE=1 SV=1 (P0DMV8) | 70 kDa | 100% |
| 29 | Cluster of cAMP-dependent protein kinase catalytic subunit alpha OS=Homo sapiens OX=9606 GN=PRKACA PE=1 SV=2 (P17612) | 41 kDa | 100% |
| 30 | Cytoplasmic aconitate hydratase OS=Homo sapiens OX=9606 GN=ACO1 PE=1 SV=3 | 98 kDa | 100% |
| 31 | Hexokinase-3 OS=Homo sapiens OX=9606 GN=HK3 PE=1 SV=2 | 99 kDa | 100% |
| 32 | Inter-alpha-trypsin inhibitor heavy chain H4 OS=Homo sapiens OX=9606 GN=ITIH4 PE=1 SV=4 | 103 kDa | 100% |
| 33 | Kininogen-1 OS=Homo sapiens OX=9606 GN=KNG1 PE=1 SV=2 | 72 kDa | 100% |
| 34 | NEDD8-activating enzyme E1 catalytic subunit OS=Homo sapiens OX=9606 GN=UBA3 PE=1 SV=2 | 52 kDa | 100% |

| | | | |
|---|---|---|---|
| 35 | Leukotriene A-4 hydrolase OS=Homo sapiens OX=9606 GN=LTA4H PE=1 SV=2 | 69 kDa | 100% |
| 36 | Nuclear autoantigenic sperm protein OS=Homo sapiens OX=9606 GN=NASP PE=1 SV=2 | 85 kDa | 100% |
| 37 | NEDD8-activating enzyme E1 regulatory subunit OS=Homo sapiens OX=9606 GN=NAE1 PE=1 SV=1 | 60 kDa | 100% |
| 38 | Transportin-2 OS=Homo sapiens OX=9606 GN=TNPO2 PE=1 SV=3 | 101 kDa | 100% |
| 39 | Hemopexin OS=Homo sapiens OX=9606 GN=HPX PE=1 SV=2 | 52 kDa | 100% |
| 40 | Xaa-Pro dipeptidase OS=Homo sapiens OX=9606 GN=PEPD PE=1 SV=3 | 55 kDa | 100% |
| 41 | Keratin, type II cytoskeletal 5 OS=Homo sapiens OX=9606 GN=KRT5 PE=1 SV=3 | 62 kDa | 100% |
| 42 | Interleukin enhancer-binding factor 2 OS=Homo sapiens OX=9606 GN=ILF2 PE=1 SV=2 | 43 kDa | 100% |
| 43 | Afamin OS=Homo sapiens OX=9606 GN=AFM PE=1 SV=1 | 69 kDa | 100% |
| 44 | Cluster of Hemoglobin subunit beta OS=Homo sapiens OX=9606 GN=HBB PE=1 SV=2 (P68871) | 16 kDa | 100% |
| 45 | Transportin-1 OS=Homo sapiens OX=9606 GN=TNPO1 PE=1 SV=2 | 102 kDa | 100% |
| 46 | Keratin, type I cytoskeletal 14 OS=Homo sapiens OX=9606 GN=KRT14 PE=1 SV=4 | 52 kDa | 100% |
| 47 | Glucosidase 2 subunit beta OS=Homo sapiens OX=9606 GN=PRKCSH PE=1 SV=2 | 59 kDa | 100% |
| 48 | Leucine-rich PPR motif-containing protein, mitochondrial OS=Homo sapiens OX=9606 GN=LRPPRC PE=1 SV=3 | 158 kDa | 100% |
| 49 | Cell division control protein 42 homolog OS=Homo sapiens OX=9606 GN=CDC42 PE=1 SV=2 | 21 kDa | 100% |
| 50 | Heterogeneous nuclear ribonucleoprotein U OS=Homo sapiens OX=9606 GN=HNRNPU PE=1 SV=6 | 91 kDa | 100% |
| 51 | tRNA (guanine(37)-N1)-methyltransferase OS=Homo sapiens OX=9606 GN=TRMT5 PE=1 SV=2 | 58 kDa | 100% |
| 52 | THUMP domain-containing protein 1 OS=Homo sapiens OX=9606 GN=THUMPD1 PE=1 SV=2 | 39 kDa | 100% |
| 53 | Cluster of Heterogeneous nuclear ribonucleoprotein R OS=Homo sapiens OX=9606 GN=HNRNPR PE=1 SV=1 (O43390) | 71 kDa | 100% |
| 54 | Lupus La protein OS=Homo sapiens OX=9606 GN=SSB PE=1 SV=2 | 47 kDa | 100% |
| 55 | ATP synthase subunit beta, mitochondrial OS=Homo sapiens OX=9606 GN=ATP5F1B PE=1 SV=3 | 57 kDa | 100% |
| 56 | Mimecan OS=Homo sapiens OX=9606 GN=OGN PE=1 SV=1 | 34 kDa | 100% |
| 57 | Protein SET OS=Homo sapiens OX=9606 GN=SET PE=1 SV=3 | 33 kDa | 100% |
| 58 | Interleukin enhancer-binding factor 3 OS=Homo sapiens OX=9606 GN=ILF3 PE=1 SV=3 | 95 kDa | 100% |
| 59 | FAS-associated factor 1 OS=Homo sapiens OX=9606 GN=FAF1 PE=1 SV=2 | 74 kDa | 100% |
| 60 | Prothrombin OS=Homo sapiens OX=9606 GN=F2 PE=1 SV=2 | 70 kDa | 100% |
| 61 | Alpha-1B-glycoprotein OS=Homo sapiens OX=9606 GN=A1BG PE=1 SV=4 | 54 kDa | 100% |
| 62 | Heparin cofactor 2 OS=Homo sapiens OX=9606 GN=SERPIND1 PE=1 SV=3 | 57 kDa | 100% |
| 63 | Adenylosuccinate synthetase isozyme 2 OS=Homo sapiens OX=9606 GN=ADSS2 PE=1 SV=3 | 50 kDa | 100% |
| 64 | 5-phosphohydroxy-L-lysine phospho-lyase OS=Homo sapiens OX=9606 GN=PHYKPL PE=1 SV=1 | 50 kDa | 100% |
| 65 | 5'-3' exoribonuclease 2 OS=Homo sapiens OX=9606 GN=XRN2 PE=1 SV=1 | 109 kDa | 100% |
| 66 | Citrate synthase, mitochondrial OS=Homo sapiens OX=9606 GN=CS PE=1 SV=2 | 52 kDa | 100% |
| 67 | 60 kDa heat shock protein, mitochondrial OS=Homo sapiens OX=9606 GN=HSPD1 PE=1 SV=2 | 61 kDa | 100% |
| 68 | Malate dehydrogenase, cytoplasmic OS=Homo sapiens OX=9606 GN=MDH1 PE=1 SV=4 | 36 kDa | 100% |
| 69 | Hemoglobin subunit alpha OS=Homo sapiens OX=9606 GN=HBA1 PE=1 SV=2 | 15 kDa | 100% |
| 70 | Rho GTPase-activating protein 1 OS=Homo sapiens OX=9606 GN=ARHGAP1 PE=1 SV=1 | 50 kDa | 100% |
| 71 | Immunity-related GTPase family Q protein OS=Homo sapiens OX=9606 GN=IRGQ PE=1 SV=1 | 63 kDa | 100% |
| 72 | Echinoderm microtubule-associated protein-like 4 OS=Homo sapiens OX=9606 GN=EML4 PE=1 SV=3 | 109 kDa | 100% |
| 73 | Tubulin beta chain OS=Homo sapiens OX=9606 GN=TUBB PE=1 SV=2 | 50 kDa | 100% |

| | | | |
|---|---|---|---|
| 74 | UV excision repair protein RAD23 homolog B OS=Homo sapiens OX=9606 GN=RAD23B PE=1 SV=1 | 43 kDa | 100% |
| 75 | Peptidyl-prolyl cis-trans isomerase FKBP4 OS=Homo sapiens OX=9606 GN=FKBP4 PE=1 SV=3 | 52 kDa | 100% |
| 76 | ATP-dependent RNA helicase A OS=Homo sapiens OX=9606 GN=DHX9 PE=1 SV=4 | 141 kDa | 100% |
| 77 | Hornerin OS=Homo sapiens OX=9606 GN=HRNR PE=1 SV=2 | 282 kDa | 100% |
| 78 | Glycerol-3-phosphate phosphatase OS=Homo sapiens OX=9606 GN=PGP PE=1 SV=1 | 34 kDa | 100% |
| 79 | Chloride intracellular channel protein 1 OS=Homo sapiens OX=9606 GN=CLIC1 PE=1 SV=4 | 27 kDa | 100% |
| 80 | Alpha-1-antitrypsin OS=Homo sapiens OX=9606 GN=SERPINA1 PE=1 SV=3 | 47 kDa | 100% |
| 81 | Complement component C9 OS=Homo sapiens OX=9606 GN=C9 PE=1 SV=2 | 63 kDa | 100% |
| 82 | cAMP-dependent protein kinase type I-alpha regulatory subunit OS=Homo sapiens OX=9606 GN=PRKAR1A PE=1 SV=1 | 43 kDa | 100% |
| 83 | Replication protein A 70 kDa DNA-binding subunit OS=Homo sapiens OX=9606 GN=RPA1 PE=1 SV=2 | 68 kDa | 100% |
| 84 | Ubiquitin-protein ligase E3A OS=Homo sapiens OX=9606 GN=UBE3A PE=1 SV=4 | 101 kDa | 100% |
| 85 | NAD kinase 2, mitochondrial OS=Homo sapiens OX=9606 GN=NADK2 PE=1 SV=2 | 49 kDa | 100% |
| 86 | Enoyl-[acyl-carrier-protein] reductase, mitochondrial OS=Homo sapiens OX=9606 GN=MECR PE=1 SV=2 | 40 kDa | 100% |
| 87 | Histidine-rich glycoprotein OS=Homo sapiens OX=9606 GN=HRG PE=1 SV=1 | 60 kDa | 100% |
| 88 | 60S acidic ribosomal protein P0 OS=Homo sapiens OX=9606 GN=RPLP0 PE=1 SV=1 | 34 kDa | 100% |
| 89 | Glutathione S-transferase P OS=Homo sapiens OX=9606 GN=GSTP1 PE=1 SV=2 | 23 kDa | 100% |
| 90 | Complement C1s subcomponent OS=Homo sapiens OX=9606 GN=C1S PE=1 SV=1 | 77 kDa | 100% |
| 91 | Hsc70-interacting protein OS=Homo sapiens OX=9606 GN=ST13 PE=1 SV=2 | 41 kDa | 100% |
| 92 | Serine/threonine-protein phosphatase 2A catalytic subunit beta isoform OS=Homo sapiens OX=9606 GN=PPP2CB PE=1 SV=1 | 36 kDa | 100% |
| 93 | Tubulin alpha-1B chain OS=Homo sapiens OX=9606 GN=TUBA1B PE=1 SV=1 | 50 kDa | 100% |
| 94 | Heterogeneous nuclear ribonucleoprotein D0 OS=Homo sapiens OX=9606 GN=HNRNPD PE=1 SV=1 | 38 kDa | 100% |
| 95 | Thioredoxin domain-containing protein 5 OS=Homo sapiens OX=9606 GN=TXNDC5 PE=1 SV=2 | 48 kDa | 100% |
| 96 | Nucleosome assembly protein 1-like 4 OS=Homo sapiens OX=9606 GN=NAP1L4 PE=1 SV=1 | 43 kDa | 100% |
| 97 | Lambda-crystallin homolog OS=Homo sapiens OX=9606 GN=CRYL1 PE=1 SV=3 | 35 kDa | 100% |
| 98 | Cysteine sulfinic acid decarboxylase OS=Homo sapiens OX=9606 GN=CSAD PE=1 SV=2 | 55 kDa | 100% |
| 99 | Clusterin OS=Homo sapiens OX=9606 GN=CLU PE=1 SV=1 | 52 kDa | 100% |
| 100 | Inter-alpha-trypsin inhibitor heavy chain H2 OS=Homo sapiens OX=9606 GN=ITIH2 PE=1 SV=2 | 106 kDa | 100% |
| 101 | m7GpppX diphosphatase OS=Homo sapiens OX=9606 GN=DCPS PE=1 SV=2 | 39 kDa | 100% |
| 102 | Cellular retinoic acid-binding protein 2 OS=Homo sapiens OX=9606 GN=CRABP2 PE=1 SV=2 | 16 kDa | 100% |
| 103 | Alpha-2-antiplasmin OS=Homo sapiens OX=9606 GN=SERPINF2 PE=1 SV=3 | 55 kDa | 100% |
| 104 | Ribonuclease inhibitor OS=Homo sapiens OX=9606 GN=RNH1 PE=1 SV=2 | 50 kDa | 100% |
| 105 | Serine/threonine-protein phosphatase 2B catalytic subunit beta isoform OS=Homo sapiens OX=9606 GN=PPP3CB PE=1 SV=2 | 59 kDa | 100% |
| 106 | Alpha-N-acetylgalactosaminidase OS=Homo sapiens OX=9606 GN=NAGA PE=1 SV=2 | 47 kDa | 100% |
| 107 | Glutathione synthetase OS=Homo sapiens OX=9606 GN=GSS PE=1 SV=1 | 52 kDa | 100% |
| 108 | Serine/threonine-protein phosphatase PP1-beta catalytic subunit OS=Homo sapiens OX=9606 GN=PPP1CB PE=1 SV=3 | 37 kDa | 100% |
| 109 | Fibromodulin OS=Homo sapiens OX=9606 GN=FMOD PE=1 SV=2 | 43 kDa | 100% |
| 110 | Ras GTPase-activating-like protein IQGAP2 OS=Homo sapiens OX=9606 GN=IQGAP2 PE=1 SV=4 | 181 kDa | 100% |
| 111 | Junction plakoglobin OS=Homo sapiens OX=9606 GN=JUP PE=1 SV=3 | 82 kDa | 100% |

| | | | |
|---|---|---|---|
| 112 | CCA tRNA nucleotidyltransferase 1, mitochondrial OS=Homo sapiens OX=9606 GN=TRNT1 PE=1 SV=2 | 50 kDa | 100% |
| 113 | EGF-containing fibulin-like extracellular matrix protein 1 OS=Homo sapiens OX=9606 GN=EFEMP1 PE=1 SV=2 | 55 kDa | 100% |
| 114 | Phosphoenolpyruvate carboxykinase [GTP], mitochondrial OS=Homo sapiens OX=9606 GN=PCK2 PE=1 SV=4 | 71 kDa | 100% |
| 115 | Alanine--tRNA ligase, cytoplasmic OS=Homo sapiens OX=9606 GN=AARS1 PE=1 SV=2 | 107 kDa | 100% |
| 116 | Ran GTPase-activating protein 1 OS=Homo sapiens OX=9606 GN=RANGAP1 PE=1 SV=1 | 64 kDa | 100% |
| 117 | Developmentally-regulated GTP-binding protein 2 OS=Homo sapiens OX=9606 GN=DRG2 PE=1 SV=1 | 41 kDa | 100% |
| 118 | Bisphosphoglycerate mutase OS=Homo sapiens OX=9606 GN=BPGM PE=1 SV=2 | 30 kDa | 100% |
| 119 | Inter-alpha-trypsin inhibitor heavy chain H3 OS=Homo sapiens OX=9606 GN=ITIH3 PE=1 SV=2 | 100 kDa | 100% |
| 120 | Malate dehydrogenase, mitochondrial OS=Homo sapiens OX=9606 GN=MDH2 PE=1 SV=3 | 36 kDa | 100% |
| 121 | Aflatoxin B1 aldehyde reductase member 2 OS=Homo sapiens OX=9606 GN=AKR7A2 PE=1 SV=3 | 40 kDa | 100% |
| 122 | Golgi reassembly-stacking protein 2 OS=Homo sapiens OX=9606 GN=GORASP2 PE=1 SV=3 | 47 kDa | 100% |
| 123 | Nucleosome assembly protein 1-like 1 OS=Homo sapiens OX=9606 GN=NAP1L1 PE=1 SV=1 | 45 kDa | 100% |
| 124 | Myosin light chain kinase, smooth muscle OS=Homo sapiens OX=9606 GN=MYLK PE=1 SV=4 | 211 kDa | 100% |
| 125 | 60S ribosomal protein L5 OS=Homo sapiens OX=9606 GN=RPL5 PE=1 SV=3 | 34 kDa | 100% |
| 126 | Calpain-2 catalytic subunit OS=Homo sapiens OX=9606 GN=CAPN2 PE=1 SV=6 | 80 kDa | 100% |
| 127 | General transcription factor II-I OS=Homo sapiens OX=9606 GN=GTF2I PE=1 SV=2 | 112 kDa | 100% |
| 128 | mRNA cap guanine-N7 methyltransferase OS=Homo sapiens OX=9606 GN=RNMT PE=1 SV=1 | 55 kDa | 100% |
| 129 | Eukaryotic translation initiation factor 2 subunit 1 OS=Homo sapiens OX=9606 GN=EIF2S1 PE=1 SV=3 | 36 kDa | 99% |
| 130 | Alpha-1-antichymotrypsin OS=Homo sapiens OX=9606 GN=SERPINA3 PE=1 SV=2 | 48 kDa | 100% |
| 131 | Fumarylacetoacetase OS=Homo sapiens OX=9606 GN=FAH PE=1 SV=2 | 46 kDa | 100% |
| 132 | Ras GTPase-activating-like protein IQGAP1 OS=Homo sapiens OX=9606 GN=IQGAP1 PE=1 SV=1 | 189 kDa | 100% |
| 133 | Heterogeneous nuclear ribonucleoprotein K OS=Homo sapiens OX=9606 GN=HNRNPK PE=1 SV=1 | 51 kDa | 100% |
| 134 | Filaggrin-2 OS=Homo sapiens OX=9606 GN=FLG2 PE=1 SV=1 | 248 kDa | 100% |
| 135 | Protein DDI1 homolog 2 OS=Homo sapiens OX=9606 GN=DDI2 PE=1 SV=1 | 45 kDa | 100% |
| 136 | Xaa-Pro aminopeptidase 1 OS=Homo sapiens OX=9606 GN=XPNPEP1 PE=1 SV=3 | 70 kDa | 100% |
| 137 | Vimentin OS=Homo sapiens OX=9606 GN=VIM PE=1 SV=4 | 54 kDa | 100% |
| 138 | F-box only protein 21 OS=Homo sapiens OX=9606 GN=FBXO21 PE=2 SV=2 | 72 kDa | 100% |
| 139 | Replication protein A 32 kDa subunit OS=Homo sapiens OX=9606 GN=RPA2 PE=1 SV=1 | 29 kDa | 100% |
| 140 | Heterogeneous nuclear ribonucleoprotein A/B OS=Homo sapiens OX=9606 GN=HNRNPAB PE=1 SV=2 | 36 kDa | 100% |
| 141 | F-actin-capping protein subunit beta OS=Homo sapiens OX=9606 GN=CAPZB PE=1 SV=5 | 31 kDa | 100% |
| 142 | Ceruloplasmin OS=Homo sapiens OX=9606 GN=CP PE=1 SV=1 | 122 kDa | 100% |
| 143 | Heat shock 70 kDa protein 4 OS=Homo sapiens OX=9606 GN=HSPA4 PE=1 SV=4 | 94 kDa | 100% |
| 144 | Alpha-2-HS-glycoprotein OS=Homo sapiens OX=9606 GN=AHSG PE=1 SV=2 | 39 kDa | 100% |
| 145 | Keratinocyte proline-rich protein OS=Homo sapiens OX=9606 GN=KPRP PE=1 SV=1 | 64 kDa | 100% |
| 146 | Caspase-14 OS=Homo sapiens OX=9606 GN=CASP14 PE=1 SV=2 | 28 kDa | 100% |
| 147 | 14-3-3 protein theta OS=Homo sapiens OX=9606 GN=YWHAQ PE=1 SV=1 | 28 kDa | 100% |
| 148 | Serine protease 1 OS=Homo sapiens OX=9606 GN=PRSS1 PE=1 SV=1 | 27 kDa | 95% |
| 149 | Uncharacterized protein C17orf80 OS=Homo sapiens OX=9606 GN=C17orf80 PE=1 SV=2 | 67 kDa | 87% |

| | | | |
|---|---|---|---|
| 150 | K1C9_HUMAN-DECOY | ? | 95% |
| 151 | Na(+)/H(+) exchange regulatory cofactor NHE-RF1 OS=Homo sapiens OX=9606 GN=SLC9A3R1 PE=1 SV=4 | 39 kDa | 95% |
| 152 | Histone acetyltransferase type B catalytic subunit OS=Homo sapiens OX=9606 GN=HAT1 PE=1 SV=2 | 50 kDa | 95% |
| 153 | Eukaryotic translation initiation factor 3 subunit D OS=Homo sapiens OX=9606 GN=EIF3D PE=1 SV=1 | 64 kDa | 95% |
| 154 | Tumor protein D54 OS=Homo sapiens OX=9606 GN=TPD52L2 PE=1 SV=2 | 22 kDa | 95% |
| 155 | EGF-containing fibulin-like extracellular matrix protein 2 OS=Homo sapiens OX=9606 GN=EFEMP2 PE=1 SV=3 | 49 kDa | 95% |
| 156 | Angiotensinogen OS=Homo sapiens OX=9606 GN=AGT PE=1 SV=3 | 53 kDa | 95% |
| 157 | Complement C3 OS=Homo sapiens OX=9606 GN=C3 PE=1 SV=2 | 187 kDa | 95% |
| 158 | 40S ribosomal protein SA OS=Homo sapiens OX=9606 GN=RPSA PE=1 SV=4 | 33 kDa | 95% |
| 159 | Signal recognition particle 19 kDa protein OS=Homo sapiens OX=9606 GN=SRP19 PE=1 SV=3 | 16 kDa | 95% |
| 160 | Complement C4-A OS=Homo sapiens OX=9606 GN=C4A PE=1 SV=2 | 193 kDa | 95% |
| 161 | Polyubiquitin-B OS=Homo sapiens OX=9606 GN=UBB PE=1 SV=1 | 26 kDa | 95% |
| 162 | Lysosome-associated membrane glycoprotein 1 OS=Homo sapiens OX=9606 GN=LAMP1 PE=1 SV=3 | 45 kDa | 95% |
| 163 | Desmoplakin OS=Homo sapiens OX=9606 GN=DSP PE=1 SV=3 | 332 kDa | 95% |
| 164 | Biglycan OS=Homo sapiens OX=9606 GN=BGN PE=1 SV=2 | 42 kDa | 95% |
| 165 | Heterogeneous nuclear ribonucleoprotein H OS=Homo sapiens OX=9606 GN=HNRNPH1 PE=1 SV=4 | 49 kDa | 95% |
| 166 | Peroxiredoxin-2 OS=Homo sapiens OX=9606 GN=PRDX2 PE=1 SV=5 | 22 kDa | 95% |
| 167 | Trypsin-3 OS=Homo sapiens OX=9606 GN=PRSS3 PE=1 SV=2 | 33 kDa | 95% |
| 168 | Profilin-2 OS=Homo sapiens OX=9606 GN=PFN2 PE=1 SV=3 | 15 kDa | 95% |
| 169 | Y-box-binding protein 1 OS=Homo sapiens OX=9606 GN=YBX1 PE=1 SV=3 | 36 kDa | 95% |
| 170 | Nucleobindin-2 OS=Homo sapiens OX=9606 GN=NUCB2 PE=1 SV=3 | 50 kDa | 95% |
| 171 | Dermcidin OS=Homo sapiens OX=9606 GN=DCD PE=1 SV=2 | 11 kDa | 95% |
| 172 | Transcription intermediary factor 1-beta OS=Homo sapiens OX=9606 GN=TRIM28 PE=1 SV=5 | 89 kDa | 95% |
| 173 | GDP-L-fucose synthase OS=Homo sapiens OX=9606 GN=GFUS PE=1 SV=1 | 36 kDa | 95% |
| 174 | Bleomycin hydrolase OS=Homo sapiens OX=9606 GN=BLMH PE=1 SV=1 | 53 kDa | 95% |
| 175 | Prostaglandin reductase 1 OS=Homo sapiens OX=9606 GN=PTGR1 PE=1 SV=2 | 36 kDa | 95% |
| 176 | FERM and PDZ domain-containing protein 2 OS=Homo sapiens OX=9606 GN=FRMPD2 PE=1 SV=3 | 144 kDa | 95% |
| 177 | 5'-3' exonuclease PLD3 OS=Homo sapiens OX=9606 GN=PLD3 PE=1 SV=1 | 55 kDa | 95% |
| 178 | Putative E3 ubiquitin-protein ligase UBR7 OS=Homo sapiens OX=9606 GN=UBR7 PE=1 SV=2 | 48 kDa | 95% |
| 179 | NHL repeat-containing protein 2 OS=Homo sapiens OX=9606 GN=NHLRC2 PE=1 SV=1 | 79 kDa | 95% |
| 180 | Protein phosphatase Slingshot homolog 3 OS=Homo sapiens OX=9606 GN=SSH3 PE=1 SV=2 | 73 kDa | 95% |
| 181 | Perilipin-4 OS=Homo sapiens OX=9606 GN=PLIN4 PE=1 SV=2 | 134 kDa | 95% |
| 182 | Opioid growth factor receptor OS=Homo sapiens OX=9606 GN=OGFR PE=1 SV=3 | 73 kDa | 95% |
| 183 | Importin subunit alpha-5 OS=Homo sapiens OX=9606 GN=KPNA1 PE=1 SV=3 | 60 kDa | 94% |
| 184 | Nardilysin OS=Homo sapiens OX=9606 GN=NRDC PE=1 SV=3 | 132 kDa | 84% |
| 185 | Rho-related GTP-binding protein RhoC OS=Homo sapiens OX=9606 GN=RHOC PE=1 SV=1 | 22 kDa | 84% |
| 186 | Skin-specific protein 32 OS=Homo sapiens OX=9606 GN=XP32 PE=1 SV=1 | 26 kDa | 84% |
| 187 | Band 4.1-like protein 2 OS=Homo sapiens OX=9606 GN=EPB41L2 PE=1 SV=1 | 113 kDa | 57% |
| 188 | LCAP_HUMAN-DECOY | ? | 32% |
| 189 | Maleylacetoacetate isomerase OS=Homo sapiens OX=9606 GN=GSTZ1 PE=1 SV=3 | 24 kDa | 32% |
| 190 | Voltage-dependent calcium channel gamma-6 subunit OS=Homo sapiens OX=9606 GN=CACNG6 PE=1 SV=1 | 28 kDa | 32% |
